# The Synaptic Architecture of Layer 5 Thick Tufted Excitatory Neurons in the Visual Cortex of Mice

**DOI:** 10.1101/2023.10.18.562531

**Authors:** Agnes L. Bodor, Casey M Schneider-Mizell, Chi Zhang, Leila Elabbady, Alex Mallen, Andi Bergeson, Derrick Brittain, JoAnn Buchanan, Daniel J. Bumbarger, Rachel Dalley, Clare Gamlin, Emily Joyce, Daniel Kapner, Sam Kinn, Gayathri Mahalingam, Sharmishtaa Seshamani, Shelby Suckow, Marc Takeno, Russel Torres, Wenjing Yin, J. Alexander Bae, Manuel A. Castro, Sven Dorkenwald, Akhilesh Halageri, Zhen Jia, Chris Jordan, Nico Kemnitz, Kisuk Lee, Kai Li, Ran Lu, Thomas Macrina, Eric Mitchell, Shanka Subhra Mondal, Shang Mu, Barak Nehoran, Sergiy Popovych, William Silversmith, Nicholas L. Turner, Szi-chieh Yu, William Wong, Jingpeng Wu, Brendan Celii, Luke Campagnola, Stephanie C Seeman, Tim Jarsky, Naixin Ren, Anton Arkhipov, The MICrONS Consortium, Jacob Reimer, H Sebastian Seung, R. Clay Reid, Forrest Collman, Nuno Maçarico da Costa

**Affiliations:** Allen Institute for Brain Science; Computer Science Department, Princeton University; Electrical and Computer Engineering Department, Princeton University; Princeton Neuroscience Institute, Princeton University; Department of Neuroscience, Baylor College of Medicine; Center for Neuroscience and Artificial Intelligence, Baylor College of Medicine; Department of Electrical and Computer Engineering, Rice University; Brain & Cognitive Sciences Department, Massachusetts Institute of Technology; Institute for Bioinformatics and Medical Informatics, University Tübingen; Institute for Computer Science, University Göttingen; International Max Planck Research School for Intelligent Systems, University Tübingen; Institute of Molecular Biology and Biotechnology, Foundation for Research and Technology Hellas, Heraklion, Greece; School of Applied and Engineering Physics, Cornell University

## Abstract

The neocortex is one of the most critical structures that makes us human, and it is involved in a variety of cognitive functions from perception to sensory integration and motor control. Composed of repeated modules, or microcircuits, the neocortex relies on distinct cell types as its fundamental building blocks. Despite significant progress in characterizing these cell types ^1–5^, an understanding of the complete synaptic partners associated with individual excitatory cell types remain elusive.

Here, we investigate the connectivity of arguably the most well recognized and studied excitatory neuron in the neocortex: the thick tufted layer 5 pyramidal cell ^6–10^ also known as extra telencephalic (ET) ^11^ neurons. Although the synaptic interactions of ET neurons have been extensively explored, a comprehensive characterization of their local connectivity remains lacking. To address this knowledge gap, we leveraged a 1 mm^3^ electron microscopic (EM) dataset.

We found that ET neurons primarily establish connections with inhibitory cells in their immediate vicinity. However, when they extend their axons to other cortical regions, they tend to connect more with excitatory cells. We also find that the inhibitory cells targeted by ET neurons are a specific group of cell types, and they preferentially inhibit ET cells. Finally, we observed that the most common excitatory targets of ET neurons are layer 5 IT neurons and layer 6 pyramidal cells, whereas synapses with other ET neurons are not as common.

These findings challenge current views of the connectivity of ET neurons and suggest a circuit design that involves local competition among ET neurons and collaboration with other types of excitatory cells. Our results also highlight a specific circuit pattern where a subclass of excitatory cells forms a network with specific inhibitory cell types, offering a framework for exploring the connectivity of other types of excitatory cells.

## Main text

Cell types have long been a focal point in connectivity studies. Cajal integrated this concept with the principle of dynamic polarization, significantly advancing the mapping of neuronal connections in various regions of the nervous system. However, Cajal faced a challenge in the neocortex, due to the “apparent disorder of the cerebral jungle, so different from the regularity and symmetry of the spinal cord and of the cerebellum”. Cajal was looking for a neuronal chain that bridged input to output and while he found it in other structures, such a chain is not obvious in the neocortex due to the strong recurrence and high overlap between dendrites and axons of most cell types. To overcome this, measuring synaptic connectivity became essential, and as synapses are visible with the electron microscope (EM) ^12–14^, this technology offers a way to reconstruct the connectivity and fine morphology of different cell types ^15–18^. Following the pioneering work of Winfried Denk and Heinz Horstmann ^19^ to scale the capacity of electron microscopy, several increasingly larger EM datasets have been generated across many species ^20–29^, including imaging of a cubic millimeter of mammalian brains at synapse resolution ^26,29^. In the mouse, this volume is sufficient to include substantial portions of the local arbor of neurons in the primary visual cortex as well as some of their inter-areal projections to neighboring higher visual regions.

Leveraging this technology, we explore the initial stages of the “neuronal chain” of layer 5 thick tufted cells, one of the most well studied and recognizable excitatory neurons of the neocortex ^6–11,30,31^. Their apical dendrite, with an exuberant tuft in the superficial layer of the cortex, receives thousands of synapses across most cortical layers integrating inputs from other cell types. These cells are also referred to as extratelencephalic (ET) ^32^ projection neurons (which we will use throughout this manuscript) and serve as the main output cell type of the neocortex, projecting subcortically and providing driver input to higher order thalamic areas ^6,7,33–36^. In some cases, ET axons bypass the thalamus and directly reach other subcortical structures like the pons^8^ or spinal cord ^37,38^, directly commanding body parts and therefore are also known as pyramidal-tract (PT) neurons.

While the extensive subcortical projections of the ET neurons play a pivotal role as a major output of the cortical circuit, they also form local connections. Understanding these local connections provides valuable insights into how the cortex implements its output function. For example, layer 5 contains a diverse population of inhibitory cells, particularly Basket and Martinotti cell ^39–41^ subclasses, both represented by numerous cell types ^42,43^. Although the interaction between ETs and this inhibitory network ^39,44–46^ has been extensively studied, including reciprocal connectivity with ETs ^39^, the extent of the targeting onto inhibitory versus excitatory cell types and the involvement of specific inhibitory types remains unclear. Resolving these fundamental questions about the connectivity with inhibitory cells has important consequences for the theories about how cortical circuits operate ^47–49^.

The local excitatory connectivity of layer 5 ET neurons has been primarily examined in terms of their recurrent connections with other ET neurons ^6,30,31,50–54^. Overall, ET to ET connections are rare but ET neurons that share similar subcortical projections have a higher probability of targeting each other ^31,54^ forming interconnected motifs ^52,53^. The ET neurons also share layer 5 with a diversity of other excitatory cells that form local and intra-telencephalic (IT) projections^55^. The connectivity with the layer 5 IT neurons has been described as rare or nonexistent ^30,31,39,54^. These results suggest that the main output neuron of the cortex does not share a copy of its output with the other cell types of the microcircuit, as is expected from some computational models ^56,57^. One possibility is that ET neurons do not target other layer 5 and layer 6 cell types that inhabit the vicinity of their axons. Alternatively, the ET neurons do communicate the results of their computation to other excitatory cell types, but these connections have eluded us because we lacked the ability to make an unbiased and comprehensive census of their connectivity. The MICrONs dataset is perfectly poised to create such a comprehensive census of connectivity as we can analyze most of the chemical synapses and potential targets of individual neurons within a single cortical area and across cortical areas. Due to the extensive body of research dedicated to ET neurons, they serve as an ideal subject to construct the first comprehensive map of the local output connections of a cortical cell type. The results from our study offer new insights into the connectivity pattern of ET neurons suggesting a different view of the involvement of this cell type in the cortical microcircuits, while still remaining consistent with previous data. The approach described in this manuscript can also now be scaled across the other excitatory cell types and across the cortical microcircuit.

### Large scale Electron Microscopy across multiple visual areas

The EM dataset used in this study ^26^ includes primary and higher order visual areas of a male mouse cortex at postnatal day 87 (p87) (see Fig1, a). This fully segmented EM dataset ^58^ is publicly available for access at www.microns-explorer.org. Within this dataset, there are approximately 200,000 cells, with around 120,000 of them being neurons based on nuclear and somatic features. Automated synapse detection methods, we have identified approximately 523 million synapses within the dataset.

The neurons within the dataset have been classified ^59–62^ into inhibitory and excitatory cell classes, along with several subclasses (Fig1, c and d, Extended Fig 1 and Extended Fig 2). This classification process initially involved the assessment of morphological features of a subset of neurons by human experts and then transitioned to dataset-wide classification using quantitative morphological and connectivity properties ^59^. The classifications used here follow the results of a companion paper ^59^. Briefly, we distinguished four inhibitory subclasses based on their output connectivity, inspired by classical neuroanatomy: Perisomatic targeting cells (PeriTC) primarily target the soma and proximal dendrites of target excitatory neurons, distal dendrite targeting cells (DistTC) primarily target distal basal and/or apical dendrites, inhibitory targeting cells (InhTC) primarily target other inhibitory neurons, and sparsely targeting cells (SparTC) synapse more diffusely onto target cells. These subclasses aligned approximately with classical anatomical and molecular classifications, although they might not always exhibit a one-to-one correspondence. For example, the PeriTC subclass encompasses basket cells from multiple molecular subclasses, such as both PV and CCK. Similarly, the DistTC subclass includes both Martinotti and non-Martinotti putative SST cells. Excitatory subclasses were predicted across the dataset using somatic and nuclear features, trained on a data-driven clustering of extensive quantitative measurements of dendritic properties and then consolidated into traditional layer groups (Layer ⅔ pyramidal, Layer 4 pyramidal, L5 pyramidal, and Layer 6 pyramidal). Layer 5 excitatory neurons were further split into three subclasses based on likely long range projection targets: thick tufted ET neurons that project subcortically, intratelencephalic (IT) neurons, also known as thin or slender tufted cells, that project to other regions of cortex and near projecting (NP) neurons, characterized by their long apical dendrites and sparse spines.^55^ For cells used in analysis, cell classes were further confirmed either using an independent spine-detection based classification model^61^ (where possible) or by expert evaluation (where not possible or in case of disagreement). Visual evaluation of the dendrites of 1,638 excitatory targets of L5-ET cells showed that our strategy was consistent with established morphological classifications (Extended Fig 11).

**Figure 1.**
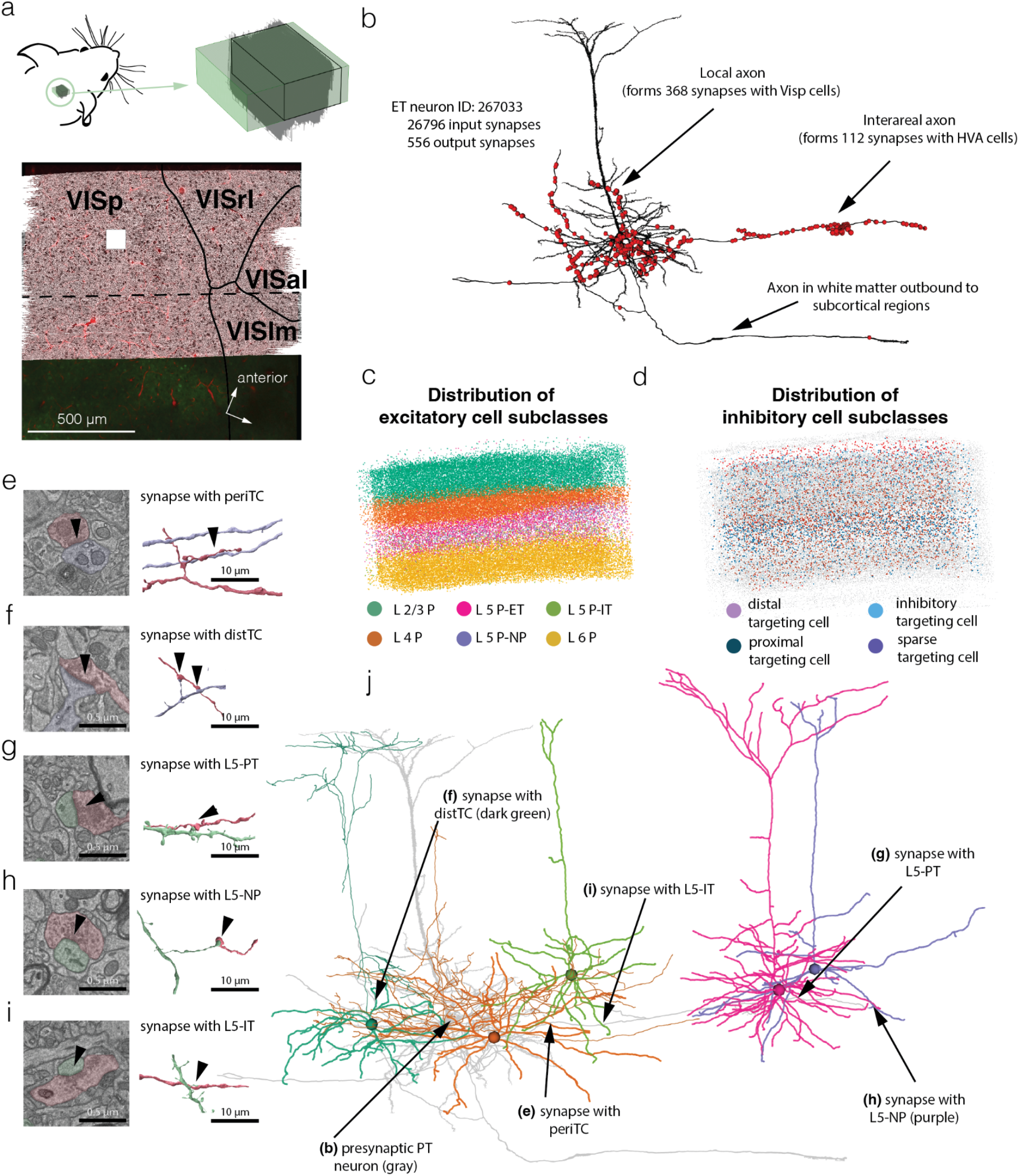
MICrONS ^26^ Electron Microscopy Dataset. **a.** left: mouse head with the location of the dataset, right: schematic view of the dataset (green). Below on the map, the top-down view of the mouse visual cortex and its area borders are presented in correspondence to the MICrONs mm3 dataset. VISp-primary visual cortex, VISrl-VISal and VISlm are higher order visual cortices. White square labels the top down view position of the column ^59^ analyzed in Figure 2. Arrow shows the anterior/posterior and medial/lateral axis. **b.** Example of the L5 thick tufted cell (black), outgoing synapses (n=556) labeled by red dots. **c.** Distribution of automated labeled excitatory cell subclasses across the volume ^60^. Each cell is represented with a colored dot. **d.** Distribution of automated labeled inhibitory cell subclasses across the volume ^60^. Similarly to **(c)** each cell is represented with a colored dot. **e-i.** Example synapses formed by the ET neuron in **(b)** with different postsynaptic cell types in layer 5. On each panel an electron micrograph (left) and the 3D representation of the same synapse (right) are visible between **(e)** an ET and a perisomatic targeting cell (periTC), **(f)** and ET and distal targeting cell (distTC), **(g)** between two ET cells, **(h)** and ET and a layer 5 near projecting (NP) cell, **(i)** and ET and a layer 5 inter-telencephalic projecting cell. On the micrograph the arrow shows the synapse, and the light red overlay represents the presynaptic site of the synapse while the postsynaptic element is colored in purple (inhibitory) or green (excitatory). Arrow and colors on the 3D reconstruction next to the synapse follow the same color code. Note, that in case of inhibitory targets there are multiple synapses between elements, while in case of excitatory targets the single synapse is the typical. **j.** 3D reconstruction of the same ET cell (light gray) as in b and together with colored reconstruction of 5 different postsynaptic targets. The location of the synapses shown in **(e-i)** are also indicated by arrows.

**Figure 2.**
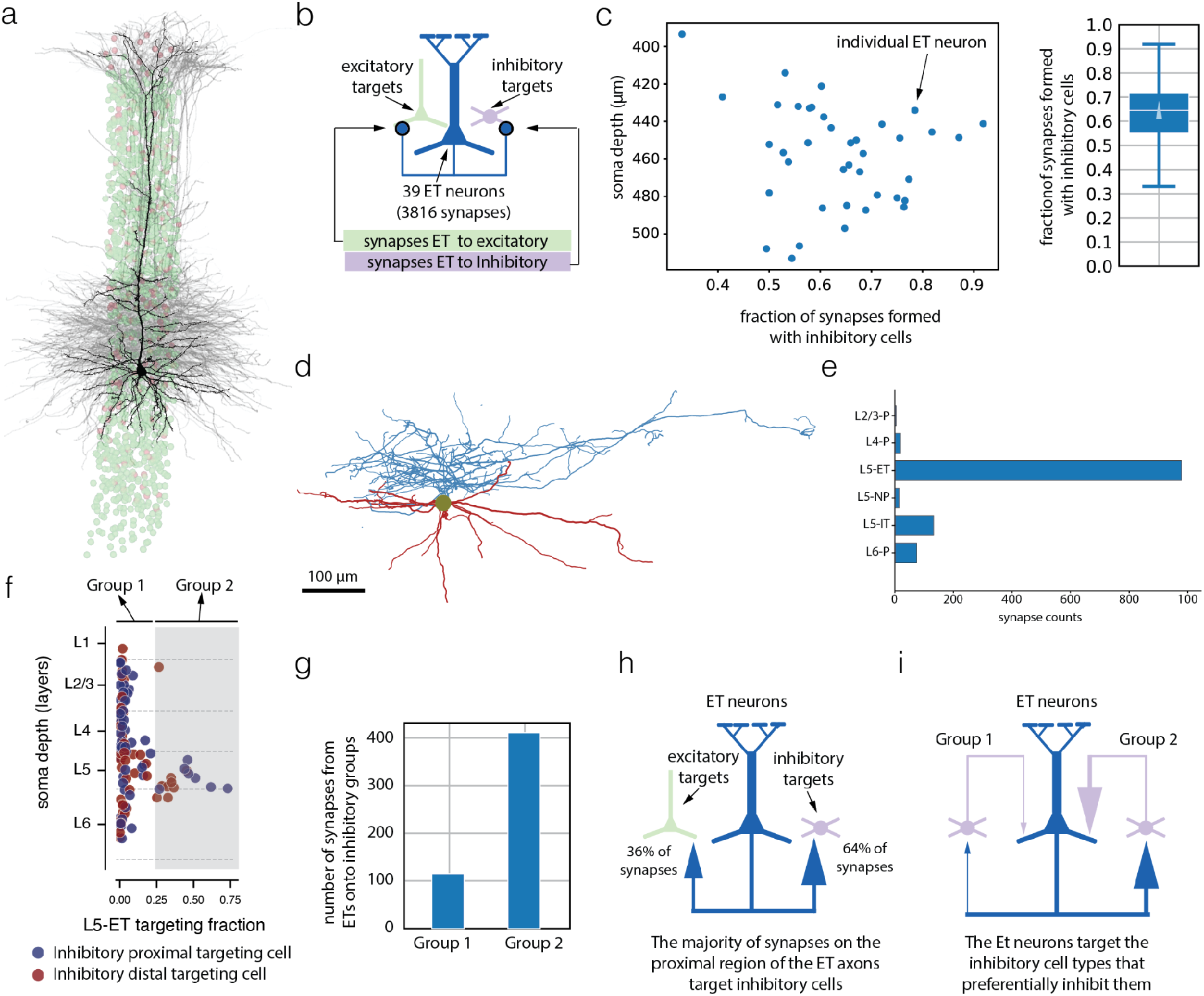
Layer 5 ET targeting of inhibitory cells. **a.** All the ET cells (n = 39, gray and black) in layer 5 reconstructed from a column in the primary visual cortex of the mouse. Dots represent nuclei of all inhibitory cells (purple) and excitatory cells (green) in the same column as the ET cells. **b.** scatter plot (right) showing the fraction of synapses formed by each individual ET cell (n=39) onto postsynaptic inhibitory cells and boxplot (left) of the same data (triangle indicates the mean fraction of ET synapses formed by inhibitory target cells). **d.** Shows an example of a layer 5 inhibitory perisomatic targeting cell (axon displayed in blue and dendrite in red) that preferentially targets ET neurons ^59^ as shown ‘e’. **e.** Histogram showing the number of synapses formed by the cell in ‘d’ with different excitatory subclasses. **f.** shows two subclasses of inhibitory cells within the column defined in (a) arranged by depth (y axis) and by fraction of synapses formed with ET cells (x axis). Perisomatic targeting cells are represented by blue dots while red dots represent the distal targeting interneurons. These inhibitory cells were further separated into 2 groups indicated by the white/gray background color. The white background inhibitory cells (Group 1) have up to 25 % of their output synapses formed with ET cells. The gray background inhibitory cells (Group 2) on the right form more than 25 % of their synapses with ET cells. **g.** Histogram showing that the ET neurons from the column in (a) form the majority of synapses with Group 2 inhibitory cells. **h.** Summary diagram showing that the majority of local synapses formed by the ET neurons are with Inhibitory cells. **i.** Summary diagram showing that the majority of local ET synapses are formed with inhibitory cells that recurrently strongly inhibit the ET neurons. The main input and output inhibitory cells of ET neurons are the same group of cells.

While most dendrites were well-segmented, the automated segmentation required additional manual proofreading to produce axons with good enough quality for analysis.For the specific focus of this study, our attention was directed towards layer 5 ET neurons (as depicted in Fig 1.b) and their postsynaptic partners (Fig 1.e-j). The axons of the ET neurons that spanned across the VISp, VISrl and VISal portion of the dataset were cleaned and extended, enabling us to examine and characterize synaptic connectivity both within the cortical region and across different cortical regions (as shown in Fig 1.b). In our analysis, we examined a total of 7586 output synapses originating from a subset of 47 ET neurons. This comprehensive examination of synaptic connections allowed us to generate detailed cell type connectivity maps, shedding light on the intricate network of synaptic interactions and revealing previously undescribed connectivity patterns.

### Layer 5 ET neurons form the majority of their local synapses with inhibitory neurons

As with any other neuron, the firing activity of ET cells has a complex interplay with the surrounding inhibitory network. Previous studies in different layers and regions have demonstrated a strong bias of excitatory neurons towards targeting inhibitory neurons in the proximal region of the axon ^47,48,63^. Thus, our study sought to first determine if a similar pronounced bias existed within ET neurons.

To address this question, we focused our analysis on ET neurons located within a well-characterized column ^59^ in the center of the V1 region of the dataset. This column, measuring 100 um x 100 um and reaching from layer 1 to white matter, encompassed all cortical layers (Fig 2.a-b). All the ET neurons in the column have been identified based on their distinctive morphological features ^59^. The axonal projections of all the ET neurons in the column (n = 39) were then proofread to ensure the presence of at least 100 synapses, which were identified using automated synapse detection methods ^58,64^. Following the removal of any synapse segmentation errors, a total of 3816 synapses were deemed suitable for analysis. To minimize potential bias in the analysis of these synapses, we examined all postsynaptic targets with orphan dendrites and spines (dendrites not attached to a neuronal soma) and, whenever possible, merged them back to their respective neurons. After this proofreading process, only 229 synapses remained detached or orphaned, of which 116 were individual spines and 113 longer stretches of dendrites, where the soma was partial or outside of the dataset. We then proceeded to classify the postsynaptic targets of ET neurons as either excitatory or inhibitory.

On average, approximately two-thirds of the synapses formed by ET neurons were found to connect with inhibitory cells, with the majority of individual presynaptic ET cells establishing more than 50% of their synapses with inhibitory neurons (Fig 2.c-d).

### Layer 5 ET forms specific sub-circuits with inhibitory cells that target ET neurons

Layer 5 inhibitory cells exhibit highly selective patterns of inhibition towards specific types of excitatory cells ^59,65,66^, and these precise connectivity patterns extend across various inhibitory cell types ^59^. Building upon this knowledge and considering the strong targeting of inhibition by ET neurons shown in the previous section (Fig 2.d), we asked whether the ET neurons form output synapses with inhibitory neurons that provide feedback inhibition to the ET neurons (i.e., ET neurons competing with each other) or if the ET neurons primarily target inhibitory cells that inhibit other types of excitatory cells. To address this question at a coarse level, we divided the inhibitory neurons within the column described in Fig 2 into two groups: Group 1 comprised neurons that strongly target ET cells, defined as having more than 25% of their outputs onto ET cells, and a Group 2 with a lower proportion of ET outputs. (Fig 2.e). The threshold of 0.25 was determined based on the ET output fraction of neurons in layer 5 and our previous work on classifying groups of inhibitory cells within this column ^59^ (see methods, Extended Fig 1). Figure 2.f illustrates an example of an inhibitory cell specifically targeting ET cells. Group 2 cells received 4 times more ET synapses in total than Group1 cells (Fig 2.g, 411 synapses onto group 2 cells versus 115 onto Group 1 cells). Hence, our results reveal that ET neurons predominantly target Group 2 cells, their primary source of inhibitory input (Extended Fig 3 uses the more detailed motif classification of Schneider-Mizell et. al ^59^ with similar conclusions).

**Figure 3.**
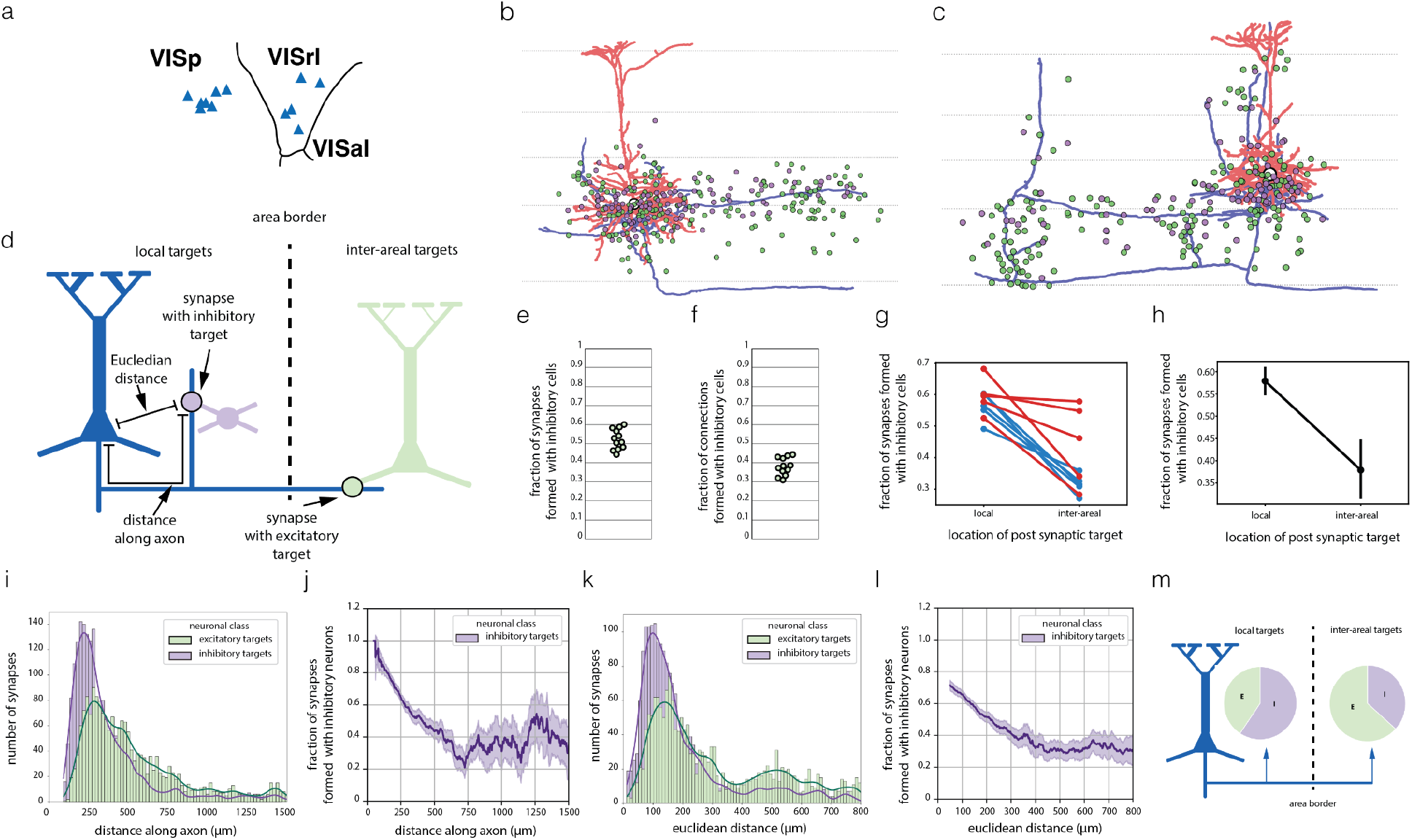
Local and inter-areal projections of ET neurons. **a.** Top down map of EM volume with the area borders and location of ET neurons (blue triangles) with clean and fully extended axons. **b. and c.** Examples of individual neurons, one in V1 (panel b) and the other in a higher order visual area (panel c). Axons are shown in blue, the soma-dendritic region is red, and dots show the locations of the somas of postsynaptic neurons targeted by the ET neuron. Inhibitory postsynaptic cells are shown as purple dots and the postsynaptic excitatory cells are shown as green dots. Horizontal lines label the borders between the cortical layers. **d.** Cartoon summarizing the key steps of the analyses. For each synapse on the axon, we determined the class of the postsynaptic neuron, distinguishing between excitatory and inhibitory cells. Additionally, we identified the location of the postsynaptic target, distinguishing between local and inter-areal connections to other brain regions within the EM volume. Moreover, we measured the euclidean distance between the synapse and the ET soma, as well as the path length distance along the axon from the synapse to the soma. **e.** Fraction of synapses formed by the ET neurons with inhibitory targets across the whole extent of the axons **f.** Fraction of the postsynaptic neurons of ET axons that are inhibitory. **g.** Fraction of synapses formed by the ET neurons with inhibitory cells, categorized into local and inter-areal targets. The data is displayed for each individual cell, with color coding indicating whether the soma of the ET neuron is located in V1 (blue) or HVA (red). **h.** Mean fraction of synapses formed by the ET neurons with inhibitory cells, categorized into local and inter-areal targets. **i.** Histogram of distance dependence of synapses with inhibitory and excitatory cells along the axon path length. **j.** Distance dependence of the ratio of inhibitory v excitatory synapses along the axon pathlength. **k.** Histogram of distance dependence of synapses with inhibitory and excitatory cells measured as the euclidean distance between the ET neuron and the synapse. **l.** Distance dependence of the ratio of inhibitory to excitatory synapses using euclidean distance between the synapse and the soma. **m.** Summary diagram showing that while locally the ET neurons preferentially target Inhibitory cell, their inter-areal projection forms the majority of their synapses with excitatory cells.

As a control and to show that this result is not just restricted to the ET neurons and inhibitory targets in the column, we also included additional perisomatic targeting cells that were postsynaptic to the 12 fully extended ET neurons shown in Fig 3. In this expanded set all the inhibitory cells fell into group 2 (Extended Fig 4).

**Fig. 4.**
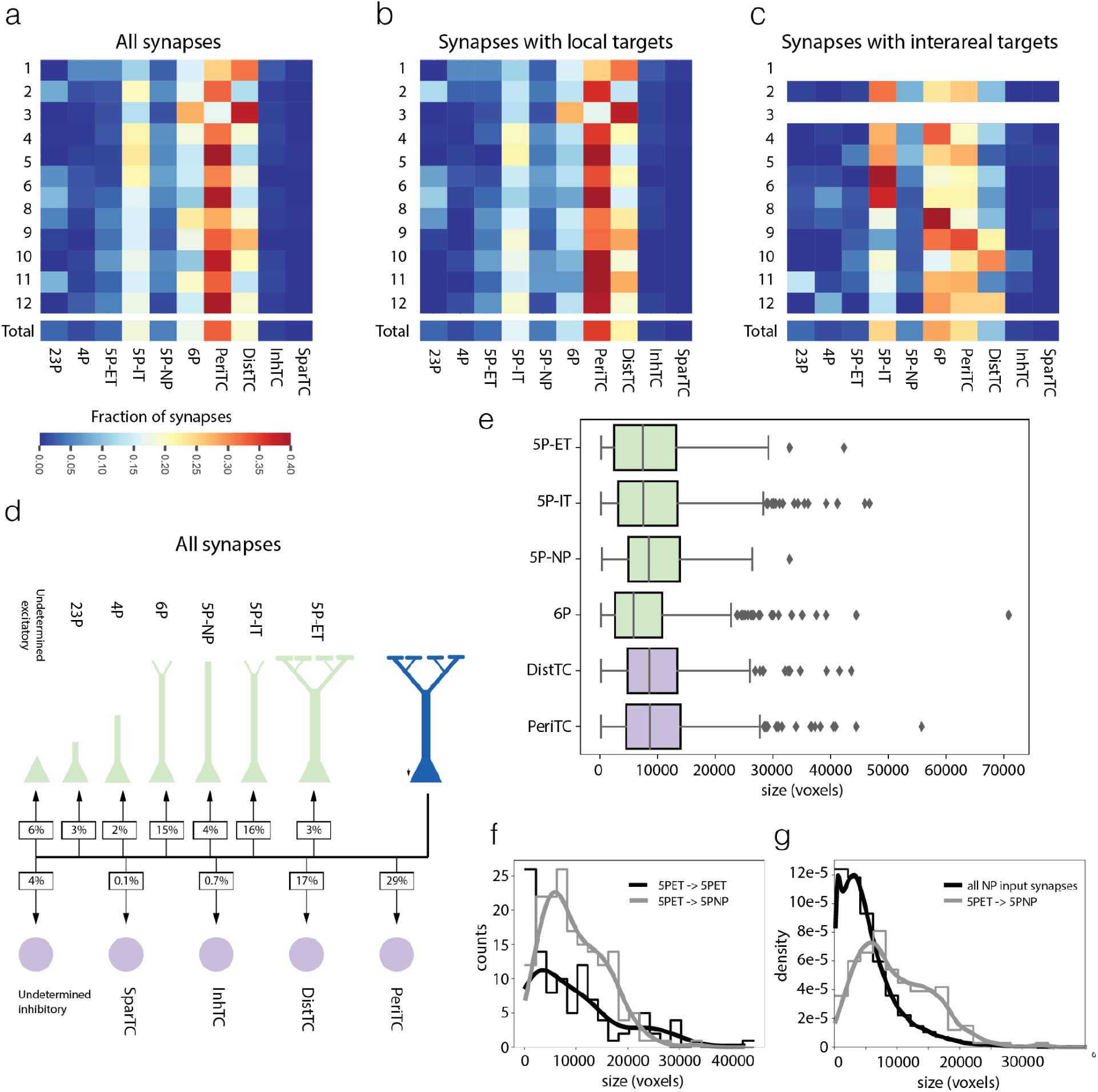
Cell type wiring diagram of ET neurons. **a.** The heat map illustrates the fraction of all synapses (local and inter-areal) formed by ET neurons with target neurons categorized by cell sub-class. Rows 0 to 11 correspond to individual neurons, while the last row represents the cumulative synapses of all neurons. **b)** Heat map as in **(a)** but representing only the synapses formed by the local axon arbor. **c)** Heat map as in **(a)** but representing only the synapses on the inter-areal axon arbor. ET neurons with less than 20 inter-areal synapses are not represented (neurons 1 and 3). **d.** Wiring diagram representing the heat maps of **(a). e.** Boxplots representing the synapse size (measured in voxels) of the synapses formed by ET neurons with different postsynaptic cell subclasses. **f.** Histogram of the distribution of synapse sizes of presynaptic ET to postsynaptic ET connections (black) and presynaptic ET to postsynaptic NP connections (gray). Smoothing of the distribution using kernel density estimation shown as lines. **g.** Histogram of the distribution of synapse sizes on the dendrites of NP neurons (black) and of synapses formed between presynaptic ET to postsynaptic NP connections (gray). Smoding of the distribution using kernel density estimation shown as lines. The distributions are shown as density instead of counts due to the large difference in the numbers of synapses used between the two histograms.

In summary, our findings revealed a circuit motif in which ET neurons form a distinct sub-circuit with inhibitory cells that exert inhibitory control back onto the same excitatory cell types (Fig 2.i). This circuit motif highlights a projection-target specific pattern of reciprocal inhibition and excitation within layer 5.

### Distinct Synaptic Connectivity Patterns of Layer 5 ET Neurons: Differentiating Local and Inter-Areal Connections

In the previous section, we demonstrated that most synapses in the proximal region of the ET axon connect with inhibitory cells. This prompts us to inquire whether the distal segments of the axonal arbor exhibit similar connections with inhibitory cells, or if they instead connect with excitatory cells. Since the distal parts of the ET axon project to neighboring regions, this has significant implications for understanding how ET neurons engage in both local and inter-areal communication. To address this we proofread the axons of 12 ET neurons to their possible full extent within the dataset (Fig 3, Extended Fig 5, four of these neurons are also located in the column, but here we used the full extent of their axonal arbors, and not just the initial 100). All the ET neurons were from layer 5B with clear thick tufted dendrites (Fig 3.a-c, Extended Fig 5) distributed across the dataset (Fig 3.a). Seven of the ET neurons were located in the primary visual cortex (VISp) and five in the higher-order visual area (HVA). The extended ET neurons received 13,095 ± 4,449 (mean ± std, range 9,122-26,796) input synapses per neuron and formed 347 ± 168 (mean ± std, range 183-783) output synapses within the dataset.

**Figure 5.**
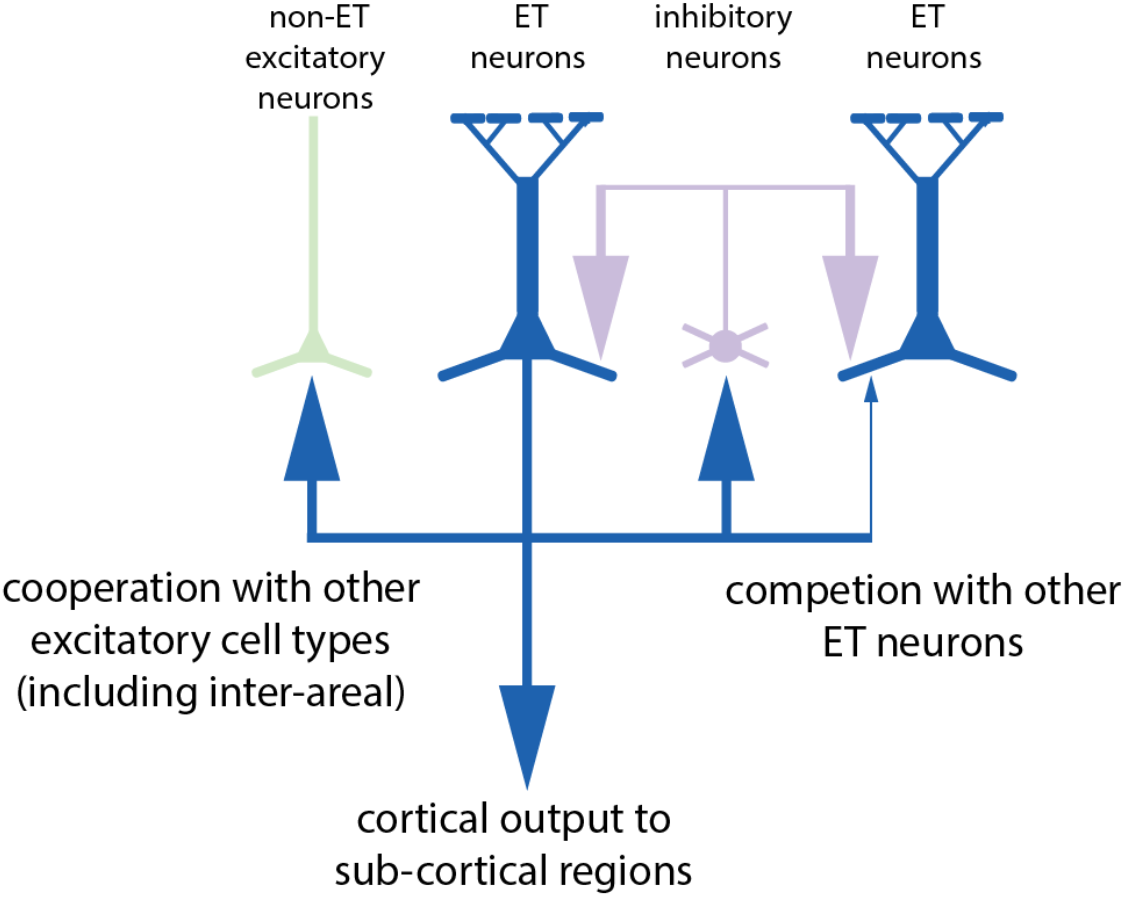
Summary diagram of the connectivity of layer 5 ET neurons.

There were 4156 synapses found on the axons of the 12 proofread ET neurons. As multiple synapses can occur between an ET and its target cells, the 4156 synapses represent 3139 connections with other individual cells in the volume, either with single or multiple synapses (Extended Fig 6). As before we manually proofread all postsynaptic targets that were not associated with a soma (i.e. orphan dendrites and spines) leaving 534 synapses formed with orphan targets.

To identify if the postsynaptic neurons are inhibitory or excitatory we followed the same procedure as before by first using two automated classifications ^60,61^ and then validating by human supervision (see methods) the targets where the two automated classifications disagreed. In total 1734 connections (including some of the orphan dendrites) had manual calls. When one considers the fully extended ET axons (Fig 3.e), 2089 synapses were formed with inhibitory cells (50%) and 1954 synapses were formed with excitatory targets (45%); for the remaining 5% we were unable to determine the class of the target. ET neurons form on average 1.1 synapses with each excitatory target and 1.8 synapses with each inhibitory target, therefore the ET neurons end up forming connections with more excitatory cells (Fig 3.f). The difference between the results of figure 2.d, where we see a preferential targeting of inhibitory cells in the proximal synapses of the axon, and Fig 3.e, where we see a more balanced targeting of inhibition and excitation in the fully extended axons, suggests a distance dependent target of inhibition. This is in fact the case (Fig 3.i) as inhibitory targets vastly dominate the output of the ET axons (in both VISp and HVAs) closer to the soma, even though inhibitory neurons are a minority of cells in the cortex. We observe distance-dependent connectivity patterns that differentially favor the activation of inhibitory and excitatory cells. Given this distance dependency of the target of inhibition and the location of the ET neurons, one corollary of this pattern of connectivity is that local projections favor the targeting of inhibitory neurons and inter-areal projections target more excitatory cells (Fig 3.g-h).

### Diversity of cell types targeted by the ET neurons reveals novel connectivity patterns

The asymmetry in the targeting of excitatory and inhibitory cell classes by ET neurons prompted us to investigate how this asymmetry was reflected in the targeting of specific cell types. While previous studies have predominantly focused on ET-to-ET connections, it remained unclear if ET neurons are indeed the main excitatory targets of other ET cells. Therefore, our aim was to comprehensively map the wiring diagram of ET neurons at the cell type level using the fully extended 12 axons described in the previous section. To achieve this, we classified excitatory and inhibitory postsynaptic cells into subclasses (see Fig 1 and extended Fig 1)^55^. In cases where the automated methods yielded conflicting results, discrepancies were observed between the class and subclass classifications, or no automated classification was available, individual verification of neurons was conducted. Corrections to cell assignments were made as needed during this verification process. The resulting connectivity matrix revealed that the most common excitatory targets of ET neurons were layer 5 IT cells (16% of synapses), followed by layer 6 pyramidal cells (15% of synapses), layer 5 NP cells (4% of synapses), and other layer 5 ET neurons (3% of synapses). Perisomatic targeting cells (29% of synapses) emerged as the most common inhibitory target and distal targeting cells (Martinotti cells, 17% of synapses) also received a considerable fraction of the ET synaptic output. The sizes of synapses in the connections between ET neurons and their postsynaptic targets show considerable overlap, as depicted in Figure 4.e. Notably, there are no significant differences observed among the synapse sizes, except for those with layer 6 pyramidal cells (statistical analysis performed using Kruskal Wallis followed by Post hoc pairwise Conover’s test for multiple comparisons, see Extended Figure 7).

The finding that layer 5 IT cells, or thin tufted neurons, and layer 6 pyramidal cells were the excitatory subclasses receiving most synapses from ET neurons, is unlikely to be due to large mistakes in cell typing given that confusion between IT and ET neurons when using soma features was minimal (10%) ^60^ and cells labeled as Layer 5 IT neurons largely have thin or slender tufted apical dendrites (Extended Fig 11). In addition, analysis of the first 100 synapses of the 39 ET neurons of the column (Fig 2) also shows synapses formed with different excitatory cell types even though with different proportions than in Fig 4 since it is just the very proximal region of the axonal path length (Extended Fig 8).

Beyond the connectivity with ET and IT neurons, we also observe two other excitatory targets that to our knowledge represent previously undescribed connections. The first is the ET to layer 6 pyramidal cells connection suggesting that the ET neurons send an efferent copy of their output to the neurons responsible for conveying feedback information to the thalamus. The second is connections between ET and NP neurons, which is not only novel but surprising given that NP neurons are rare and receive far fewer synapses than other layer 5 neurons ^59^. Given the relatively low density of synapses formed by both ET axons and NP dendrites, it is unlikely for them to randomly encounter each other in the neuropil, suggesting that this can be a highly specific cell type to cell type connection by the ET neurons. Supporting this specificity, the NP spines involved in the ET - NP connection were very long, and could reach more than 10 µm (as shown in the example of Figure 1h). The connections between ET and NP neurons, while comprising a similar fraction of synapses as the ET to ET connections, show distinct distributions in synapse sizes (as depicted in Figure 4.f, KS test p value 0.009). Notably, the ET to NP connection lacks the presence of very small and very large synapses observed in the ET to ET connections. This finding emphasizes the significance of considering cell type-specific differences when analyzing synaptic output ^67^. The MICrONS dataset also provides an opportunity to examine how the ET to NP input compares to the broader distribution of synapse sizes onto NP neurons. Our findings indicate that the ET-NP connection is not only highly specific but also comprises some of the largest synapses that the NP neuron receives (Fig 4.g). This highlights the unique and significant role of the ET neurons in driving synaptic input to NP neurons, and emphasizes the specialized nature of the ET-NP connection.

Together our results uncover a set of new connections from the ET neurons with their excitatory targets that implies that they share their output with other excitatory cell types in the cortical microcircuit. Together with the other findings in this manuscript, our results suggest a circuit architecture where ET neurons compete with each other (or small subnetworks of ET neurons compete with each other) to represent the output of the cortical microcircuits while conveying this output as an efferent copy to other excitatory cell types within and across cortical areas (Fig 5).

## Discussion

In this study we used a 1 mm^3^ electron microscopy volumetric dataset of mouse visual cortex including primary and higher order visual areas ^26^ to analyze the connectivity of the ETs and their synaptic partners. We find that ET neurons form the majority of their local synapses with inhibitory cells that reciprocally target the ET neurons (Figs 2.h and 2.i). The size of the EM volume also allowed us to study for the first time ET inter-areal synapses (Fig 3.k). Of the synapses that are formed with excitatory cell types, only a minority is involved in recurrent excitation with other ET neurons, and the most common targets are layer 5 IT cells as well as layer 6 pyramidal neurons (Fig 4.d). This suggests a circuit architecture that supports local competition between ET neurons and cooperation with other excitatory cell types ^68^, including in inter-areal projections (Fig 5). The fact that layer 5 ET neurons, being the principal output cell of the cortical microcircuit, transmit its output not just subcortically but also to other excitatory cell types is also consistent with theoretical ideas of predictive coding in cortical micro-circuits ^56,57^, that require relaying an efferent copy of the output of a cortical microcircuit to other excitatory cells across the cortical hierarchy.

### Connections with inhibitory cells

Like other cell type groups, the ETs are heterogeneous in their morphology, connectivity, and location within L5 ^6,31,54,69^. Previously, two groups were identified: the upper ETs located in Layer 5A and the lower ones in layer 5B ^6^. In this study we examined 47 ET cells, and for 12 of those (all located in L5B) we fully extended their axonal arbors within the EM volume. The local arbor of most ET neurons overwhelmingly targets inhibitory cells, such a pattern of connectivity has also been described in other excitatory cell types in the mouse ^47,48,63,70^ and it is different from the connectivity results of cat and monkey ^71,72^ where excitatory cells target other excitatory cells in higher proportions. Since neurons in the primary visual cortex of the mouse receive a considerably higher density of synapses than neurons in the primary visual cortex of cat or monkey ^73–78^, it has previously been proposed that a more robust presence of recurrent inhibition is necessary ^47^. However, it’s important to note that the distribution of the fraction of synapses formed with inhibitory cells (Figure 2.d) exhibits a wide range. This suggests significant variations in the downstream targets of individual ET cells or ET cell types.

The findings presented here, along with companion manuscripts ^59,66^ and a recent study ^65^, collectively reveal an interplay between ET neurons and specific inhibitory cell types. This observation not only sheds light on the connectivity of the ET neurons, but also offers a framework for exploring the connectivity of other excitatory cell populations, as for example layer 5 IT and layer 5 NP neurons all inhabit the same space as the layer 5 ET neurons and each also receive input from dedicated inhibitory cell types ^59,65,66^.

Our results strongly indicate that just as there are inhibitory cells that selectively target ET neurons, the ET neurons themselves exhibit a specific preference for targeting these inhibitory cell types in return. This reciprocal targeting pattern highlights a unique and bidirectional relationship between excitatory and inhibitory cell types, potentially playing a crucial role in shaping the functional dynamics of the cortical microcircuit. This strong reciprocity with specific inhibitory cell types has also been recently reported for layer 5 IT neurons ^65^.

The basket cells and the Martinotti cells involved in this microcircuit with ET neurons are morphological types usually found in the deeper layers, especially in Layer 5. The morphology of the basket cell in particular, with a horizontally elongated axonal arbor lying flat in L5 ^41,46,59,79,80^ (Fig 2.f), is perfectly suited to reach the somatic region of ETs. Since the arbors of these basket cells can spread extensively in layer 5 (400 to 800 µm), they also can inhibit a large number of ETs. While we show that the basket cells are reciprocally interconnected with ETs, they are also known to target each other, commonly form autapses, and are coupled with gap junctions, forming a specific subnetwork in the cortex suited for generating oscillation ^81–88^. These oscillations can then serve as a clock, and this clock has been argued to provide a temporal window for neuronal integration or work as a coincidence detector ^85^. As the frequency of the oscillations is tied to the strength of the reciprocal connections between excitatory and inhibitory cells it is worth considering how the connectivity patterns that we observe with the ET neurons affects this integration window. To support this discussion we used a network model of generalized leaky-integrate-and-fire (GLIF) neurons representing the L5 ET circuit, consisting of L5 ET cells, L5 Basket Cells, and L5 Martinotti Cells (Extended Data Figure 9.a and 9.b). This model was based upon an earlier model of the mouse primary visual cortex ^89^ from which we extracted the neuronal models for L5, but varied the connectivity (see Methods) to different configurations, including the one matching the experimental findings reported here. As expected, the L5 ET cells in the circuit exhibit oscillations in their firing rates. Configurations of the circuit that reflect the observed connectivity with strong recurrence between the L5 ET neurons and inhibitory cell types (and sparse connections among L5 ET neurons) shifts the oscillations to higher frequencies (Extended Data Figure 9.h) in the gamma range. Conversely, configuration where we reduce the fraction of synapses associated with the recurrent interaction with inhibitory cells or remove recurrency all together in the circuit model lowers the frequency of oscillations (Extended Data Figures 9.f, 9.g and 9.h).

Not only the basket cells described above, but also Martinotti cells, that synapse onto ETs ^90^ contributing to this oscillations, are identified as nicotinic acetylcholine receptor α2 subunit (Chrna2)-positive Martinotti cells (Chrna2MCs) by a previous study ^44^. We also found that ET-preferring Martinotti cells were predicted to belong to a Chrna2+ cell type when classified by their morphology (Gamlin et al., 2023). The motif of ETs-Chrna2MCs-ETs is observed in many cortical areas, and one functional consequence of this motif is frequency-dependent disynaptic inhibition (FDDI). However, the function of FDDI within the broader cortical circuitry is still in debate ^40,44,91–95.^.

One common used network motif to describe the interaction between excitation and inhibition in cortical microcircuits is the one of winner take all networks (WTA). Such a network would be well poised to provide an effective mechanism to select the population of L5 ETs that are active at any given point in time, thus selecting and limiting the cortical output. The configuration of the connectivity, plays a role on how the WTA behaves. To illustrate this concept, we utilized the circuit model described above (Extended Data Figure 9). By increasing the fraction of synapses involved in the recurrent connections between ET neurons and inhibitory cells (and reducing the recurrent connections between ET to ET neurons), as we observe in the data, leads to a sharpening of the tuning properties of the neuron (Extended Data Figure 9.c,d,e) compared to a configuration with less synapses involved in the loop between ET neurons and inhibitory cells. Combined, the modeling results indicate that sparse L5 ET targets and extensive innervation of inhibitory cells by L5 ET cells, observed in the EM data, may support sharpening in processing of stimuli, as well as gamma-frequency firing in L5 (Extended Data Figure 9).

Key questions remain unanswered in the interplay between ET neurons and their inhibitory partners, which could be addressed in future studies. Firstly, shedding light on whether different inhibitory cell subclasses act synergistically or through disinhibition on the ETs is essential. Since perisomatic targeting cells primarily target the somatic region and distal targeting cells target the distal-apical dendritic region of the ETs, understanding their interplay would provide insights into ETs’ apical dendritic signal processing. Additionally, while our findings focus on cell-type interactions, it is well-established that ET neurons form subnetworks among themselves. Further exploring the interplay of inhibition with these sub-networks would be important for a comprehensive understanding.

### Comparison with previous mapping of ET connections with excitatory neurons

Most of the previous work on the connectivity of ET neurons has focused on recurrent connections with other ET neurons ^6,30,50–52^, and while we do find ET to ET connections we also find that other excitatory cell types receive a much higher fraction of the synaptic output of ET neurons. Therefore, our results reveal a much richer synaptic architecture than previously described, but they also pose the conundrum as to why these cell types connections have not been previously noted. For some connections, such as the ET to Layer 6 pyramidal cells or the ET to NP cells, the answer might be as simple as no one was looking for them, though our results are consistent with previous reports of a diversity of pyramidal cell targets <u>(Kozloski et al. 2001)</u>. Most of the connectivity studies on ET cells have been based on paired recordings that target the cell types of interest (e.g. ET to ET connections) and extensive sampling of all possible connections has not been realized. In this respect, in EM volumes such as the one used in this study ^26^ (www.microns-explorer.org) once an axon is proofread and all cell types in the volume are classified ^60^, one can obtain a comprehensive list of the synaptic partners. To minimize bias, we not only proofread the axons of the ET cells but also manually checked and corrected the synapses onto orphan objects (e.g. detached spines) to ensure that the results were not due to any bias in the automatic segmentation. The very low rate of detached spines (<5% of synapses) makes it unlikely to affect our results.

The findings of this study, which reveal a substantial fraction of synapses formed by ET neurons with layer 5 thin tufted or IT neurons, may initially appear to be inconsistent with the prevailing interpretation of the existing literature ^39^ that focus on recurrent connections between ET to ET neurons and assumes ET to IT connections to be very rare. While ET to IT connections have been predicted ^98^ and described ^31,54^, they have also been unreported in other studies ^30^. Synapse size or number of synapses per connection in the ET to IT connection are unlikely to be the solution to this conundrum since they are similar across the different excitatory cell types targeted by the ET neurons. The exact choice of ET neurons to fully reconstruct in this manuscript, which are from layer 5b, is also unlikely to be the reason, since targeting of layer 5 IT cells is also clear from all the other layer 5 ET neurons analyzed (see results and Extended Fig 8). One important consideration is that the connection probability that is typically measured in slice electrophysiology experiments, and the cell type specific output distribution of synapses we report in this study are affected differently by factors such as the number and spatial distribution of potential targets. For example, if a post synaptic cell type is very common but receives relatively few synapses, it might be difficult to find the exact individual pairs of this connection with paired recordings. Given that L5-IT cells outnumber ET cells but receive fewer synapses per neuron ^59^, we wanted to more directly compare our data to previous reports.

Therefore we decided to simulate a large-scale connectivity experiment based on slice electrophysiology where VISp neurons were labeled with a cell-type specific Cre line and pairs of nearby labeled cells were probed for the existence of a synaptic connection ^30^. In the electrophysiology experiment, sampling was biased toward smaller lateral distances (Extended Data Figure 10.a), where connections would be more likely. To match this sampling experiment in the EM data, for each fully reconstructed L5-ET cell in VISp, we assembled a collection of all L5-IT and L5-ET cells and assigned a sample weight based on lateral distance to match the empirical sampling (Extended Data Figure 10.b). We then performed 10,000 weighted random draws of the same number of pairs as in Campagnola et al (N=82 for L5-IT targets, N=739 for L5-ET targets) and counted the number of synaptically connected pairs observed per draw. For L5-IT targets the observed number of connections in the synapse physiology experiment was zero, while the EM-based samples had a median of two connections; 5.2% of draws showed zero connections (Extended Data Figure 10.c), making this difference barely non-significant. For L5-ET targets, the observed number of connections was 55, while the EM-based samples had a median of 75 connections (Extended Data Figure 10.d). We counted 55 or less connections in only 0.5% of draws, thus the EM data did show significantly more ET-to-ET connections than observed, but only by approximately 1.4 times (75/55 = 1.36). Together, our results qualitatively agree with previous observations that the probability of connection is higher in the L5-ET to L5-ET connection than in the L5-ET to L5-IT connection but also demonstrate how different measures can highlight different aspects of connectivity and reveal previously unknown connections between cell types..

### A road map for the comprehensive mapping of cell type connectivity

This manuscript focuses on the connectivity of ET neurons but the approach presented here can be applied and iterated across all other cortical excitatory cell types. This is possible because volumes such as the Microns dataset are large enough to contain a comprehensive map of local connectivity, allowing us to unveil a comprehensive map of connections with post-synaptic partners as we have done here for the ET neurons.

One important aspect of this characterization is the concept of cell types, which have long been a focal point in connectivity studies, but this and other recent connectivity studies ^59,65,66^ show that the specificity in cell-type to cell-type connectivity is much higher than previously described. This increasing recognition of specificity highlights the importance of connectivity as a critical component of defining cell-types themselves. While here we describe a single excitatory cell type, this work demonstrates the feasibility of this comprehensive mapping and how it can unveil new connections, even in cell types that historically have been extensively studied. This approach opens the way for a mapping of the cortical microcircuit at synapse resolution not only across all other cell types, but also across cortical areas. While this connectivity certainly constrains the computations that the cortex can perform, its operation also depends on gene expression and its functional properties at any point in time. It is not a stretch of the imagination to think that in the near future one will be able to relate connectomics, gene expression and functional properties of all cells in the cortex, and in an accompanying paper we begin to relate these properties for a subset of inhibitory cells ^99^.

## Methods

### Mouse Lines

All procedures were approved by the Institutional Animal Care and Use Committees (IACUC) of Baylor College of Medicine and the Allen Institute. All results described here are from a single male mouse, age 65 days at onset of experiments, expressing GCaMP6s in excitatory neurons via SLC17a7-Cre and Ai162 heterozygous transgenic lines (recommended and generously shared by Hongkui Zeng at Allen Institute for Brain Science; JAX stock 023527 and 031562, respectively). In order to select this animal, 31 (12 female, 19 male) GCaMP6-expressing animals underwent surgery as described below. Of these, 8 animals were chosen based on a variety of criteria including surgical success and animal recovery, the accessibility of lateral higher visual areas in the cranial window, the degree of vascular occlusion, and the success of cortical tissue block extraction and staining. Of these 8 animals, one was chosen for 40 nm slicing and EM imaging based on overall quality using these criteria.

### Timeline

DOB: 12/19/17

Surgery: 2/21/18 (P64)

Two-photon imaging start: 3/4/18 (P75)

Two-photon imaging end: 3/9/18

(P80) Structural Stack: 3/12/18 (P83)

Perfusion: 3/16/18 (P87)

### Surgery and Two Photon Imaging

Before the anatomical work described in this study the mouse underwent surgery and in vivo 2-photon calcium imaging at Baylor College of Medicine. The details of this *in vivo* functional imaging and surgeries are described in MICrONS Consortium et al ^26^.

### Tissue Preparation

After optical imaging at Baylor College of Medicine, candidate mice were shipped via overnight air freight to the Allen Institute. All procedures were carried out in accordance with the Institutional Animal Care and Use Committee at the Allen Institute for Brain Science. All mice were housed in individually ventilated cages, 20-26 C, 30-70% Relative Humidity, with a 12-hour light/dark cycle. Mice were transcardially perfused with a fixative mixture of 2.5% paraformaldehyde, 1.25% glutaraldehyde, and 2 mM calcium chloride, in 0.08 M sodium cacodylate buffer, pH 7.4. After dissection, the neurophysiological recording site was identified by mapping the brain surface vasculature. A thick (1200 um) slice was cut with a vibratome and post-fixed in perfusate solution for 12 – 48 h. Slices were extensively washed and prepared for reduced osmium treatment (rOTO) based on the protocol of Hua and colleagues ^100^. All steps were performed at room temperature, unless indicated otherwise. 2% osmium tetroxide (78 mM) with 8% v/v formamide (1.77 M) in 0.1 M sodium cacodylate buffer, pH 7.4, for 180 minutes, was the first osmication step. Potassium ferricyanide 2.5% (76 mM) in 0.1 M sodium cacodylate, 90 minutes, was then used to reduce the osmium. The second osmium step was at a concentration of 2% in 0.1 M sodium cacodylate, for 150 minutes. Samples were washed with water, then immersed in thiocarbohydrazide (TCH) for further intensification of the staining (1% TCH (94 mM) in water, 40 ℃, for 50 minutes). After washing with water, samples were immersed in a third osmium immersion of 2% in water for 90 minutes. After extensive washing in water, lead aspartate (Walton’s (20 mM lead nitrate in 30 mM aspartate buffer, pH 5.5), 50 ℃, 120 minutes) was used to enhance contrast. After two rounds of water wash steps, samples proceeded through a graded ethanol dehydration series (50%, 70%, 90% w/v in water, 30 minutes each at 4 ℃, then 3 x 100%, 30 minutes each at room temperature). Two rounds of 100% acetonitrile (30 minutes each) served as a transitional solvent step before proceeding to epoxy resin (EMS Hard Plus). A progressive resin infiltration series (1:2 resin:acetonitrile (e.g. 33% v/v), 1:1 resin:acetonitrile (50% v/v), 2:1 resin acetonitrile (66% v/v), then 2 x 100% resin, each step for 24 hours or more, on a gyratory shaker) was done before final embedding in 100% resin in small coffin molds. Epoxy was cured at 60 °C for 96 hours before unmolding and mounting on microtome sample stubs.

The sections were then collected at a nominal thickness of 40 nm using a modified ATUMtome (RMC/Boeckeler) onto 6 reels of grid tape ^101,102^.

### Transmission Electron Microscopy Imaging

The parallel imaging pipeline described here^102^ converts a fleet of transmission electron microscopes into high-throughput automated image systems capable of 24/7 continuous operation. It is built upon a standard JEOL 1200EX II 120kV TEM that has been modified with customized hardware and software. The key hardware modifications include an extended column and a custom electron-sensitive scintillator. A single large-format CMOS camera outfitted with a low distortion lens is used to grab image frames at an average speed of 100 ms. The autoTEM is also equipped with a nano-positioning sample stage that offers fast, high-fidelity montaging of large tissue sections and an advanced reel-to-reel tape translation system that accurately locates each section using index barcodes for random access on the GridTape. In order for the autoTEM system to control the state of the microscope without human intervention and ensure consistent data quality, we also developed customized software infrastructure piTEAM that provides a convenient GUI-based operating system for image acquisition, TEM image database, real-time image processing and quality control, and closed-loop feedback for error detection and system protection etc. During imaging, the reel-to-reel GridStage moves the tape and locates targeting aperture through its barcode. The 2D montage is then acquired through raster scanning the ROI area of tissue. Images along with metadata files are transferred to the data storage server. We perform image QC on all the data and reimage sections that fail the screening.

### Volume Assembly

The volume assembly pipeline used in this study^58,103^ starts by first correcting the images in the serial section for lens distortion effects. A non-linear transformation of higher order is computed for each section using a set of 10 x 10 highly overlapping images collected at regular intervals during imaging. The lens distortion correction transformations should represent the dynamic distortion effects from the TEM lens system and hence require an acquisition of highly overlapping calibration montages at regular intervals. Overlapping image pairs are identified within each section and point correspondences are extracted for every pair using a feature based approach. In our stitching and alignment pipeline, we use SIFT feature descriptors to identify and extract these point correspondences. Per image transformation parameters are estimated by a regularized solver algorithm. The algorithm minimizes the sum of squared distances between the point correspondences between these tile images. Deforming the tiles within a section based on these transformations results in a seamless registration of the section. A downsampled version of these stitched sections are produced for estimating a per-section transformation that roughly aligns these sections in 3D. A process similar to 2D stitching is followed here, where the point correspondences are computed between pairs of sections that are within a desired distance in z direction. The per-section transformation is then applied to all the tile images within the section to obtain a rough aligned volume. MIPmaps are utilized throughout the stitching process for faster processing without compromise in stitching quality.

The rough aligned volume is rendered to disk for further fine alignment. The software tools used to stitch and align the dataset is available in our github repository https://github.com/AllenInstitute/render-modules. The volume assembly process is entirely based on image meta-data and transformations manipulations and is supported by the Render service (https://github.com/saalfeldlab/render).

Cracks larger than 30 um in 34 sections were corrected by manually defining transforms. The smaller and more numerous cracks and folds in the dataset were automatically identified using convolutional networks trained on manually labeled samples using 64 x 64 x 40 nm^3^ resolution image. The same was done to identify voxels which were considered tissue. The rough alignment was iteratively refined in a coarse-to-fine hierarchy (Wetzel et al. 2016), using an approach based on a convolutional network to estimate displacements between a pair of images (Mitchell et al. 2019). Displacement fields were estimated between pairs of neighboring sections, then combined to produce a final displacement field for each image to further transform the image stack. Alignment was first refined using 1024 x 1024 x 40 nm^3^ images, then 64 x 64 x 40 nm^3^ images.

The composite image of the partial sections was created using the tissue mask previously computed. Pixels in a partial section which were not included in the tissue mask were set to the value of the nearest pixel in a higher-indexed section that was considered tissue. This composite image was used for downstream processing, but not included with the released images.

### Segmentation

Remaining misalignments were detected by cross-correlating patches of image in the same location between two sections, after transforming into the frequency domain and applying a high-pass filter. Combining with the tissue map previously computed, a mask was generated that sets the output of later processing steps to zero in locations with poor alignment. This is called the segmentation output mask.

Using the method outlined in Lee et al.^104^, a convolutional network was trained to estimate inter-voxel affinities that represent the potential for neuronal boundaries between adjacent image voxels. A convolutional network was also trained to perform a semantic segmentation of the image for neurite classifications, including (1) soma+nucleus, (2) axon, (3) dendrite, (4) glia, and (5) blood vessels. Following the methods described in Wu et al. ^105^, both networks were applied to the entire dataset at 8 x 8 x 40 nm^3^ in overlapping chunks to produce a consistent prediction of the affinity and neurite classification maps. The segmentation output mask was applied to the predictions.

The affinity map was processed with a distributed watershed and clustering algorithm to produce an over-segmented image, where the watershed domains are agglomerated using single-linkage clustering with size thresholds ^106,107^. The over-segmentation was then processed by a distributed mean affinity clustering algorithm ^106,107^ to create the final segmentation. We augmented the standard mean affinity criterion with constraints based on segment sizes and neurite classification maps during the agglomeration process to prevent neuron-glia mergers as well as axon-dendrite and axon-soma mergers.

### Synapse detection & assignment

A convolutional network was trained to predict whether a given voxel participated in a synaptic cleft. Inference on the entire dataset was processed using the methods described in ^105^ using 8 x 8 x 40 nm^3^ images. These synaptic cleft predictions were segmented using connected components, and components smaller than 40 voxels were removed.

A separate network was trained to perform synaptic partner assignment by predicting the voxels of the synaptic partners given the synaptic cleft as an attentional signal ^108^. This assignment network was run for each detected cleft, and coordinates of both the presynaptic and postsynaptic partner predictions were logged along with each cleft prediction.

### Nucleus detection

A convolutional network was trained to predict whether a voxel participated in a cell nucleus. Following the methods described in ^105^, a nucleus prediction map was produced on the entire dataset at 64 x 64 x 40 nm^3^. The nucleus prediction was thresholded at 0.5, and segmented using connected components.

### Selection of L5 ET cells for analysis

For the analysis of ET cells in the column shown in Figure 2 we used all the cells identified as ET using the dendritic and synapse features described below (see: *Quantitative characterization of excitatory neuron morphological subclasses*). As layer 5A and layer 5B ^6^ are known to contain different types of ET neurons, for the 12 ET neurons with comprehensive extended axons (Figures 3 and 4) we selected cells from the lower portion of L5 (L5B). As we wanted to compare local and inter-areal targeting patterns we chose cells for which the inter-areal portion of the axon was in good condition.

### Manual proofreading of dendritic and axonal processes

Following the methods described previously ^67,109^ proofreaders used a modified version of Neuroglancer with annotation capabilities as a user interface to make manual split and merge edits to neurons. Proofreading was aided by on-demand highlighting of branch points and tips on user-defined regions of a neuron based on rapid skeletonization (https://github.com/AllenInstitute/Guidebook). This approach quickly directed proofreader attention to potential false merges and locations for extension, as well as allowed a clear record of regions of an arbor that had been evaluated. For dendrites, we checked all branch points for correctness and all tips to see if they could be extended. False merges of simple axon fragments onto dendrites were often not corrected in the raw data, since they could be computationally filtered for analysis after skeletonization (see next section). Detached spine heads and detached dendrites of postsynaptic targets of ET neurons were also proofread to evaluate the possibility of merging them to a segment containing a soma. Dendrites that were proofread are identified in CAVE table “proofreading_status_public_release” as “clean”

Axons of ET neurons and inhibitory neurons considered in this study began by “cleaning” axons of false merges by looking at all branch points. We then performed an extension of axonal tips, the degree of this extension depended on the neuron:

1. Comprehensive extension: For the 12 ET neurons used in Figures 3 and 4, each axon end and branch point was visited and checked to see if it was possible to extend until either their biological completion or reached an incomplete end (incomplete ends were due to either the axon reaching the borders of the volume or an artifact that curtailed its continuation).
2. Substantial extension: For the inhibitory neurons used in this study, each axon branch point was visited and checked, many but not all ends were visited and many but not all ends were done.
3. At least 100 synapses: For all other ET neurons (with the exception of the 12 that were comprehensively extended), the axons were extended until at least 100 synapses were present on the axon to get a sampling of their local output connectivity profile.

Axons that were proofread are identified in CAVE table “proofreading_status_public” as “clean” and the proofreading strategy associated with each axon is described in the CAVE table “proofreading_strategy”.

### Quantitative characterization of excitatory neuron morphological subclasses

To quantitatively characterize the excitatory neuron morphology of cells that were part of the column shown in Figure 2.a, we computed features based only on neuron dendrites and soma. This has been described in the accompanying manuscript of Schneider-Mizell et al ^59^ for morphological types (m-types) and was also used to train the models that use soma and nucleus features ^60^. For the present manuscript we group all the layer 2/3 m-types into a single subclass, we group all the layer 4 m-types into a single subclass and we group all the layer 6 m-types into a single subclass. We also group all the layer 5 inter-telencephalic (thin or slim tufted dendrites) m-types into a single sub-class.

As described in Schneider-Mizell et al ^59^ the features were:

1. Median distance from branch tips to soma per cell.
2. Median tortuosity of the path from branch tips to soma per cell. Tortuosity is measured as the ratio of path length to the Euclidean distance from tip to soma centroid.
3. Number of synaptic inputs on the dendrite.
4. Number of synaptic inputs on the soma.
5. Net path length across all dendritic branches.
6. Radial extent of dendritic arbor. We define “radial distance” to be the distance within the same plane as the pial surface. For every neuron, we computed a pia-to-white-matter line, including a slanted region in deep layers, passing through its cell body. For each skeleton vertex, we computed the radial distance to the pia-to-white-matter line at the same depth. To avoid any outliers, the radial extent of the neuron was defined to be the 97th percentile distance across all vertices.
7. Median distance to soma across all synaptic inputs.
8. Median synapse size of synaptic inputs onto the soma.
9. Median synapse size of synaptic inputs onto the dendrites.
10. Dynamic range of synapse size of dendrite synaptic inputs. This was measured as the difference between 95th and 5th percentile synapse sizes.
11. Shallowest extent of synapses, based on the 5th percentile of synapse depths.
12. Deepest extent of synapses, based on the 95th percentile of synapse depths.
13. Vertical extent of synapses, based on the difference between 95th and 5th percentile of synapse depths.
14. Median linear density of synapses. This was measured by computing the net path length and number of synapses along 50 depth bins from layer 1 to white matter and computing the median. A linear density was found by dividing synapse count by path length per bin, and the median was found across all bins with nonzero path length.
15. Median radius across dendritic skeleton vertices. To avoid the region immediately around the soma from having a potential outlier effect, we only considered skeleton vertices at least 30 micron from the soma.

Three additional sets of features used component decompositions. To more fully characterize the absolute depth distribution of synaptic inputs, for each excitatory neuron, we computed the number of synapses in each of 50 depth bins from the top of layer 1 to the surface of white matter (bin width approx 20 µm). We z-scored synapse counts for each cell and computed the top six components using SparsePCA.The loadings for each of these components based on the net synapse distribution were used as features.

To characterize the distribution of synaptic inputs relative to the cell body instead of cortical space, we computed the number of synapses in 13 soma-adjusted depth bins starting 100 \Micron above and below the soma. As before, synapse counts were z-scored and we computed the top five components using SparsePCA.The loadings for each of these components were used as additional features.

To characterize the relationship with branching to distance, we measured the number of distinct branches as a function of distance from the soma at ten distances, every 30 µm starting at 30 µm from the soma and continuing to 300 µm.

For robustness relative to precise branch point locations, the number of branches were computed by finding the number of distinct connected components of the skeleton found in the subgraph formed by the collection of vertices between each distance value and 10 µm toward the soma.We computed the top three singular value components of the matrix based on branch count vs distance for all excitatory neurons, and the loadings were used as features.

All features were computed after a rigid rotation of 5 degrees to flatten the pial surface and translation to zero the pial surface on the y axis. Features based on apical classification were not explicitly used to avoid ambiguities based on both biology and classification. Using this collection of features, we clustered excitatory neurons by running phenograph ^110^ 100 times with 97\% of cells included each time. Phenograph finds a nearest neighborhood graph based on proximity in the feature space and clusters by running the Leiden algorithm for community detection on the graph. Here, we used a graph based on 10 nearest neighbors and clustered with a resolution parameter of 1.3. These values were chosen to consistently separate layer 5 ET, IT, and NP cells from one another, a well established biological distinction. A co-clustering matrix was assembled with each element corresponding to the number of times two cells were placed in the same cluster. To compute the final consensus clusters, we performed agglomerative clustering with complete linkage based on the co-clustering matrix, with the target number of clusters set by a minimum Davies-Bouldin score and a maximum Silhouette score.

Clusters were then named based on the most frequent manually defined cell type within the cluster and reordered based on median soma depth.

### Automated Cell Class and Subclass Classification using somatic features

We analyzed the nucleus segmentations for features such as volume, surface area, fraction of membrane within folds, and depth in cortex. We trained an SVM machine classifier to use these features to detect which nucleus detections were likely neurons within the volume, with 96.9% precision and 99.6% recall. This model was trained based upon data from an independent dataset, and the performance numbers are based upon evaluating the concordance of the model with the manual cell type calls within the volume. This model predicted 82,247 neurons within the larger subvolume. For the neurons, we extracted additional features from the somatic region of the cell, including its volume, surface area, and density of synapses. Dimensionality reduction on this feature space revealed a clear separation between neurons with well segmented somatic regions (n=69,957) from those with fragmented segmentations or sizable merges with other objects (n=12,290). Combining those features with the nucleus features, we trained a multi-layer perceptron classifier to distinguish excitatory from inhibitory neurons amongst the well-segmented subset, using the 80% of the manual labeled data as a training set, and 20% as a validation set to choose hyper-parameters. After running the classifier across the entire dataset, we then tested the performance by sampling an additional 350 cells (250 excitatory and 100 inhibitory). We estimate from this test that the classifier had an overall accuracy of 97% with an estimated 96% precision and 94% recall for inhibitory calls.

### Expert Cell Classification

In the cases where there wasn’t consensus between the NEURD classification and the nucleus features classifications regarding the class (excitatory/inhibitory) the decision was made by humans. Neurons that had spiny dendrites and formed asymmetric synapses with mostly spines were assigned the excitatory class, while neurons that had smooth or spiny dendrites but form more symmetric synapses that target mostly spines and dendritic shafts were labeled as inhibitory. Occasionally subclass labels were also assigned by humans according to the following criteria. L5 ET cells have a large soma and large folded nucleus, a thick apical dendrite which forms a large apical tuft in the upper layers of the cortex. ET apical dendrites sometimes form 2 main branches around the middle of the cortex and both branches continue parallel to each other, upwards. Beside the dense basal dendrites it also has several dendritic branches coming out from the apical dendrite closely above the soma. Dendrites covered densely with spines beside the short proximal segment.

Layer 5 intra-telencephalic cell (IT), which were recognised based on smaller soma and thinner apical dendrites, also a small or sometimes missing apical tuft. Layer 5 near -projecting cell (NP) had small soma and very few, thin and long basal dendrites and one thin apical with small or missing tuft. This cell type has very few dendritic spines and even some of those spines have very long atypical spine necks.

For this study the two most relevant inhibitory cell subclasses were the perisomatic targeting cells (also described as basket cells^111^) and the distal targeting cells (most of them Martinotti cells ^112^). The perisomatic targeting cells formed boutons surrounding the soma and proximal dendrites of pyramidal (or other) cells. They usually have large soma, and smooth dendrites (though there were a few examples where they could be spiny), and a large axon arbor which is heavily myelinated. Boutons are usually large and often several of them target the same pyramidal cell soma or proximal region. The distal targeting cells had usually two axonal arborization, one in the upper layers reaching layer 1 and one in the lower layers. They rarely form synapses with the soma of pyramidal cells and their dendrites are very spiny.

Whenever possible we also labeled manually dendritic orphans according to their class (excitatory or inhibitory). Dendrites were labeled as excitatory when they were rich on spines with smooth rounded heads and had few shaft synapses (that were largely but not exclusively inhibitory synapses). Two types of dendrites were labeled as inhibitory. The first type was smooth, rich in mitochondria and received excitatory synapses on the shaft. The second type had a rugged surface with spines (also rugged) and received excitatory input also on the shaft. Orphan spines were left unclassified since the ET neurons formed synapses with excitatory and inhibitory spines. Small segments of dendrites that were sparsely spiny and the spines had smooth heads were also left unclassified because they could either belong to a sparsely spiny inhibitory cell or a layer 5 NP pyramidal neuron.

### L5 circuit model with generalized leaky-integrate-and-fire (GLIF) neurons

#### Model

The L5 circuit model was built as a modified version of a bio-realistic model of the mouse primary visual cortex ^89^, from which we extracted the neuronal models for L5 (Extended Data Fig 9.a and 9.b). The L5 circuit model consists of 3 cell types: L5 ET cells, L5 Pvalb cells (Basket Cells), and L5 Sst cells (Martinotti Cells), which are represented by 40 unique generalized leaky-integrate-and-fire (GLIF) neuron models. The cells were uniformly distributed in a cylindrical domain with a radius 650 um and a specified depth range for L5 (-500 to -660 um), according to experimentally observed densities. To avoid boundary artifacts ^113^, our analyses focused on the cells within the core cylinder (radius < 200 um). Within this core cylinder, there are a total of 1134 point neurons, including 639 L5 ET cells, 242 L5 Pvalb cells, and 253 L5 Sst cells (number of cells in the whole model: 6523 (L5 ET), 2866 (L5 Pvalb), and 2537(L5 Sst)). The synaptic connection properties, such as connection probability and synaptic weight distributions, were modeled based on experimental data from the Allen Institute, including the Synaptic Physiology dataset [portal.brain-map.org/explore/connectivity/synaptic-physiology] ^30^ and the IARPA MICrONS dataset [www.microns-explorer.org/cortical-mm3] ^26^.

Here we specifically adjusted the connection probabilities of L5 ET cells to these 3 cell types based on the 12 proofread L5 ET cells. In the base model developed using these data, 8% of all connections from L5 ET cells target other L5 ET cells, 49% target L5 Pvalb cells (PeriTC, basket), and 43% target L5 Sst cells (distTC, Martinotti).

To examine how the fraction of inhibitory cells among the L5 ET synaptic targets affect the activity of the model, we also built models with various fractions of inhibitory vs. excitatory neurons among the targets of connections from L5 ET neurons, in addition to the base model. Specifically, the fraction is the number of L5 ET connections targeting L5 Pvalb and L5 Sst (with a fixed ratio of around 49:43 as noted above) divided by the total number of connections from L5 ET cells. We built 11 model variations with the fraction ranges from 0 to 1, incremented by 0.1. The total number of the L5 ET connections, the synaptic weight distribution, and connection rules are the same for all these models.

#### Simulation configuration

The cells received external inputs from two sources. The first is background input, which roughly represents the input from the rest of the brain outside this circuit. Here we incorporated 100 background input units that fire at 250 Hz with a Poisson distribution of spikes. Each cell in the model received input from 4 randomly selected units to introduce variability in the circuit. The weights were optimized to generate spiking activities consistent with the spontaneous firing rates observed in the Allen Institute Visual Coding Neuropixels data (Siegle et al., 2021). For the optimization process, we adopted a two-step approach. In the first step, we optimized the background weights for the model without recurrent connections using dichotomy. During this process, we conducted a simulation with an initial set of background weights with sufficiently large values. Through iterative adjustments using dichotomy optimization, we updated the weights until the firing rate of each cell model converges to the target firing rate with a 5% tolerance. In the second step, we utilized the optimized weights obtained from the first step as the initial condition for the model with recurrent connections. During each iteration, we updated the weight by 5% for the cell type that exhibited the largest difference from the target firing rate. This iterative procedure continued until the firing rate of all cell types reached a convergence within 10% of the target firing rate. We performed this optimization process respectively for the base model and each model variation. Notably, we also experimented with optimizing the background weights for the base model only and then applying those optimized weights to the model variations. The results were found to be similar to the results presented here.

The second source of input to the L5 circuit model was also a Poisson spike train at 1 kHz, but with the weight to each neuron determined by a Gaussian function (Extended Data Fig 9.a). In the model, each neuron i was assigned a preferred angle θ*_i_*, spanning from 0 to 360 degrees, forming a ring in this abstract preferred angle space. This Gaussian input is an abstract representation of an external input that targets a certain angle θ_target_. Here, the weight of the Gaussian input to neuron i was 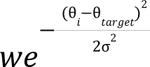, where w is the maximum weight, which was tuned to make the firing rates of the neurons at the target angle consistent with the firing rates at the preferred direction of the drifting gratings stimulus observed in the Visual Coding Neuropixels data (Siegle et al., 2021), and is the standard deviation of the Gaussian function, which was set to 60 degrees. This weight is tuned for the base model and then applied to all the model variations.

Here we conducted simulations for each model under 2 conditions: with recurrent connections and without the recurrent connections. The external input weights are the same for both conditions, which were optimized for the model with recurrent connections as described above. For each condition of each model, we conducted 24 simulation trials, with the Gaussian input target angle θ*_target_* ranging from 0 to 360 degrees, incremented by 15 degrees.

#### Gaussian curve fitting for the firing rates of L5 ET cells

In each simulation trial, we analyzed the average firing rate of the L5 ET cells as a function of their preferred angle relative to the target angle of the Gaussian input (referred to as the “relative angle”, Extended Data Figure 9.c). The L5 ET cells were categorized based on their relative angles, with a bin size of 10 degrees, to calculate the average firing rate for each bin. Subsequently, we fit a Gaussian curve to these average firing rates using least squares optimization. The optimization process was performed using the “optimize.curve_fit()” function from the SciPy Python package.

#### Power spectrum analysis for the population firing rates of the L5 ET cells

In each simulation trial, we conducted power spectrum analysis on the population firing rates of the L5 ET cells. The power spectrum was estimated using the total spike counts of all L5 ET cells with a time window of 2.5 ms (total duration: 9 sec). Welch’s method was employed for the analysis, using a segment size of 500 ms and a 50% overlap (resulting in a frequency resolution of 2 Hz). This analysis was performed using the ‘signal.welch()’ function from the SciPy Python package. The peak frequency was the frequency with the highest power. The full width at half maximum (FWHM) of the peak was estimated using the “signal.peak_widths()” function from the SciPy Python package.

### Simulation of multipatch recordings

To determine the empirical sampling distance distribution, we used the “Small” database from Campagnola et al. ^30^ and queried for all pairs with experiment type “standard multipatch”, species “mouse”, and “has_probed” value “True”. Following the analysis from that work, layer 5 cells with Cre line *tlx3* were considered to be L5-ITs, while Cre lines *sim1* and *fam84b* were considered to be L5-ET neurons. We fit the distribution of all probed lateral distances *d* with a normal distribution truncated at d=0 by finding the maximum likelihood parameters via the optimize module of Scipy (mu = -29.5 µm; sigma = 92.2 µm).

To determine potential cell pairs, we looked at all cells classified as L5-ET or L5-IT within 300 microns of any of the L5-ET cells with fully proofread axons in V1. Classification used the perisomatic features in Elabaddy et al ^60^, with status as a pyramidal cell confirmed either via spine-density based classifiers where possible or manual validation for cells where not. For each pair, we computed the lateral distance from presynaptic L5-ET cell to the potential target, and assigned it a sampling weight proportional to the empirical lateral distance distribution at that distance. To replicate an experiment where both pre- and postsynaptic cells were labeled, we then performed weighted random sampling with replacement from the list of L5-ET to L5-IT pairs (N=44,526 potential pairs) and from the list of L5-ET to L5-ET pairs (N=6264 pairs) separately, counting the number of synaptically connected pairs from a collection of N random draws (N=82 for L5-IT targets, 739 for L5-ET targets, matching samples from Campagnola et al.). 10,000 sets of random pairs were sampled for each target cell type. Sampling was done using the “random” module of Numpy with the default random number generator.

## ACKNOWLEDGMENTS

The authors thank David Markowitz, the IARPA MICrONS Program Manager, who coordinated this work during all three phases of the MICrONS program. We thank IARPA program managers Jacob Vogelstein and David Markowitz for co-developing the MICrONS program. We thank Jennifer Wang, IARPA SETA for her assistance.

The work was supported by the Intelligence Advanced Research Projects Activity (IARPA) via Department of Interior/Interior Business Center (DoI/IBC) contract numbers D16PC00003, D16PC00004, and D16PC0005. The U.S. Government is authorized to reproduce and distribute reprints for Governmental purposes notwithstanding any copyright annotation thereon. HSS also acknowledges support from NIH/NINDS U19 NS104648, NIH/NEI R01 EY027036, NIH/NIMH U01 MH114824, NIH/NIMH U01 MH117072 NIH/NINDS R01 NS104926, NIH/NIMH RF1 MH117815, NIH/NIMH RF1 MH123400 and the Mathers Foundation, as well as assistance from Google, Amazon, and Intel. N. M. C. also acknowledges support from NIH 1RF1MH128840-01 and NSF NeuroNex 2 award 2014862. AK acknowledges support by the National Institute of Biomedical Imaging and Bioengineering of the National Institutes of Health under Award Number R01EB029813, by the National Institute of Neurological Disorders and Stroke of the National Institutes of Health under Award Numbers R01NS122742 and U24NS124001, and by the National Institute Of Mental Health of the National Institutes of Health under Award Number U01MH130907. The content is solely the responsibility of the authors and does not necessarily represent the official views of the National Institutes of Health.

We thank Rob Young for managing the stitching and alignment pipeline at the Allen Institute for Brain Science (AIBS). We thank John Philips, Sill Coulter and the Program Management team at the AIBS for their guidance for project strategy and operations. We thank Hongkui Zeng, Ed Lein, Christof Koch and Allan Jones for their support and leadership. We thank the Manufacturing and Processing Engineering team at the AIBS for their help in implementing the EM imaging and sectioning pipeline. We thank Brian Youngstrom, Stuart Kendrick and the Allen Institute IT team for support with infrastructure, data management and data transfer. We thank the Facilities, Finance, and Legal teams at the AIBS for their support on the MICrONS contract. We thank Stephan Saalfeld, Khaled Khairy and Eric Trautman for help with the parameters for 2D stitching and rough alignment of the dataset. We thank Staci Sorensen for discussions on cell types.

We thank Garrett McGrath for computer system administration, and May Husseini and Larry and Janet Jackel for project administration at Princeton University.

We thank Brock Wester, William Gray-Roncal, Sandy Hider, Tim Gion, Daniel Xenes, Jordan Matelsky, Caitlyn Bishop, Derek Pryor, Dean Kleissas, Luis Rodriguez and Miller Wilt from John Hopkins University Applied Physics lab for providing data assessments on the neural circuit reconstruction and infrastructure through BOSSdb

We thank Frances Chance, Brad Aimone, and everyone at Sandia National Laboratories for their support and assistance.

We would like to thank the “Connectomics at Google” team for developing Neuroglancer and computational resource donations, in particular J. Maitin-Shepard for authoring neuroglancer and help creating the reformatted sharded multi-resolution meshes and imagery files used to display the data. We would like to thank Amazon and the AWS Open Science platform for providing computational resources. We’d like to also thank Intel for their assistance.

We thank the Allen Institute for Brain Science founder, Paul G. Allen, for his vision, encouragement and support. Disclaimer: The views and conclusions contained herein are those of the authors and should not be interpreted as necessarily representing the official policies or endorsements, either expressed or implied, of IARPA, DoI/IBC, or the U.S. Government.

