## Supplemental Figures for "The Synaptic Architecture of Layer 5 Thick Tufted Excitatory Neurons in the Visual Cortex of Mice"

### SUPPLEMENTAL INFORMATION

#### Extended Figure 1

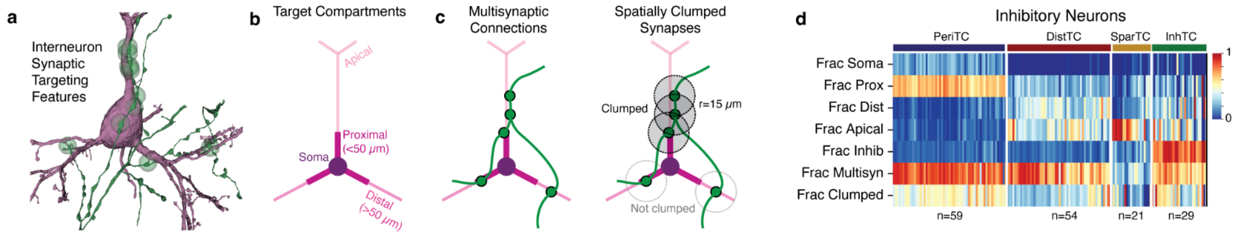


Extended Fig 1. Adapted from Figure 2 of Schneider-Mizell et. al (Schneider-Mizell et al. 2023). Data-driven characterization of inhibitory cell subclasses. **a.** For determining inhibitory cell subclasses, we used the high resolution neuroanatomical properties of axons (green), synapses (green spheres), and their postsynaptic target dendrites and soma (purple) to measure properties of synaptic output of inhibitory neurons. The example depicts a basket cell axon and pyramidal cell soma and proximal dendrites. **b.** Dendritic compartment definitions for excitatory neurons. For each excitatory neuron compartment, we measured the fraction of inputs (among outputs onto excitatory neurons only). Synapses onto inhibitory neurons were considered as a fifth compartment. **c.** To describe the anatomical properties of typical connections, we measured two properties: the fraction of all synapses that were part of multisynaptic connections (left) and the fraction of synapses in multisynaptic connections onto the same target that were close together (<15 microns) along the path of the axon. **d.** Heatmap of feature properties versus class labels based on using linear discriminant analysis to classify neurons into four subclasses from on a subset of manual labels.

Extended Figure 2

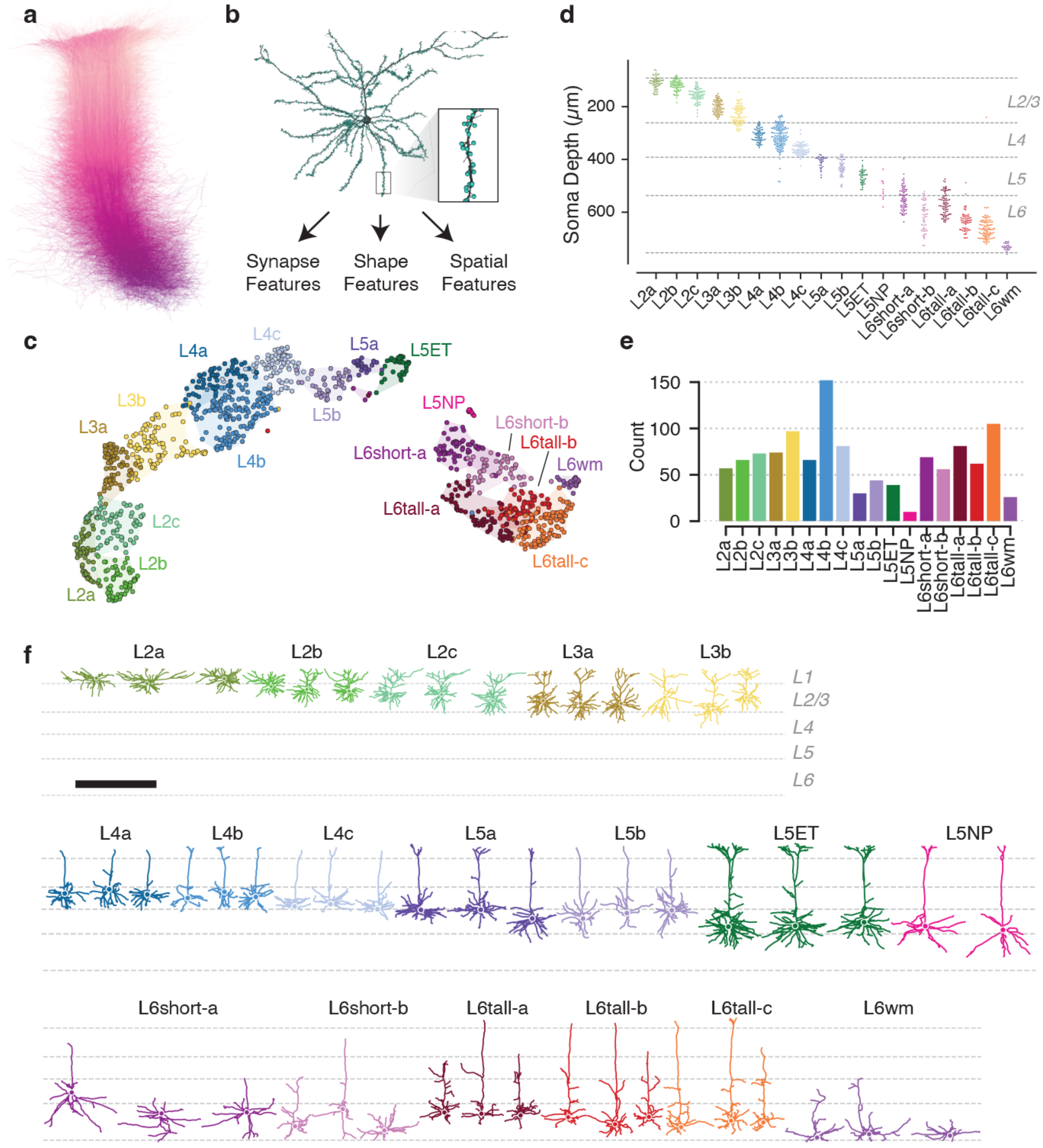


Extended Fig 2. Adapted from Figure 1 and 3 of Schneider-Mizell et. al (Schneider-Mizell et al. 2023).

Data-driven characterization of excitatory cell subclasses. **a.** Excitatory neuron dendrites were reconstructed across all layers of visual cortex. **b, c.** A collection of morphological, synaptic, and spatial features were computed for each neuron (**b**) and used to cluster cells into anatomical categories (**c**). Clustering results were computed in the full feature space and visualized with

UMAP. **d**. Soma depth distribution for each cluster. **e**. Number of cells of each type along the column (N=1188 cells in total). **f.** Anatomical examples of each excitatory class.

Extended Figure 3

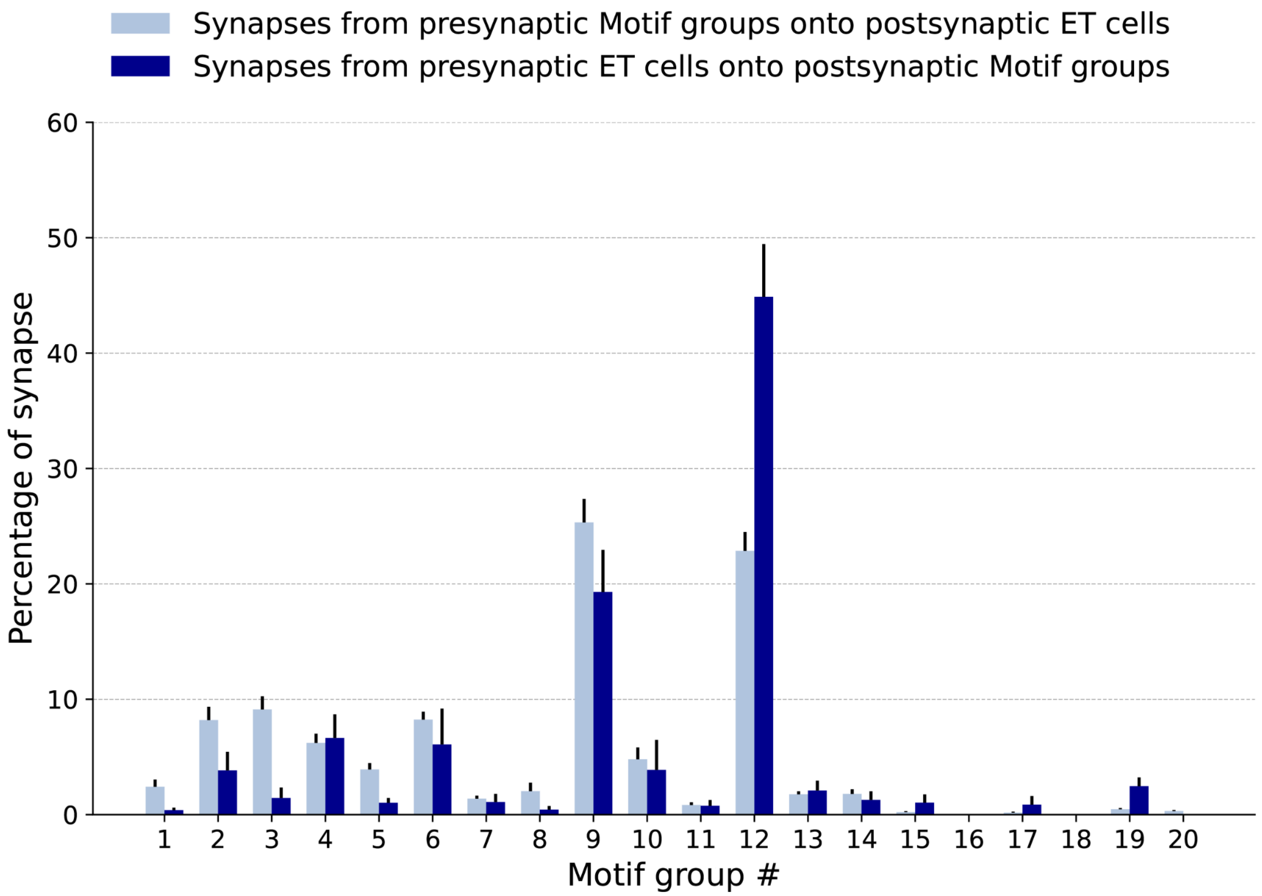


**Extended Data Figure 3.** Histogram showing that the ET neurons from the column in Figure2.a target the inhibitory cells in return preferentially inhibit the ET neurons. This is an extension of Figure 2.g, and instead of having the inhibitory cell divided into just 2 groups they are divided according to the 20 motifs described in the accompanying manuscript of Schneider-Mizell et al.[59] The dark blue histogram shows the distribution of synapses from the presynaptic ET neurons onto the different motif groups. The light blue histogram shows the distribution of synapses from the presynaptic motif groups neurons onto ET neurons. Data is shown as mean and SEM.

Extended Figure 4

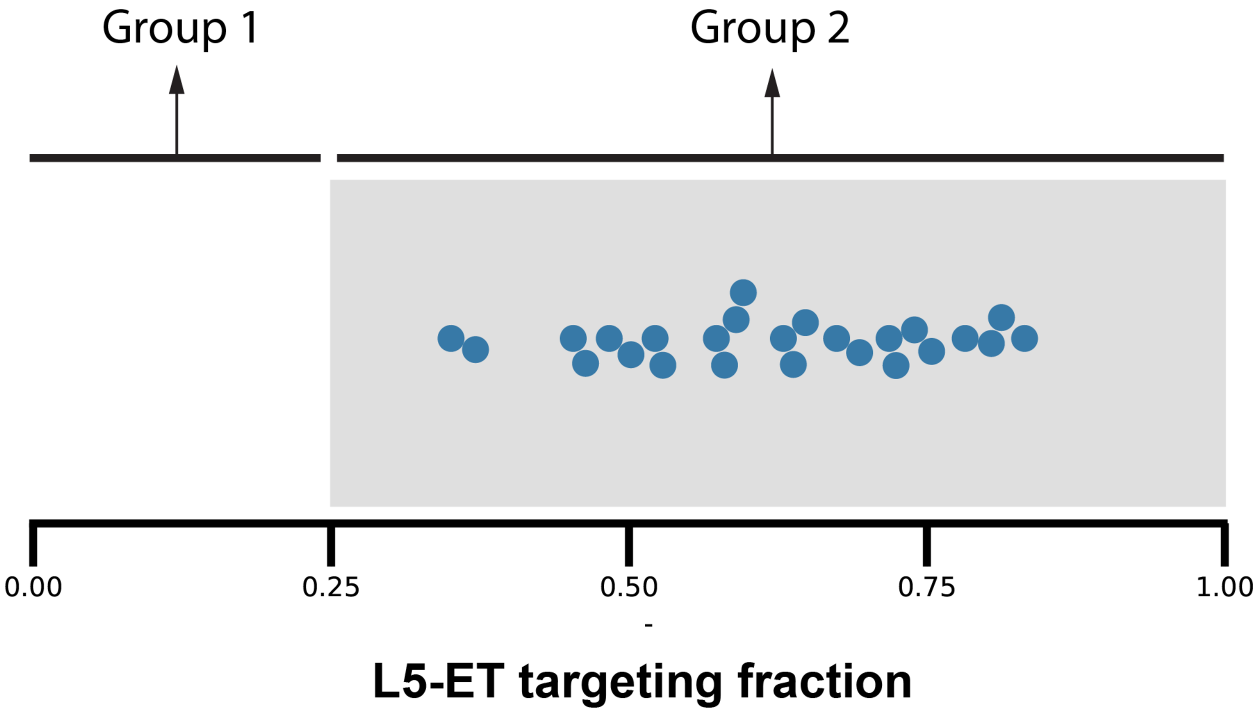


**Extended Figure 4.** Swarm plots showing the fraction of synapses formed by a selection of perisomatic targeting cells onto postsynaptic ET neurons. In this plot, the inhibitory cells chosen for analysis and proofreading were the ones that received the highest number of synapses from the 12 proofread ET neurons shown in Extended Figure 5.

#### Extended Figure 5

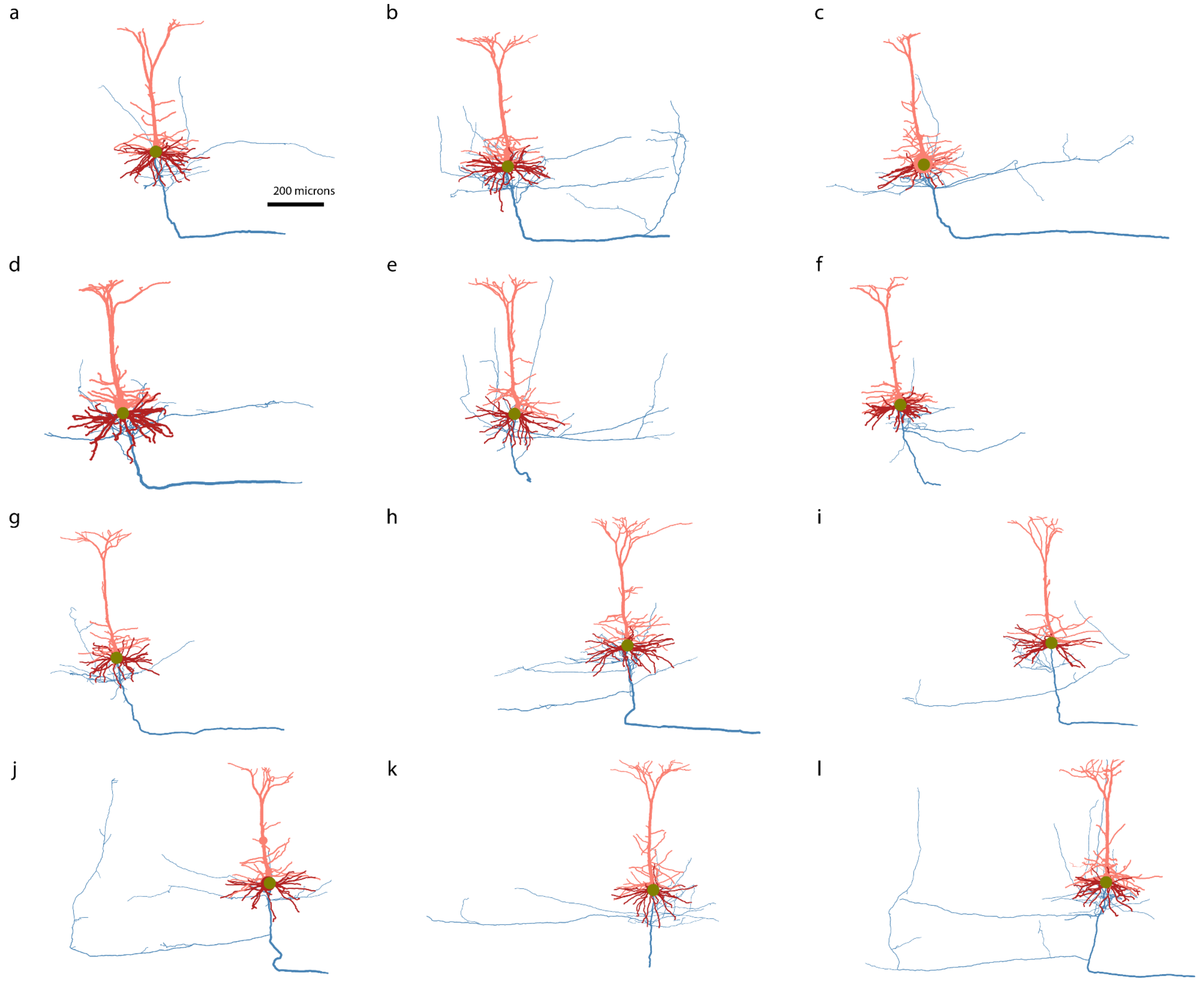


**Extended Figure 5. Morphology of ET neurons.** Panels **a-l** show the gross morphology of the dendrites and axons of the 12 ET neurons used for the analysis in Figures 3 and 4. The L5 ET axons display a distinct pattern of myelination along their main vertical branch, with gaps occurring approximately every 3-5 times. At these points, 1-4 branches emerge, which are unmyelinated initially but may acquire myelin further down their length. The upper two branch points typically give rise to local branches that primarily remain in the vicinity of the dendritic area, mainly in layers 4, 5, and 6. While some rare branches may extend into upper layers, they tend to stay within the width of the dendritic arbors. The lower branch points of the main axon generate the segments with inter-areal projections, which exhibit diverse morphologies. Some of these branches may extend over long distances, with segments being myelinated or partially myelinated along the way, only to arborize again unmyelinated at their target location. These long-range axons are typically sparse in terms of forming synapses and show more variable targeting profile, likely due to the proximity to the edges of the dataset. The main branch of the L5 ET axon remains myelinated as it enters the white matter.

Extended Figure 6

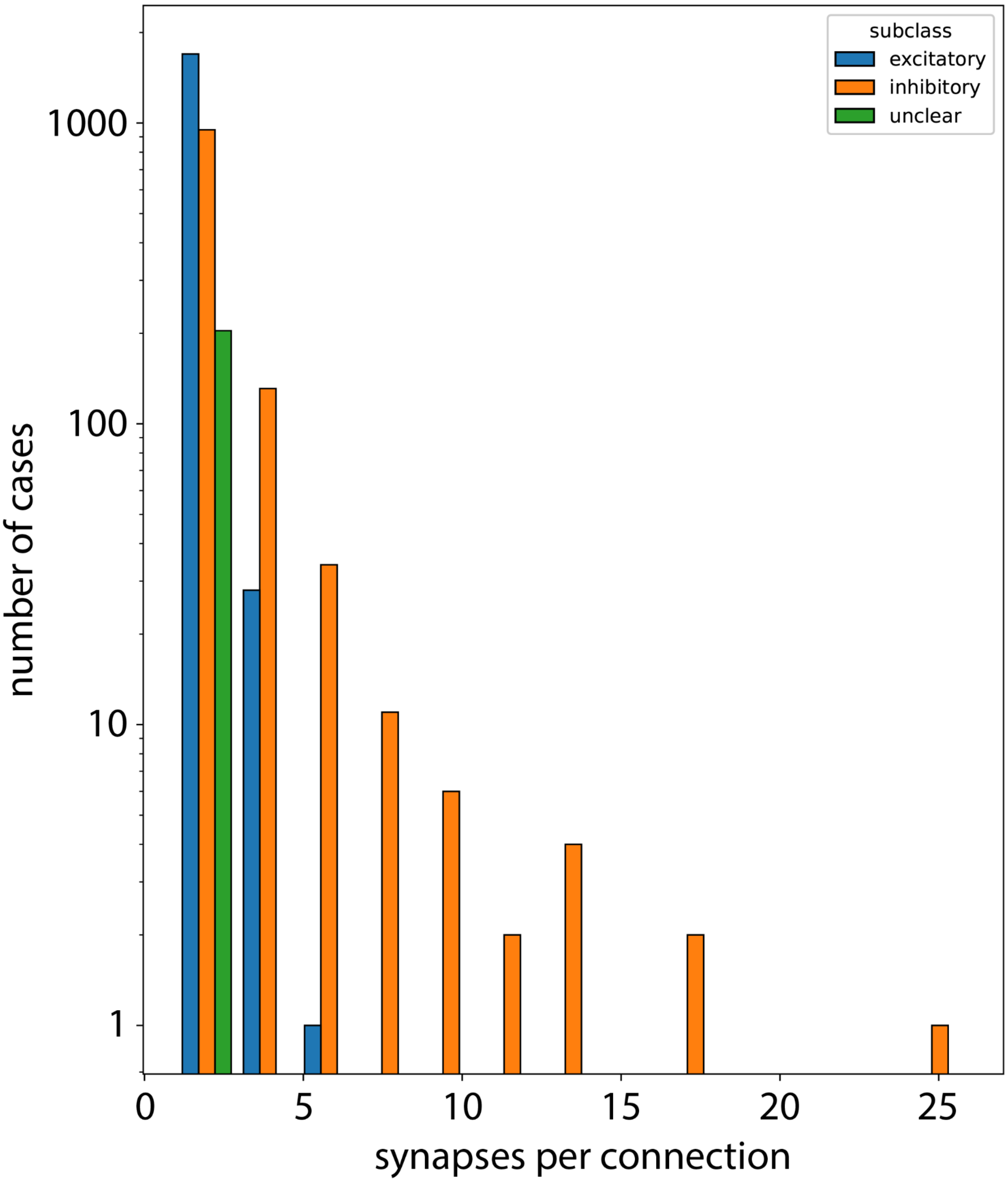


**Extended Figure 6**: Number of synapses per connection arranged by cell class

### Extended Figure 7

| | L5-ET | L5-IT | L5-NP | L6-P | PTC | DTC |
|---|---|---|---|---|---|---|
| L5-ET | 1.0 | 1.0 | 1.0 | 8.1e-01 | 1.0 | 1.0 |
| L5-IT | 1.0 | 1.0 | 0.9 | 1.7e-04 | 1.1e-01 | 1.9e-01 |
| L5-NP | 1.0 | 0.9 | 1.0 | 4.1e-05 | 1.0 | 1.0 |
| L6-P | 0.8 | 1.7e-04 | 4.1e-05 | 1.0 | 6.7e-13 | 1.0e-10 |
| PTC | 1.0 | 0.1 | 1.0 | 6.7e-13 | 1.0 | 1.0 |
| DTC | 1.0 | 0.2 | 1.0 | 1.0e-10 | 1.0 | 1.0 |

**Extended Figure 7**: Results of Conover's post hoc pairwise test for multiple comparisons.

Extended Figure 8

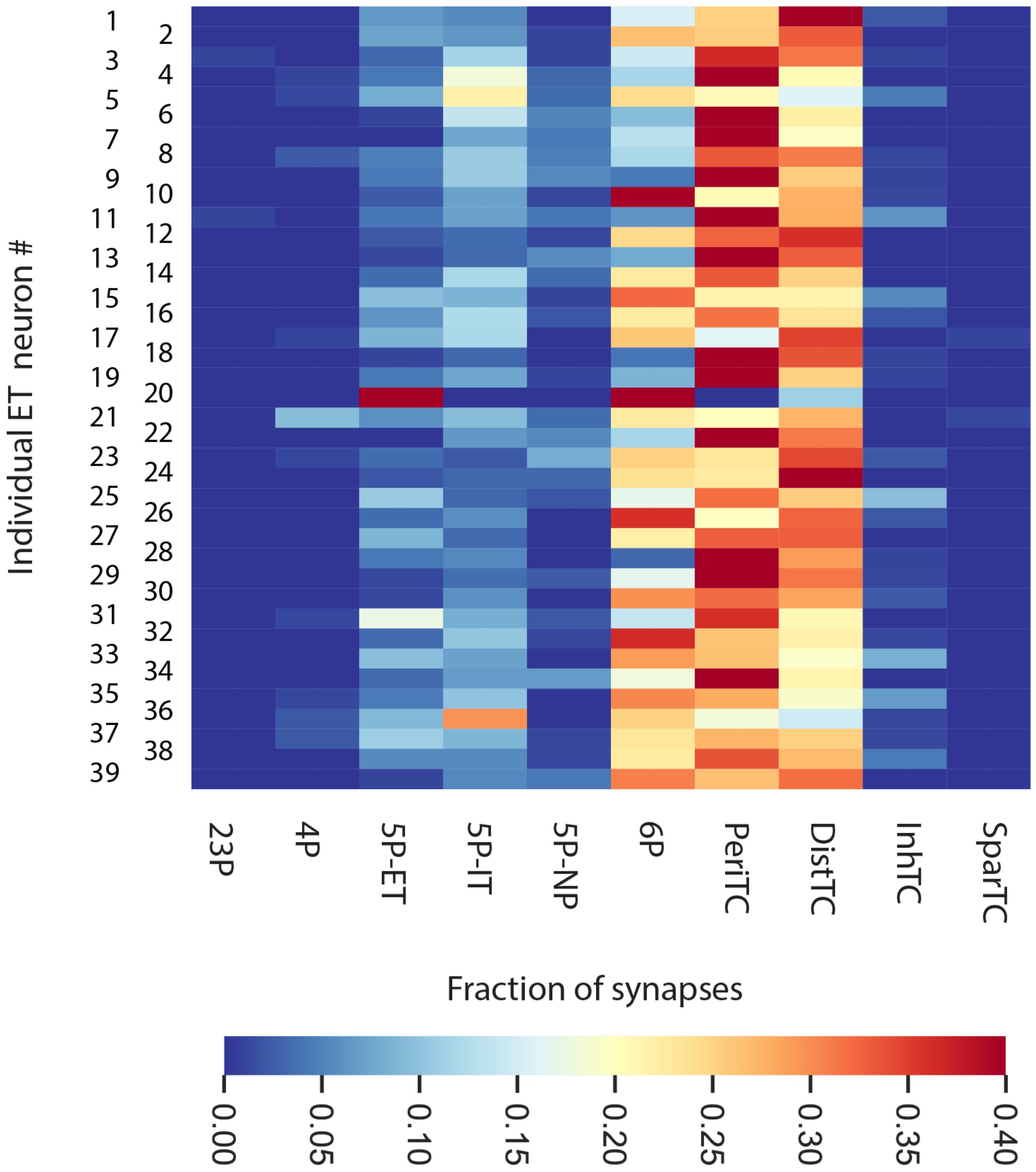


**Extended Figure 8.** Heat plots of the output connectivity of the column 39 ET cells with model

The heat map illustrates the fraction of 100 proximal synapses formed by ET neurons in the column of Figure 2 with target neurons categorized by cell sub-class. Rows correspond to individual neurons.

### Extended Figure 9

To investigate how the sparsity of excitatory targets affect the activity of the circuit, we built a L5 circuit model of generalized leaky-integrate-and-fire (GLIF) neurons, consisting of L5 ET cells, L5 Pvalb cells (Basket Cells), and L5 Sst cells (Martinotti Cells, Extended Fig 9.a and 9.b). This model was based upon an earlier model of the mouse primary visual cortex (Billeh et al., 2020), from which we extracted the neuronal models for L5, modifying the connectivity according to experimental findings reported here (see Methods). Here we specifically adjusted the connection probabilities of L5 ET cells based on the 12 proofread L5 ET neurons. To simulate the response of the circuit to external input, the model received a Poisson spike train at 1 kHz. Neurons were assigned a 'preferred angle' spanning from 0 to 360 degrees, i.e., forming a ring in this abstract preferred angle space, and the weight of the Poisson input to each neuron was determined by a Gaussian function on this ring. To investigate how the connection pattern affects the activity, we compare the performance of the model with recurrent connections: neurons receive external inputs, plus the synaptic inputs from the 3 cell types in the circuit, to that of the model without recurrent connections: neurons only receive external inputs, and they are independent from other neurons in the circuit. We found that the circuit with the recurrent connections shows a narrowing effect in the firing rate as a function of the preferred angle, which suggests that the tuning to the incoming stimulation is sharpened by the recurrent connections in this L5 circuit (Extended Fig 9.c and 9.d). Notably, the sharpening is weaker when the fraction of L5 ET targets increases (Extended Fig 9.ee), indicating that sparse excitatory and extensive inhibitory connectivity in the circuit favor sharpening of the circuit output relative to the input. In addition, L5 ET cells in the circuit exhibit gamma-range oscillations in their firing rates, whereas in the absence of recurrent connections, the peak frequency is observed at a lower frequency (Extended Fig 9.f and 9.g) The peak shifts again to a lower frequency when the fraction of L5 ET targets increases (Extended Fig 9.h). These findings indicate that sparse L5 ET targets and extensive innervation of inhibitory cells by L5 ET cells, observed in the EM data, may support sharpening in processing of stimuli, as well as gamma-frequency firing in L5.

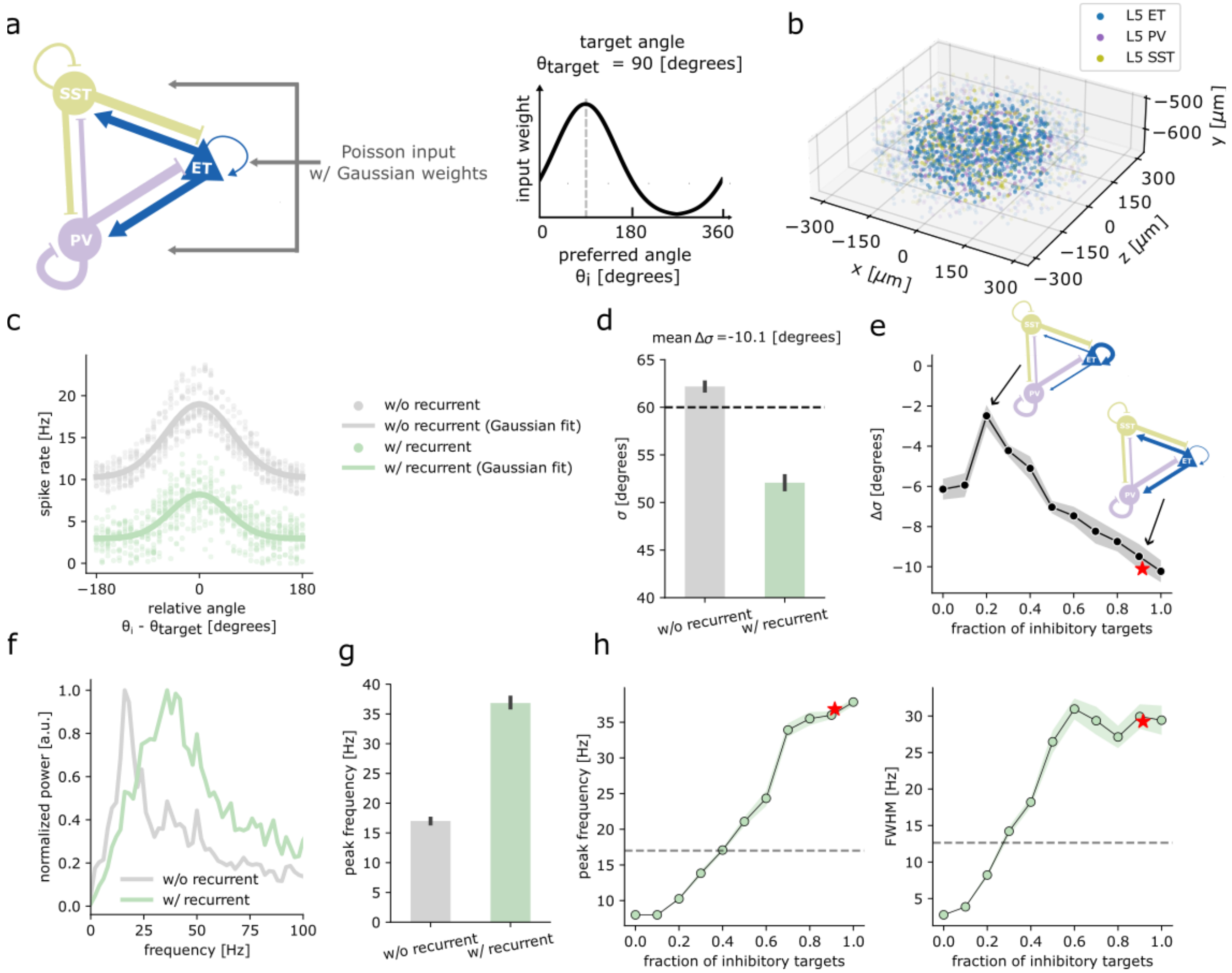


**Extended Figure 9. A L5 circuit model of GLIF neurons.** a) Left, a cartoon illustration of the L5 circuit model. The lines represent the recurrent connections, where the thickness roughly corresponds to the connection probability. The connection probabilities from L5 ET (blue lines) are adjusted based on the 12 proofread ET cells, whereas the others (purple and olive lines) are from the Synaptic Physiology dataset (Campagnola et al. 2022). Right, the weight of the Poisson input to each neuron, as a function of the neuron's preferred angle relative to the target angle. This example shows a trial with a target angle of 90 degrees, where neurons with a preferred angle of 90 degrees received the strongest input. Please note that the illustration does not depict the background inputs that neurons also receive. b) Visualization of the model in the cylindrical domain. Each dot represents the location of an individual neuron. The neurons within the core (radius<200) are highlighted. The radius of the cylinder is 650 um, but only the domain with radius<300 um is shown here for better visualization. Edges are not shown in this visualization. c) The firing rates of L5 ET cells receiving a Poisson input with Gaussian weights in the model with recurrent connections (as shown in panel (a)) or without recurrent connections (similar to panel (a) but without the blue, purple, and olive lines), for an example simulation. Each dot represents the firing rate of an individual neuron. The x axis is the preferred angle of each neuron relative to the target angle that receives the strongest Poisson input in a given simulation trial. The lines are the Gaussian fits of the firing rates for each condition. d) Standard deviation of the Gaussian fit for the model with or without recurrent connections. The error bars are +/- 1 standard error from simulations targeting different angles. The dotted line represents the standard deviation of the Gaussian weights of the Poisson input. e) Difference between the standard deviations of the Gaussian fit for the model with recurrent connections and the one without recurrent connections. Each dot corresponds to a model with a specific fraction of L5 inhibitory neurons among the L5 ET synaptic targets in this circuit (L5 ET, L5 Pvalb, and L5 Sst), ranging from 0 to 1. The connectivity from L5 Pvalb and L5 Sst are not changed. The two inserted cartoons illustrate the model configuration when the fraction is 0.2 (20% of the connections from L5 ET in the circuit target L5 Pvalb and Sst and 80% of them target L5 ET), and the model configuration when the fraction is observed from the 12 proofread ET cells (red star, same connectivity as shown in panel (a)). See how the thickness of the blue lines is different. The shaded area is +/- 1 standard error from simulations targeting different angles. f) Normalized power spectrum of the L5 ET firing rate in the model with or without recurrent connections, for an example simulation (same model as in panels (a), (b), (c) and (d)). The power spectrum is normalized by the peak value. g) Frequency of the peak of the power spectrum for the model with or without recurrent connections. The error bars are +/- 1 standard error from simulations targeting different angles (same model as in panel (a), (b), (c) and (d)). h) Left, peak frequency of the model with recurrent connections. Right, full width at half maximum (FWHM) of the peak of power spectrum of the model with recurrent connections. The dotted lines represent the results of the model without recurrent connections. The dots and the stars represent the same models as in panel (e).

### Extended Figure 10

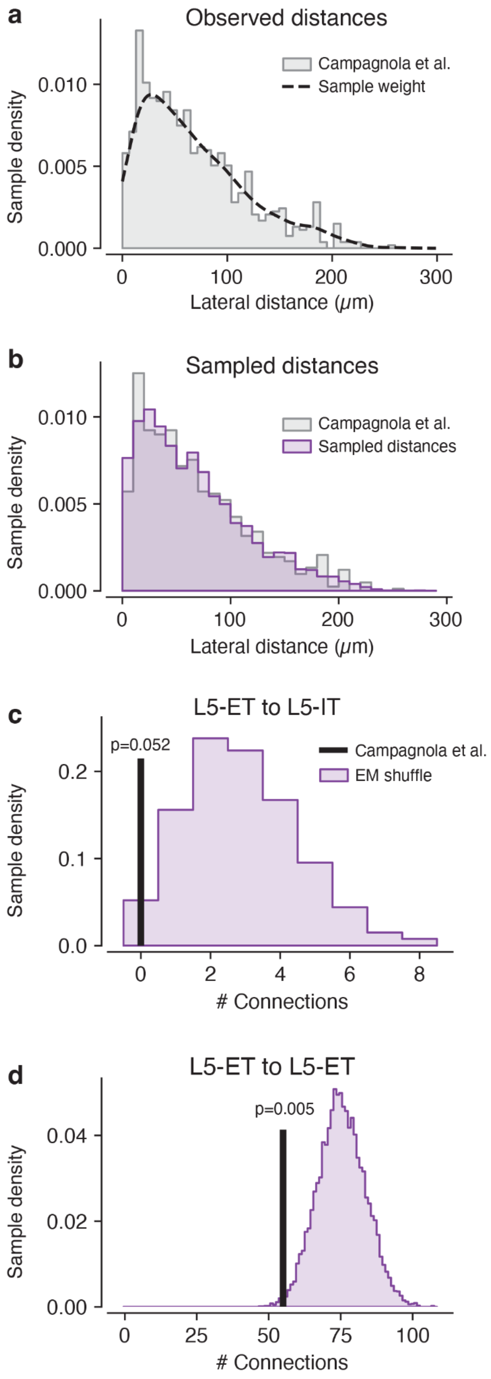


**Extended Figure 10. Simulation of synapse physiology experiments within an EM dataset. a.** Histogram showing the distribution of samples distance in the synapse physiology experiments of Campagnola and colleagues [30]. **b.** Histogram comparing the distribution of sample distances in the original synapse physiology data (gray) and in the simulated experiment in the EM volume (purple). **c.** Histogram showing the results of 10,000 random draws of L5-ET to L5-IT connections, weighted by the distribution of sample distance shown in (b). Each random draw contained the same number of pairs as in Campagnola et al (N=82 for L5-IT targets) and counted the number of synaptically connected pairs observed per draw. The black vertical line represents the result of the original synapse physiology experiment. **d.** Histogram showing the results of 10,000 random draws of L5-ET to L5-ET connections, weighted by the distribution of sample distance shown in (b). Each random draw contained the same number of pairs as in Campagnola et al (N=739 for L5-ET targets) and counted the number of synaptically connected pairs observed per draw. The black vertical line represents the result of the original synapse physiology experiment.

### Extended Figure 11

**Extended Figure 11:** Skeletons of the dendrites of 1,638 excitatory neurons postsynaptic to the 12 layer 5 ET neurons shown in Extended Figure 4. The neurons are ordered according to the soma depth. Layer 3 pyramidal cells are shown in green, Layer 4 pyramidal cells are shown in orange, Layer 5 IT pyramidal cells are shown in purple, Layer 5 ET neurons are shown in pink, Layer 5 NP neurons are shown green and layer 6 pyramidal cells are shown in yellow. Below each neuron there is a unique nucleus ID that can be used to visualize the neuron in the publicly available repository https://www.microns-explorer.org/cortical-mm3 using the 'Launch app' link.

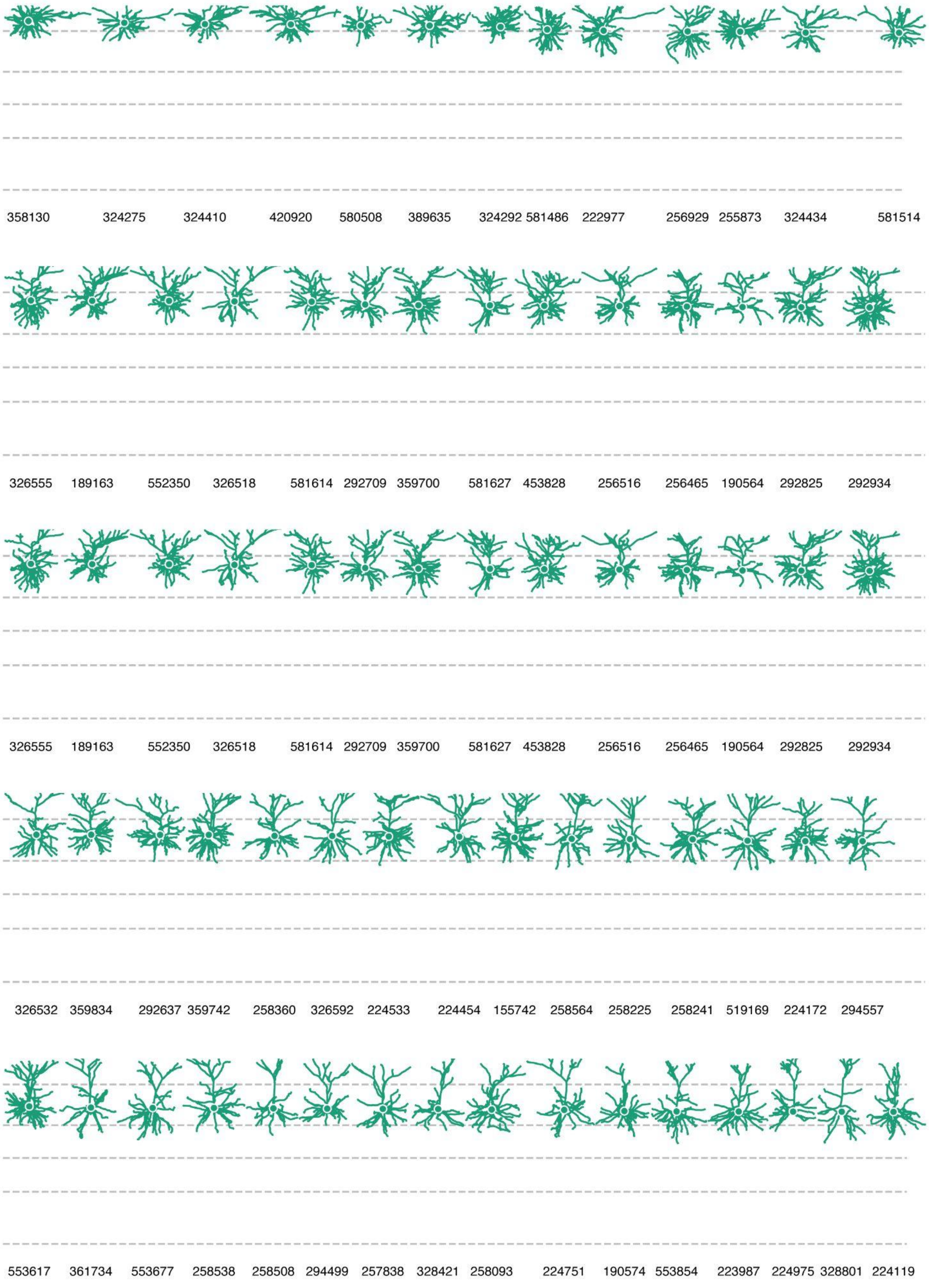
358130 324275 324410 420920 580508 389635 324292 581486 222977 256929 255873 324434 581514
326555 189163 552350 326518 581614 292709 359700 581627 453828 256516 256465 190564 292825 292934
326555 189163 552350 326518 581614 292709 359700 581627 453828 256516 256465 190564 292825 292934
326532 359834 292637 359742 258360 326592 224533 224454 155742 258564 258225 258241 519169 224172 294557
553617 361734 553677 258538 258508 294499 257838 328421 258093 224751 190574 553854 223987 224975 328801 224119

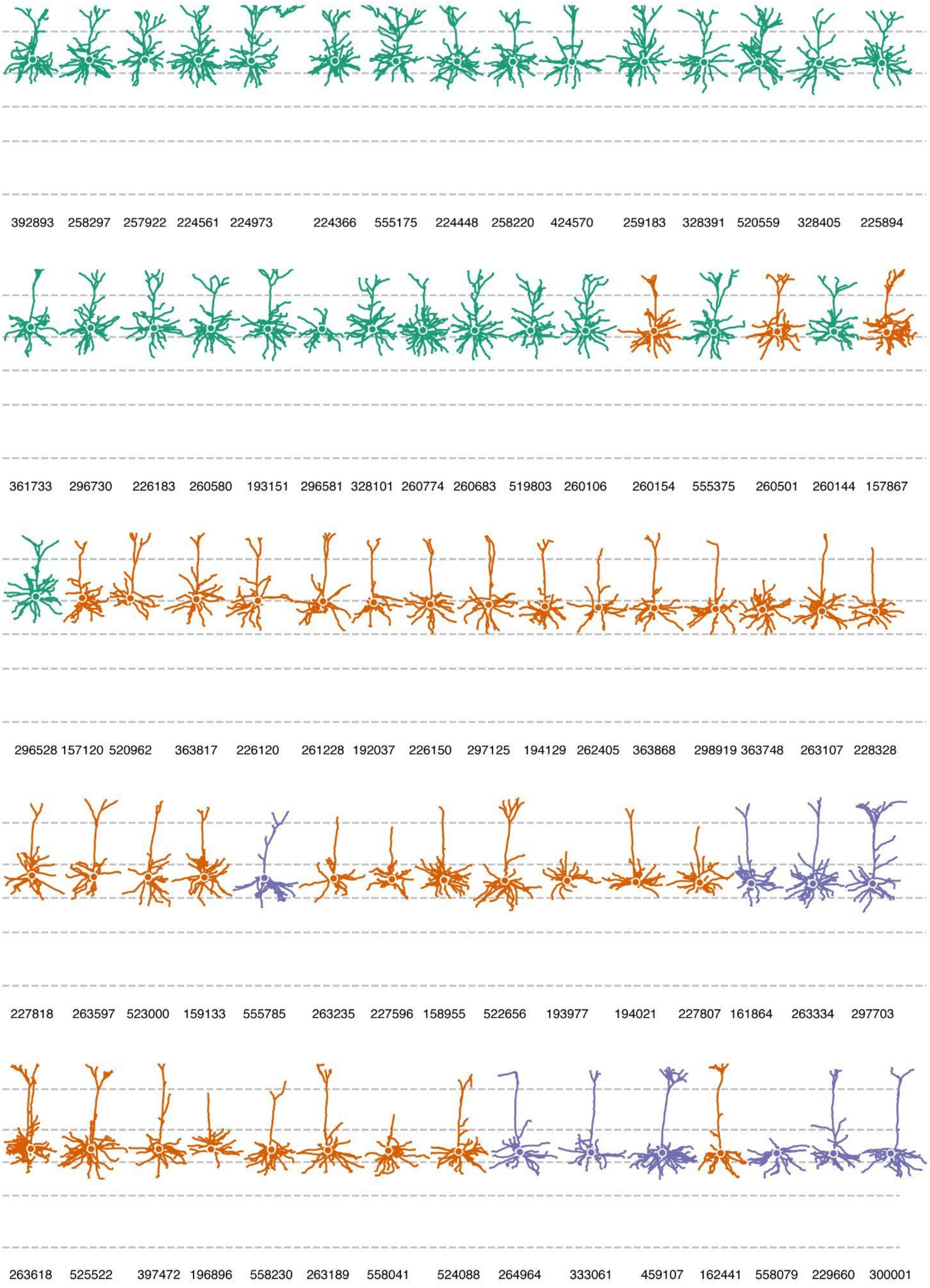
392893 258297 257922 224561 224973 224366 555175 224448 258220 424570 259183 328391 520559 328405 225894
361733 296730 226183 260580 193151 296581 328101 260774 260683 519803 260106 260154 555375 260501 260144 157867
296528 157120 520962 363817 226120 261228 192037 226150 297125 194129 262405 363868 298919 363748 263107 228328
227818 263597 523000 159133 555785 263235 227596 158955 522656 193977 194021 227807 161864 263334 297703
263618 525522 397472 196896 558230 263189 558041 524088 264964 333061 459107 162441 558079 229660 300001

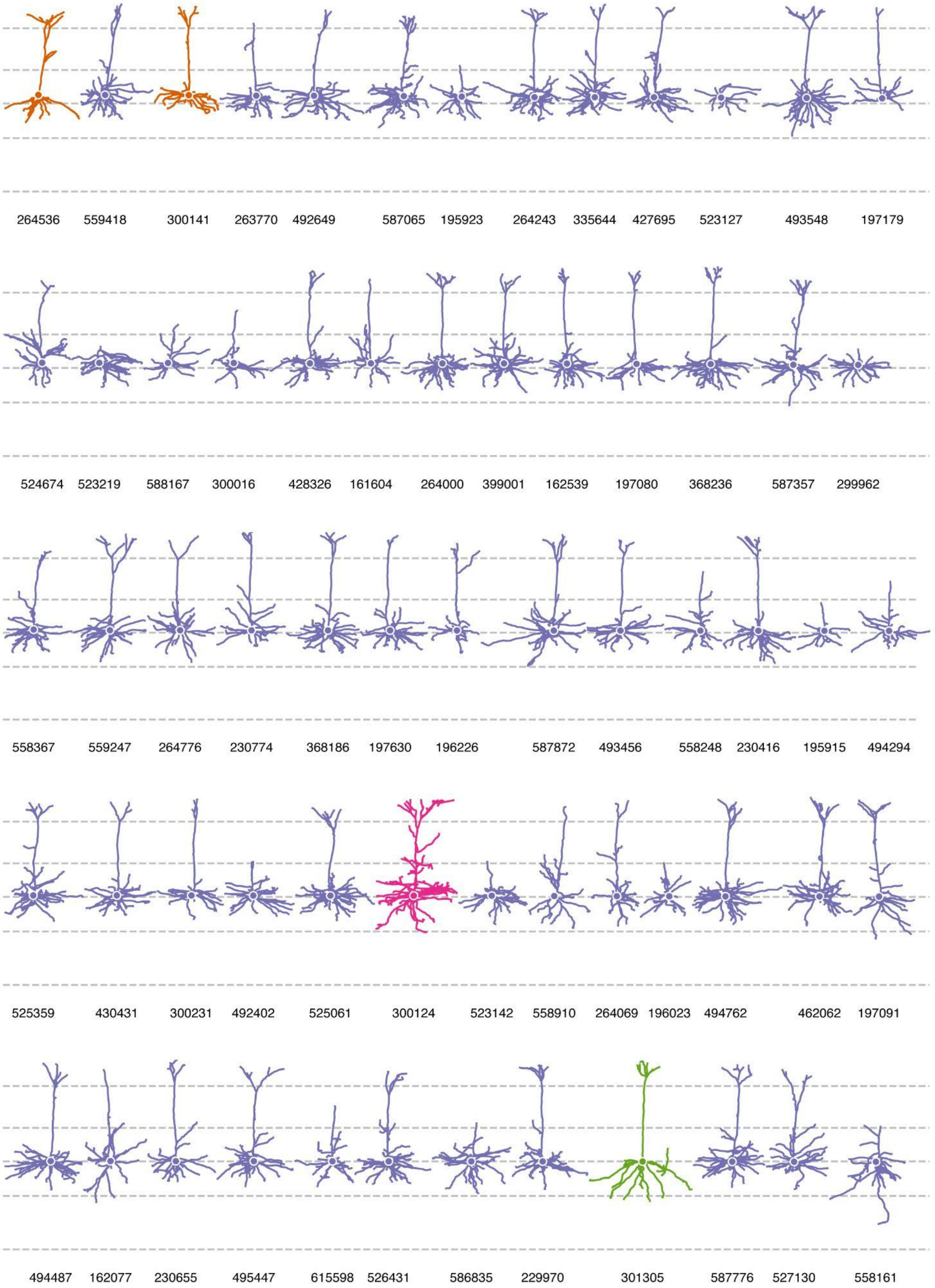
264536 559418 300141 263770 492649 587065 195923 264243 335644 427695 523127 493548 197179
524674 523219 588167 300016 428326 161604 264000 399001 162539 197080 368236 587357 299962
558367 559247 264776 230774 368186 197630 196226 587872 493456 558248 230416 195915 494294
525359 430431 300231 492402 525061 300124 523142 558910 264069 196023 494762 462062 197091
494487 162077 230655 495447 615598 526431 586835 229970 301305 587776 527130 558161

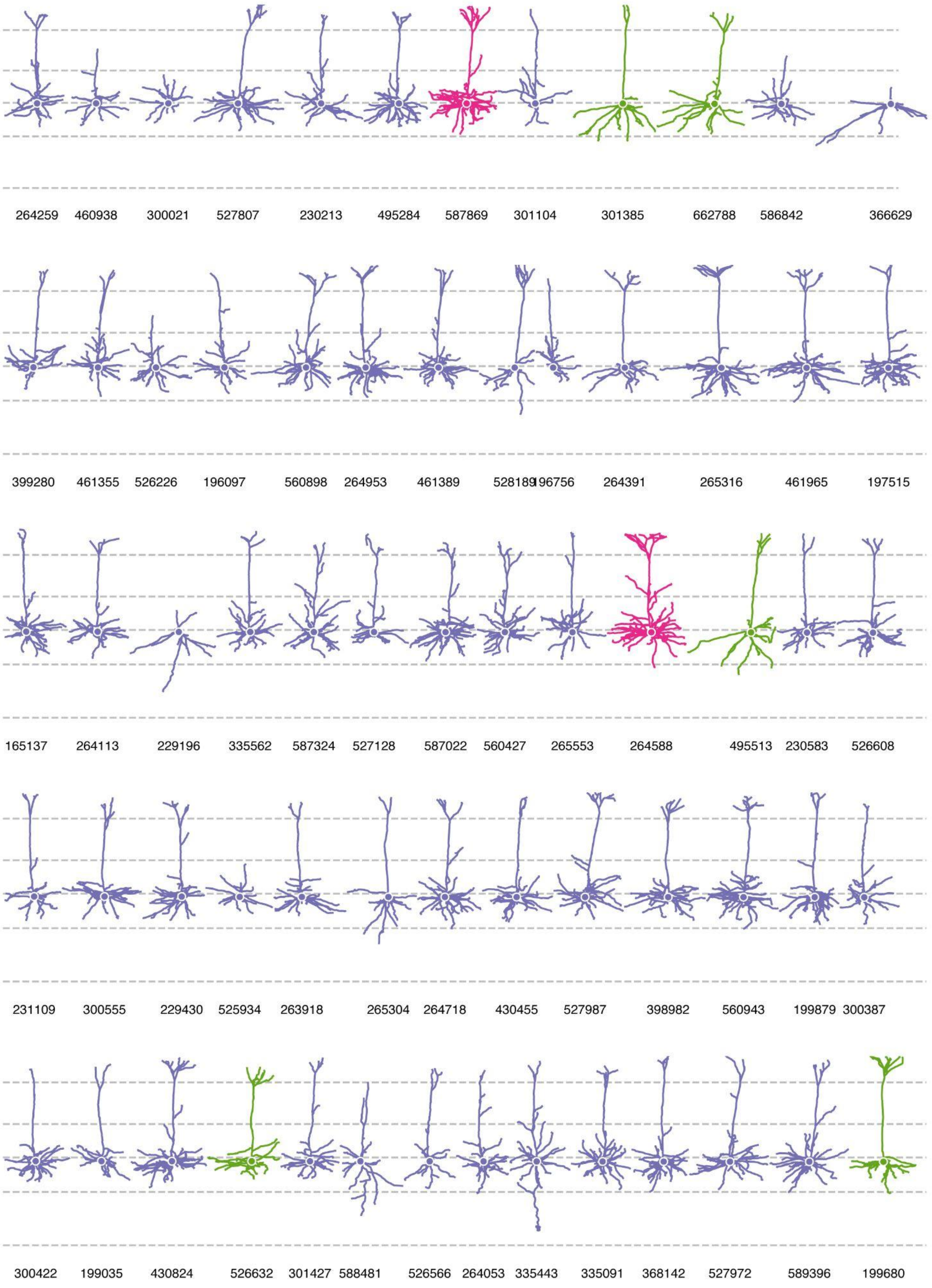
264259 460938 300021 527807 230213 495284 587869 301104 301385 662788 586842 366629
399280 461355 526226 196097 560898 264953 461389 528189 196756 264391 265316 461965 197515
165137 264113 229196 335562 587324 527128 587022 560427 265553 264588 495513 230583 526608
231109 300555 229430 525934 263918 265304 264718 430455 527987 398982 560943 199879 300387
300422 199035 430824 526632 301427 588481 526566 264053 335443 335091 368142 527972 589396 199680

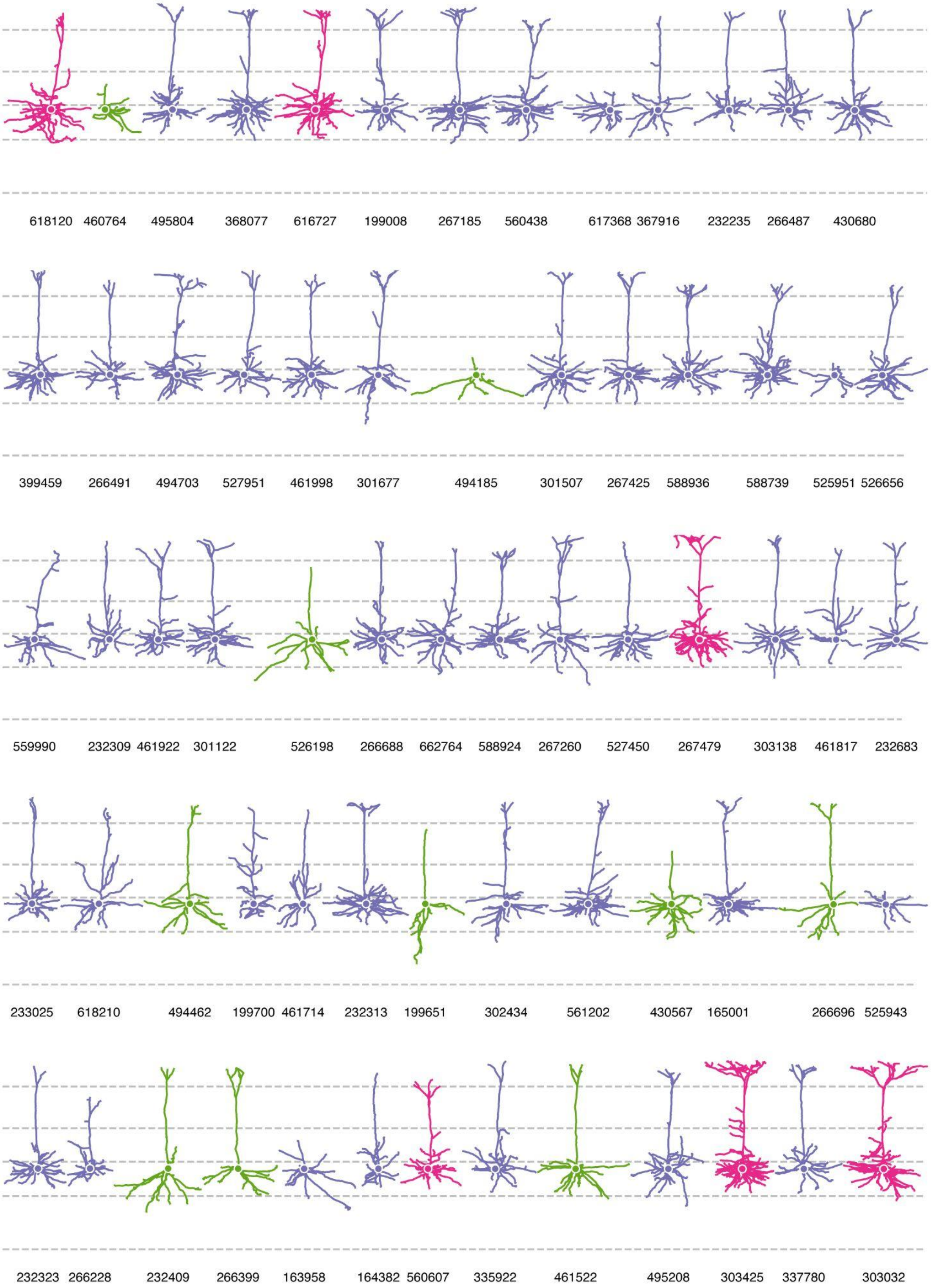
618120 460764 495804 368077 616727 199008 267185 560438 617368 367916 232235 266487 430680
399459 266491 494703 527951 461998 301677 494185 301507 267425 588936 588739 525951 526656
559990 232309 461922 301122 526198 266688 662764 588924 267260 527450 267479 303138 461817 232683
233025 618210 494462 199700 461714 232313 199651 302434 561202 430567 165001 266696 525943
232323 266228 232409 266399 163958 164382 560607 335922 461522 495208 303425 337780 303032

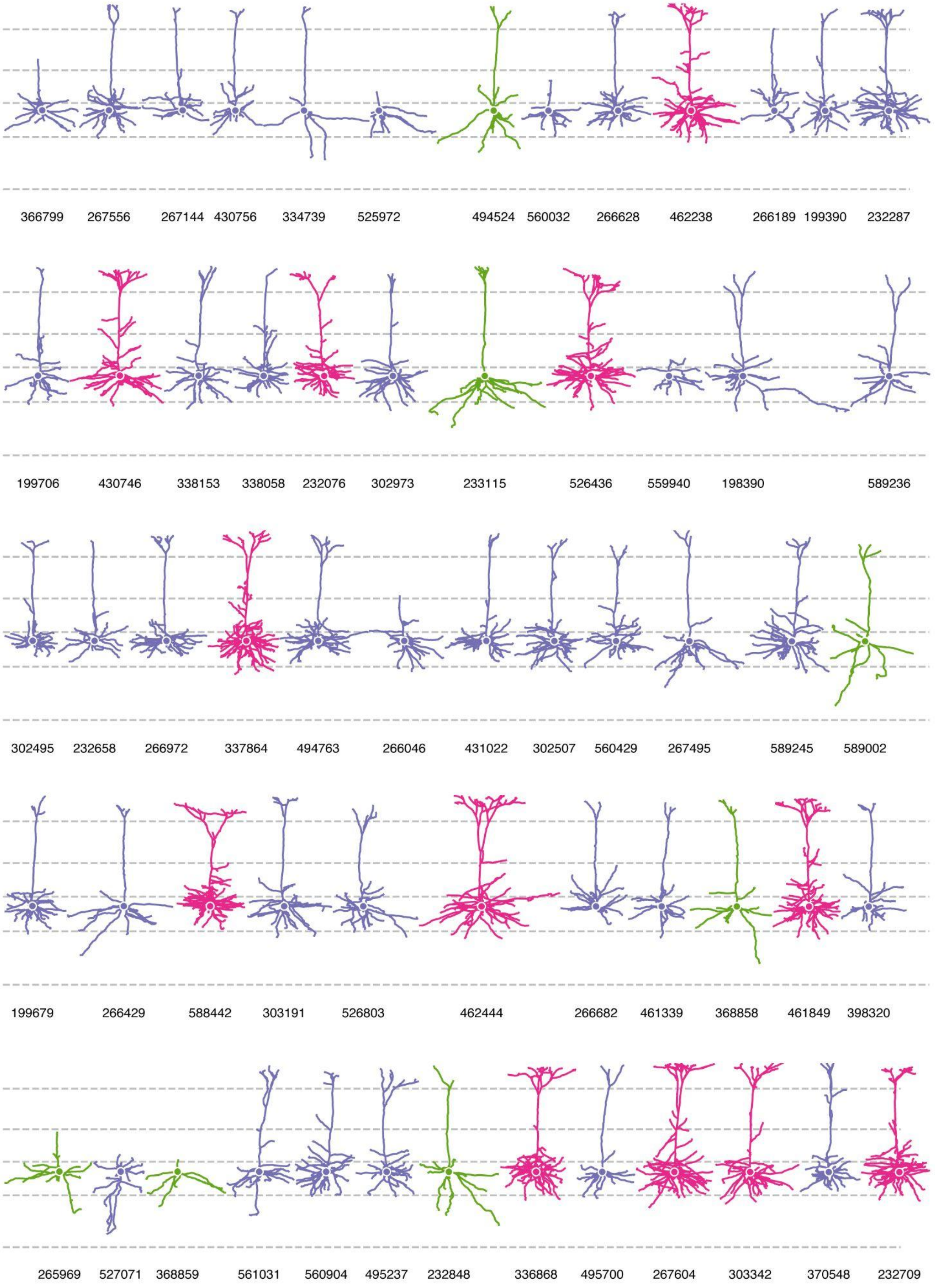
366799 267556 267144 430756 334739 525972 494524 560032 266628 462238 266189 199390 232287
199706 430746 338153 338058 232076 302973 233115 526436 559940 198390 589236
302495 232658 266972 337864 494763 266046 431022 302507 560429 267495 589245 589002
199679 266429 588442 303191 526803 462444 266682 461339 368858 461849 398320
265969 527071 368859 561031 560904 495237 232848 336868 495700 267604 303342 370548 232709

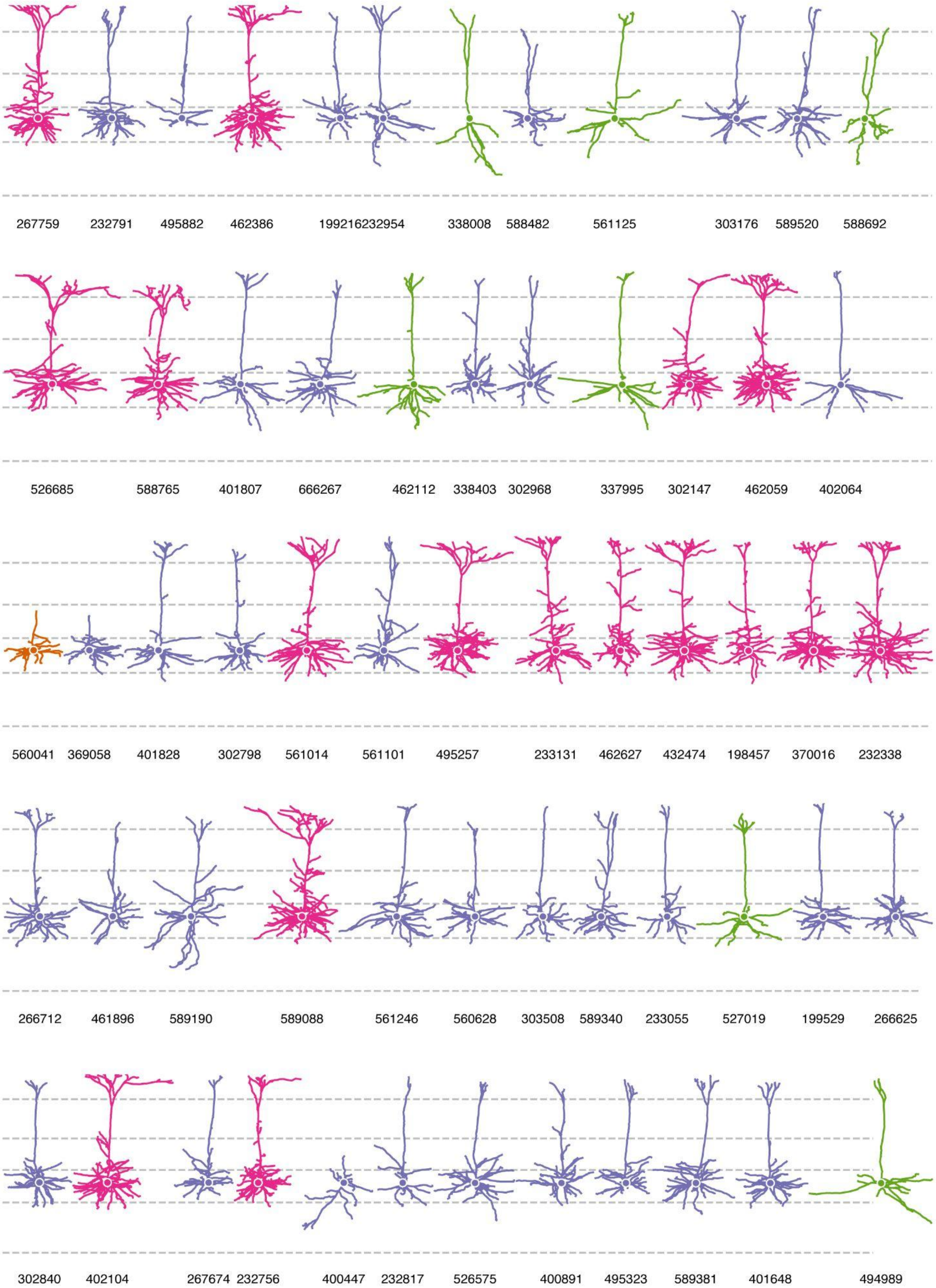

267759
232791
495882
462386
199216232954
338008
588482
561125
303176
589520
588692
526685
588765
401807
666267
462112
338403
302968
337995
302147
462059
402064
560041
369058
401828
302798
561014
561101
495257
233131
462627
432474
198457
370016
232338
266712
461896
589190
589088
561246
560628
303508
589340
233055
527019
199529
266625
302840
402104
267674
232756
400447
232817
526575
400891
495323
589381
401648
494989

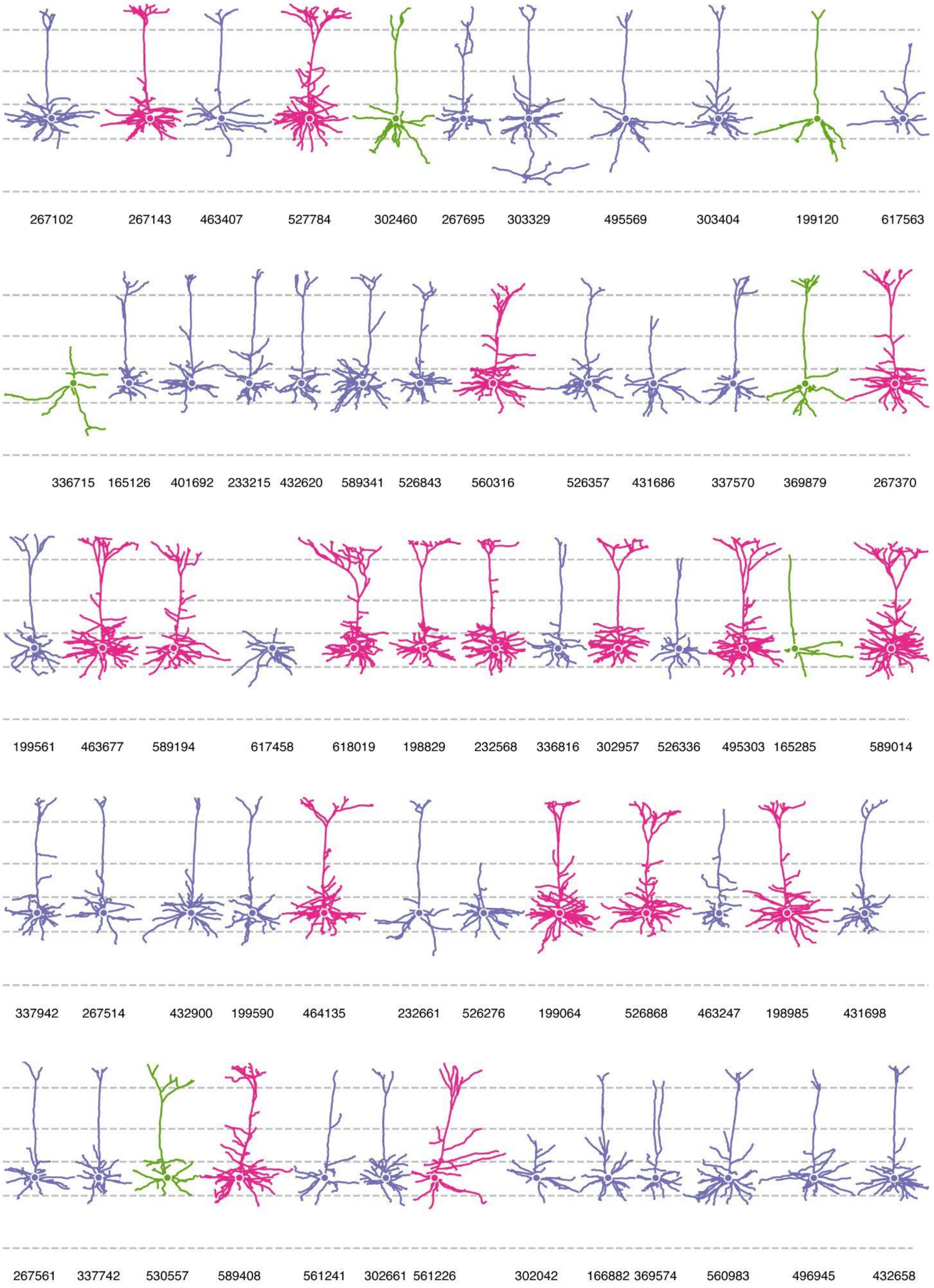

267102
267143
463407
527784
302460
267695
303329
495569
303404
199120
617563
336715
165126
401692
233215
432620
589341
526843
560316
526357
431686
337570
369879
267370
199561
463677
589194
617458
618019
198829
232568
336816
302957
526336
495303
165285
589014
337942
267514
432900
199590
464135
232661
526276
199064
526868
463247
198985
431698
267561
337742
530557
589408
561241
302661
561226
302042
166882
369574
560983
496945
432658

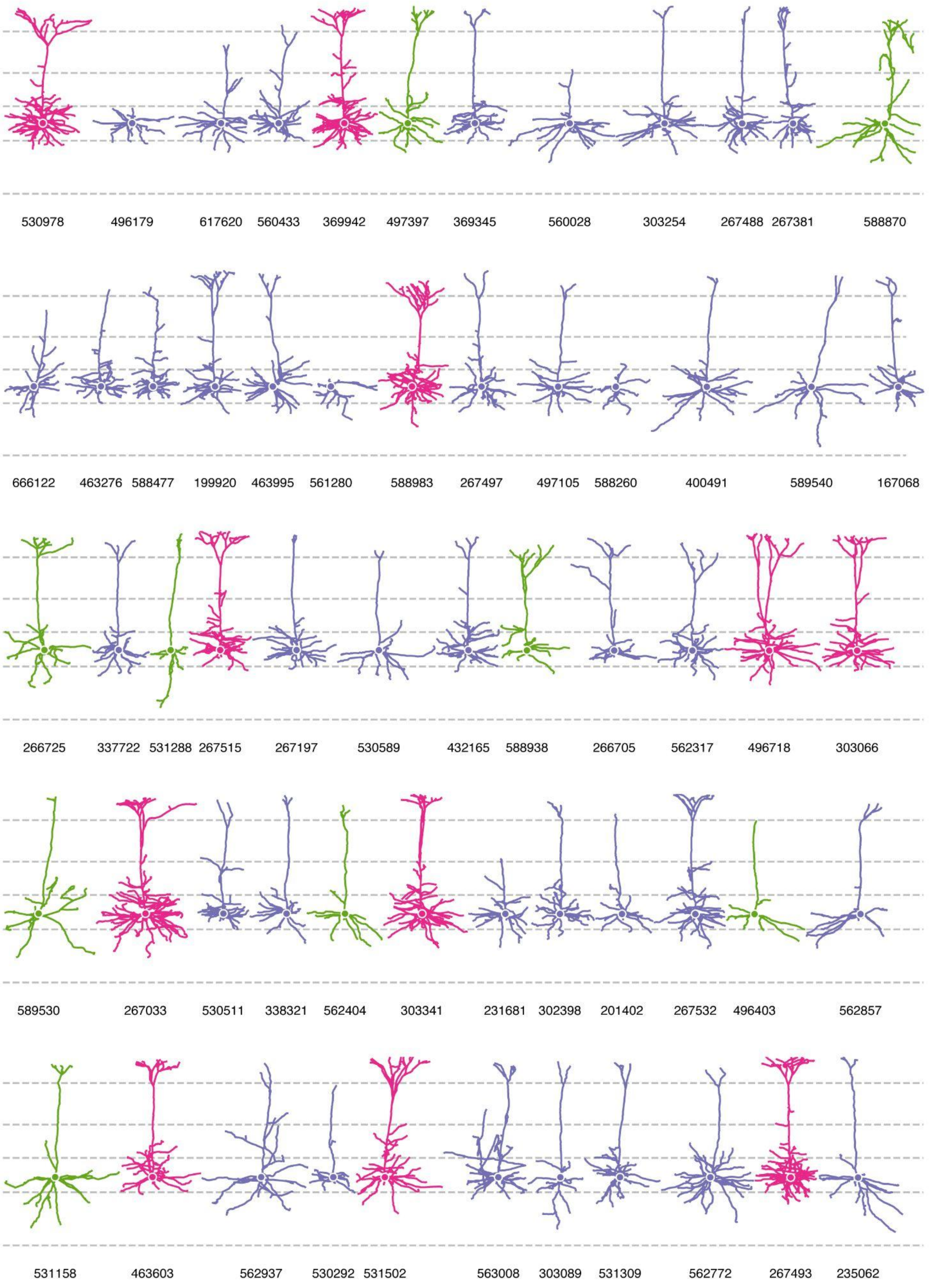

530978 496179 617620 560433 369942 497397 369345 560028 303254 267488 267381 588870
666122 463276 588477 199920 463995 561280 588983 267497 497105 588260 400491 589540 167068
266725 337722 531288 267515 267197 530589 432165 588938 266705 562317 496718 303066
589530 267033 530511 338321 562404 303341 231681 302398 201402 267532 496403 562857
531158 463603 562937 530292 531502 563008 303089 531309 562772 267493 235062

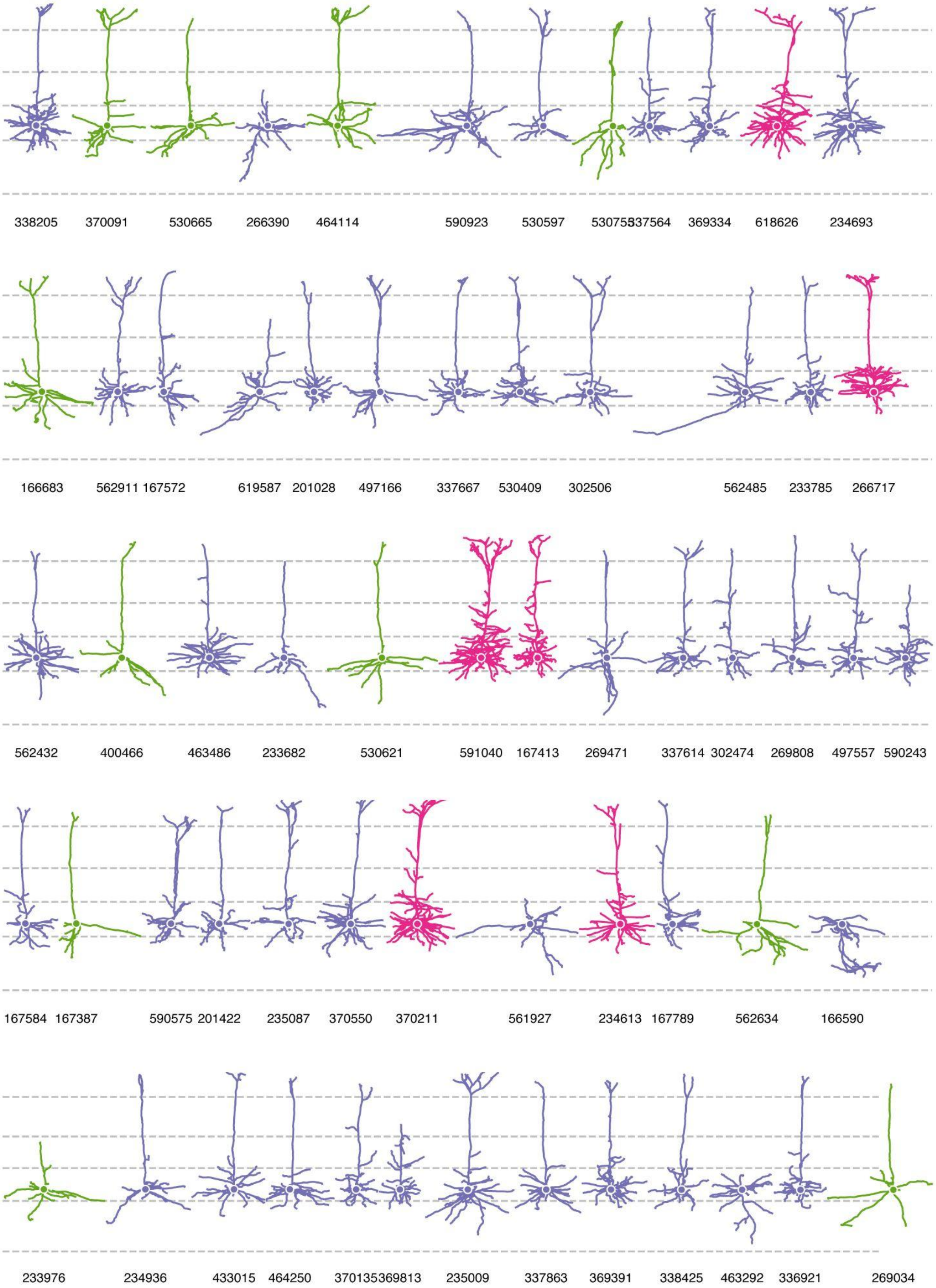
338205 370091 530665 266390 464114 590923 530597 53075337564 369334 618626 234693
166683 562911 167572 619587 201028 497166 337667 530409 302506 562485 233785 266717
562432 400466 463486 233682 530621 591040 167413 269471 337614 302474 269808 497557 590243
167584 167387 590575 201422 235087 370550 370211 561927 234613 167789 562634 166590
233976 234936 433015 464250 370135369813 235009 337863 369391 338425 463292 336921 269034

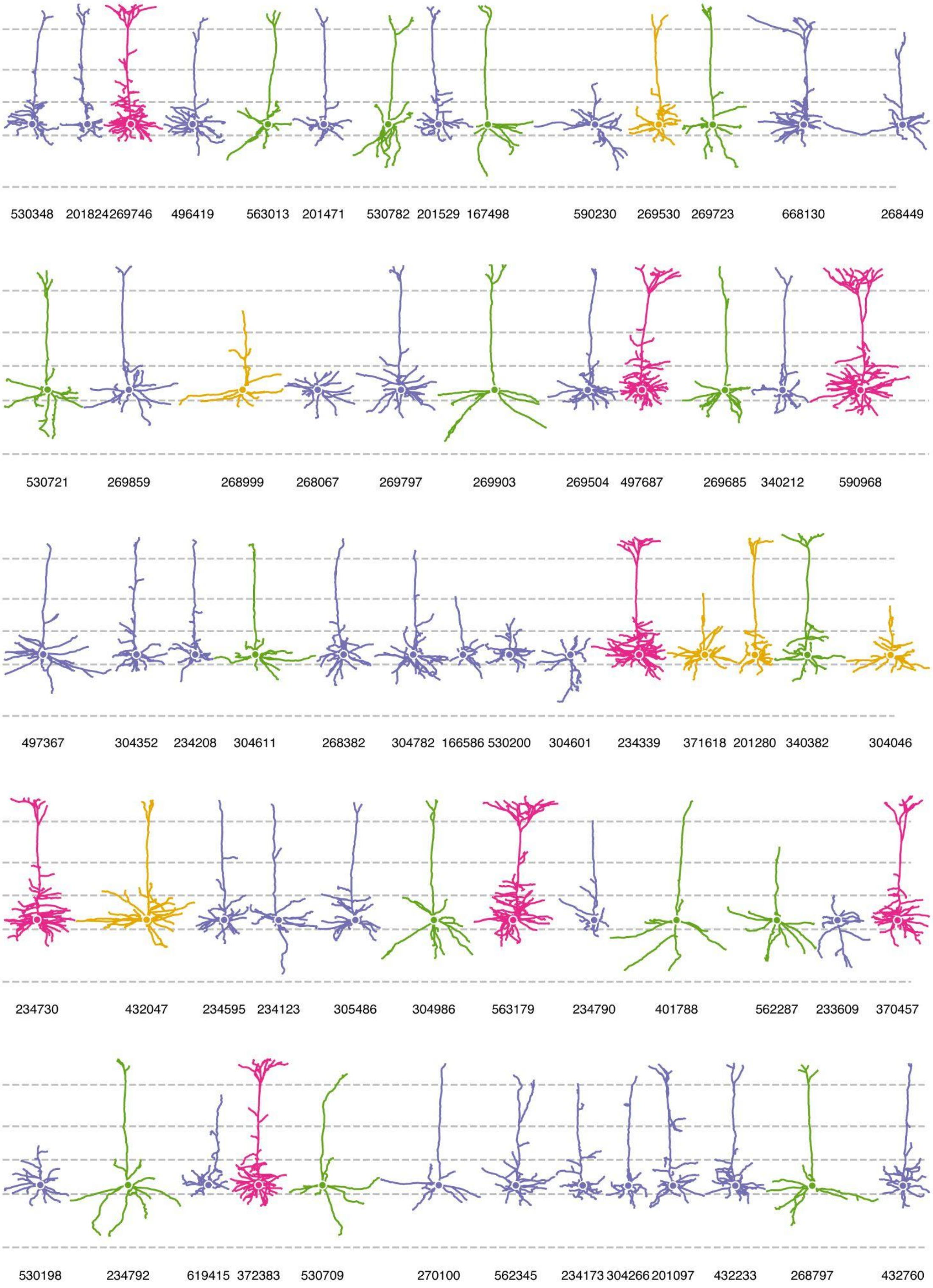
530348
201824
269746
496419
563013
201471
530782
201529
167498
590230
269530
269723
668130
268449
530721
269859
268999
268067
269797
269903
269504
497687
269685
340212
590968
497367
304352
234208
304611
268382
304782
166586
530200
304601
234339
371618
201280
340382
304046
234730
432047
234595
234123
305486
304986
563179
234790
401788
562287
233609
370457
530198
234792
619415
372383
530709
270100
562345
234173
304266
201097
432233
268797
432760

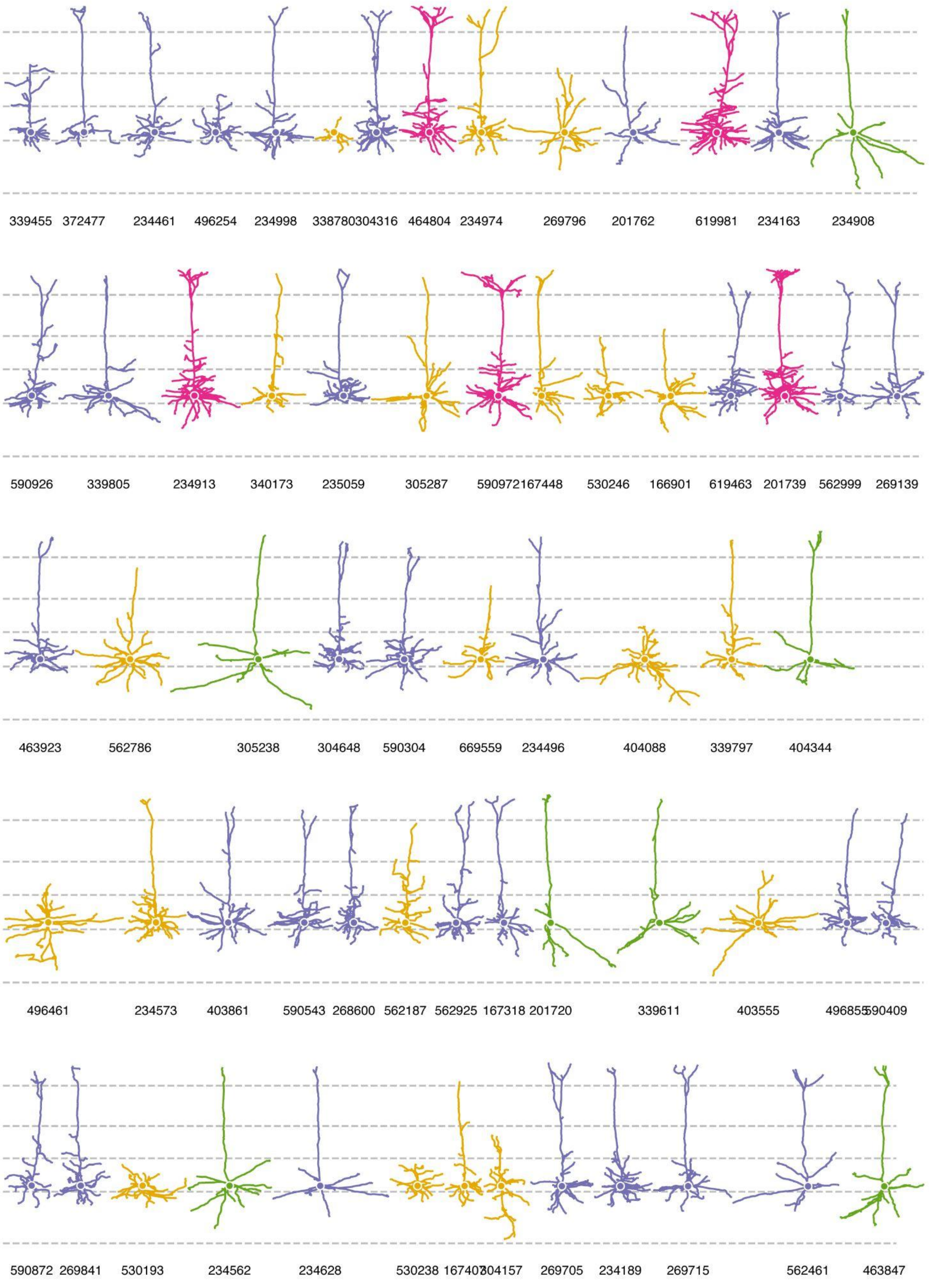
339455 372477 234461 496254 234998 338780304316 464804 234974 269796 201762 619981 234163 234908
590926 339805 234913 340173 235059 305287 590972167448 530246 166901 619463 201739 562999 269139
463923 562786 305238 304648 590304 669559 234496 404088 339797 404344
496461 234573 403861 590543 268600 562187 562925 167318 201720 339611 403555 496855590409
590872 269841 530193 234562 234628 530238 167407304157 269705 234189 269715 562461 463847

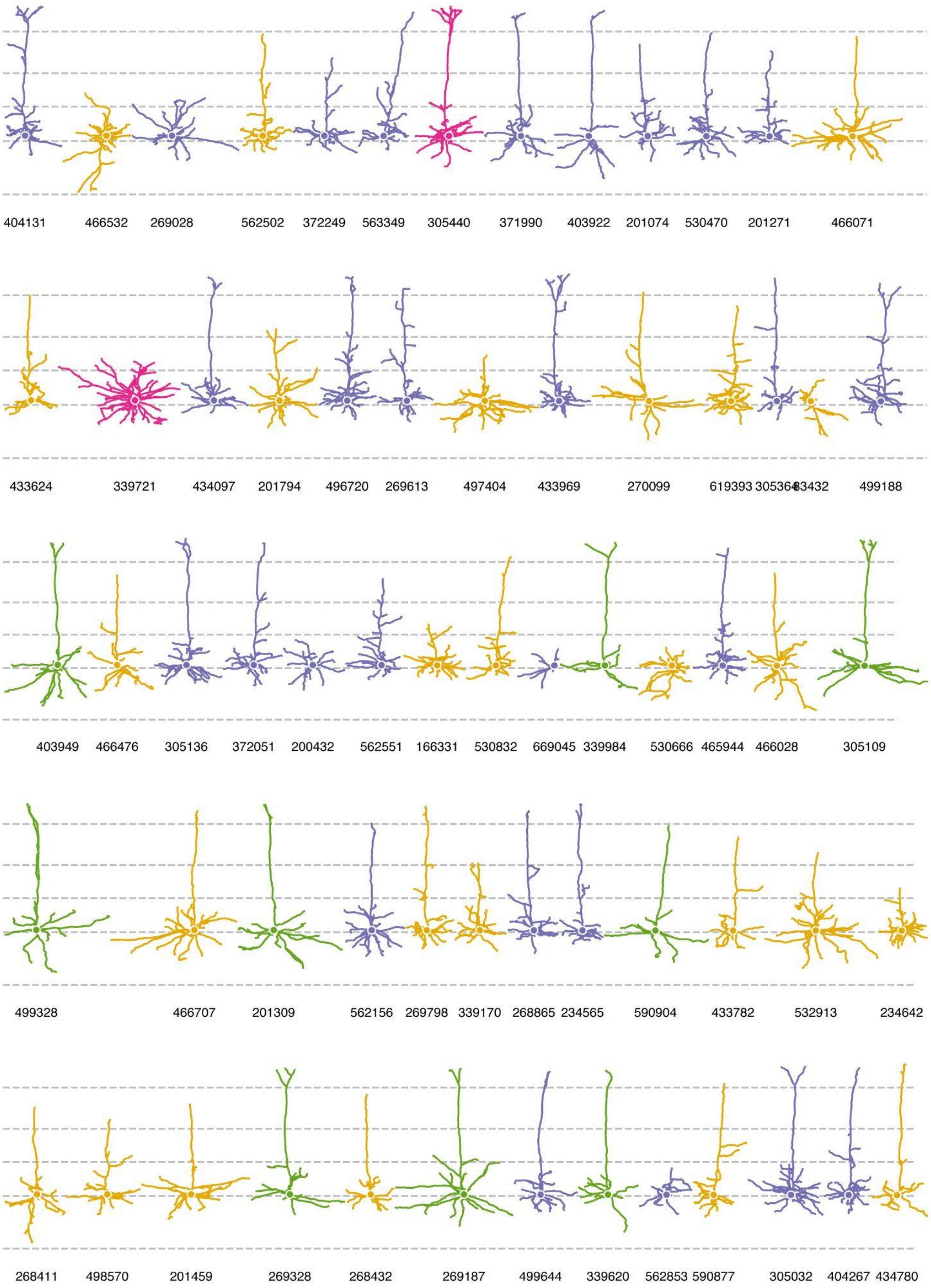
404131
466532
269028
562502
372249
563349
305440
371990
403922
201074
530470
201271
466071
433624
339721
434097
201794
496720
269613
497404
433969
270099
619393
305364
83432
499188
403949
466476
305136
372051
200432
562551
166331
530832
669045
339984
530666
465944
466028
305109
499328
466707
201309
562156
269798
339170
268865
234565
590904
433782
532913
234642
268411
498570
201459
269328
268432
269187
499644
339620
562853
590877
305032
404267
434780

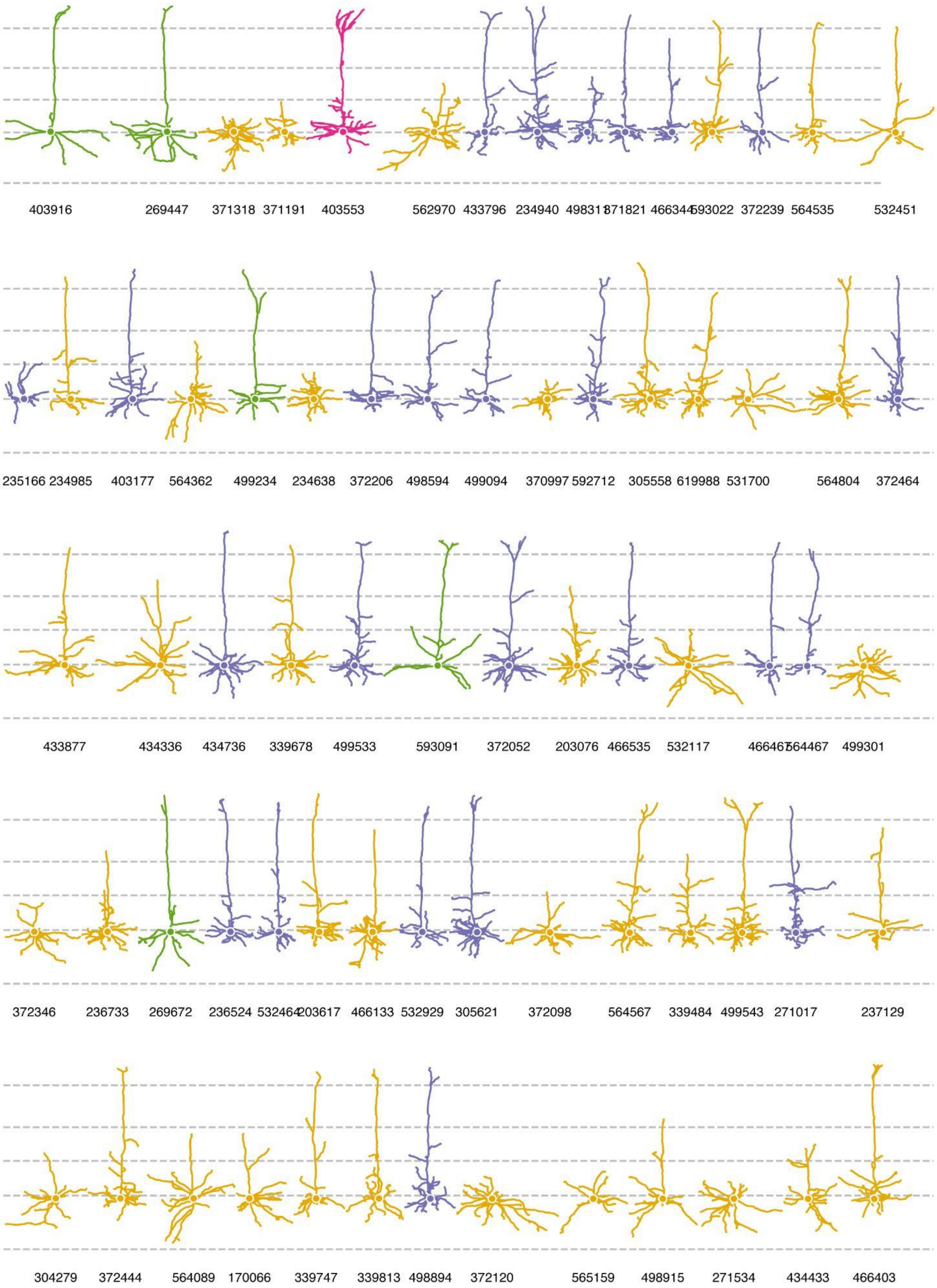
403916
269447
371318
371191
403553
562970
433796
234940
371821
593022
372239
564535
532451
235166
234985
403177
564362
499234
234638
372206
498594
499094
370997
592712
305558
619988
531700
564804
372464
433877
434336
434736
339678
499533
593091
372052
203076
466535
532117
564467
499301
372346
236733
269672
236524
203617
466133
532929
305621
372098
564567
339484
499543
271017
237129
304279
372444
564089
170066
339747
339813
498894
372120
565159
498915
271534
434433
466403

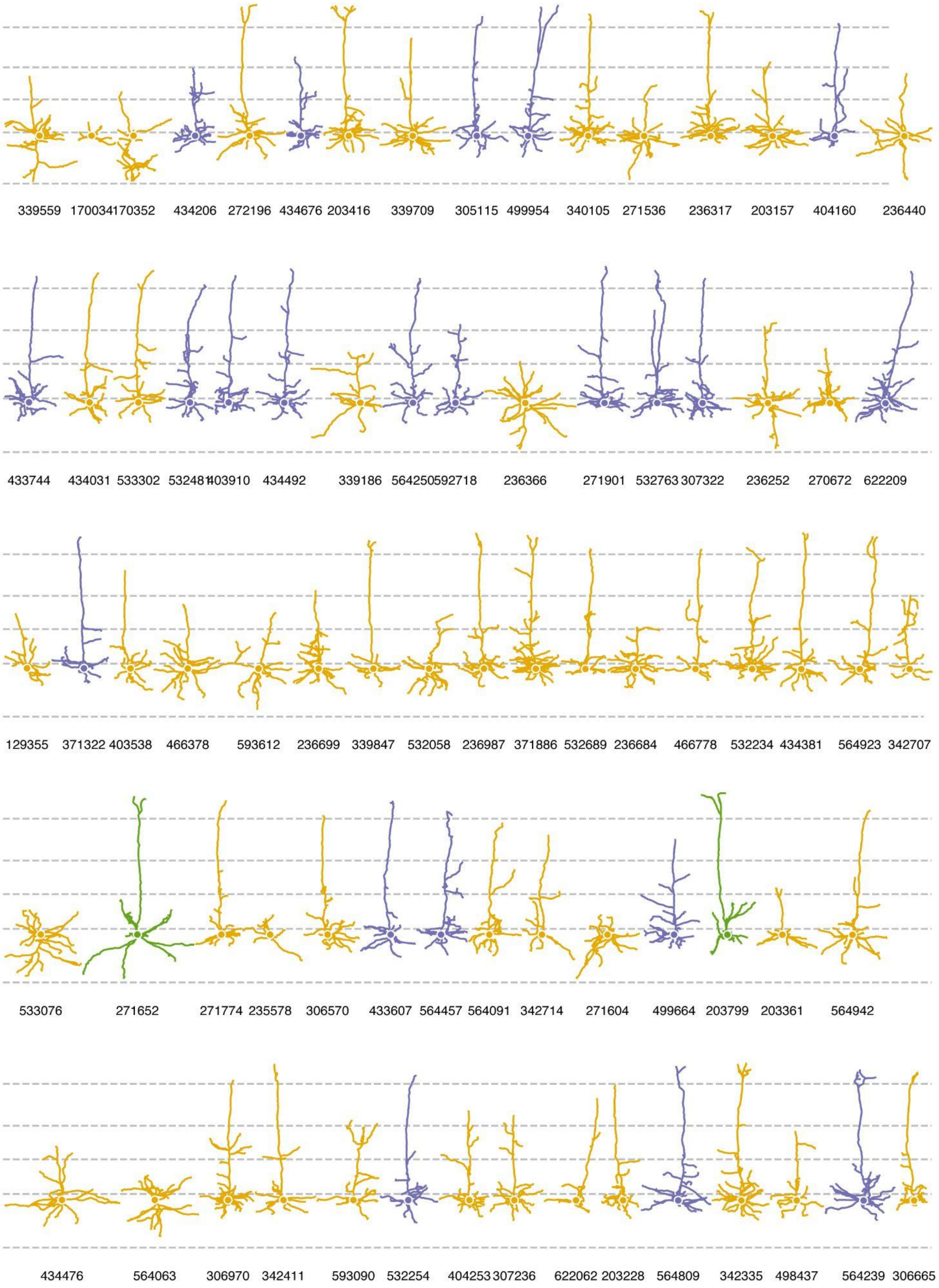
339559
170034
170352
434206
272196
434676
203416
339709
305115
499954
340105
271536
236317
203157
404160
236440
433744
434031
533302
53248
403910
434492
339186
564250
592718
236366
271901
532763
307322
236252
270672
622209
129355
371322
403538
466378
593612
236699
339847
532058
236987
371886
532689
236684
466778
532234
434381
564923
342707
533076
271652
271774
235578
306570
433607
564457
564091
342714
271604
499664
203799
203361
564942
434476
564063
306970
342411
593090
532254
404253
307236
622062
203228
564809
342335
498437
564239
306665

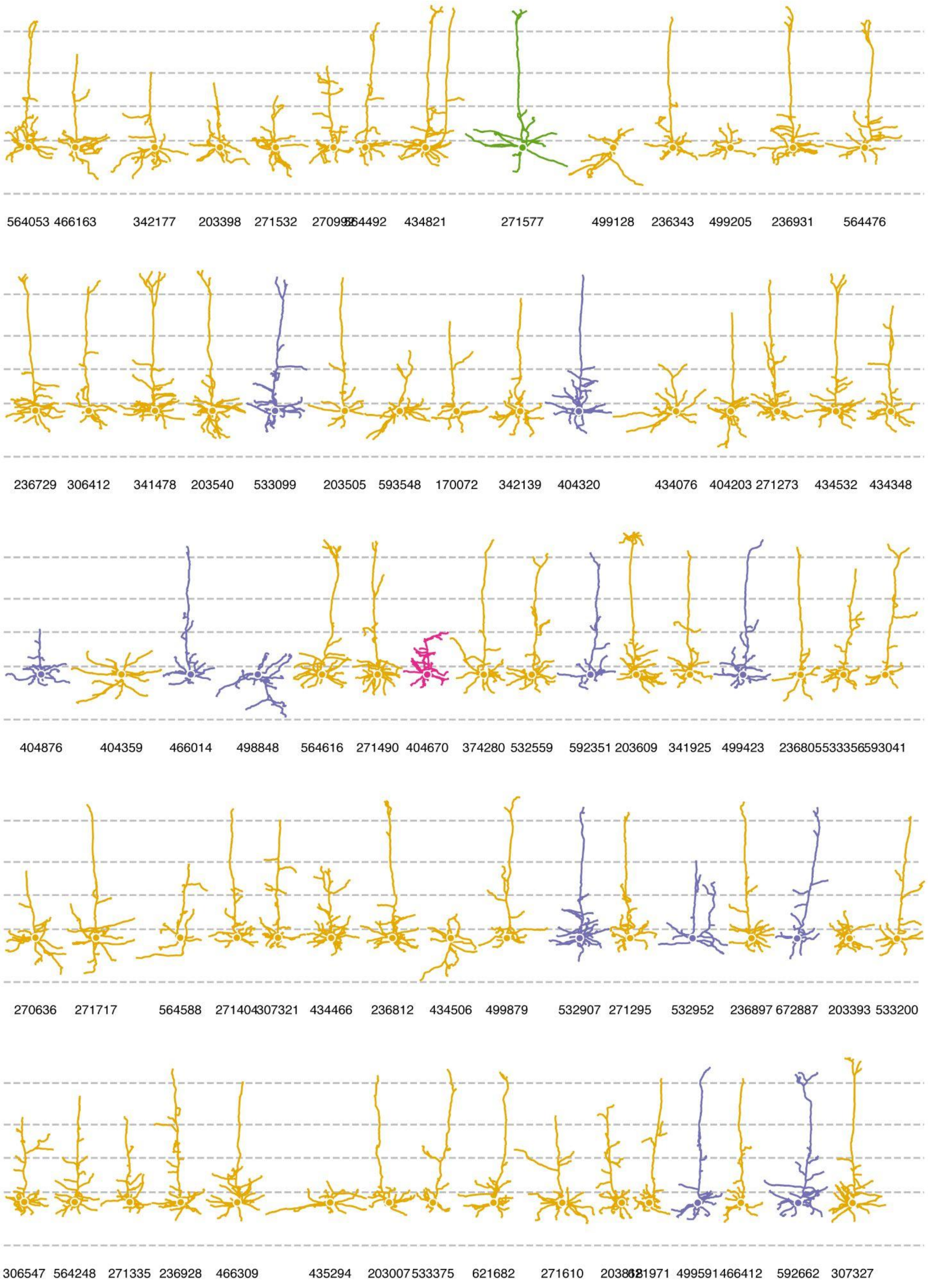
564053
466163
342177
203398
271532
564492
434821
271577
499128
236343
499205
236931
564476
236729
306412
341478
203540
533099
203505
593548
170072
342139
404320
434076
404203
271273
434532
434348
404876
404359
466014
498848
564616
271490
404670
374280
532559
592351
203609
341925
499423
236805533356593041
270636
271717
564588
271404307321
434466
236812
434506
499879
532907
271295
532952
236897
672887
203393
533200
306547
564248
271335
236928
466309
435294
203007533375
621682
271610
499591466412
592662
307327

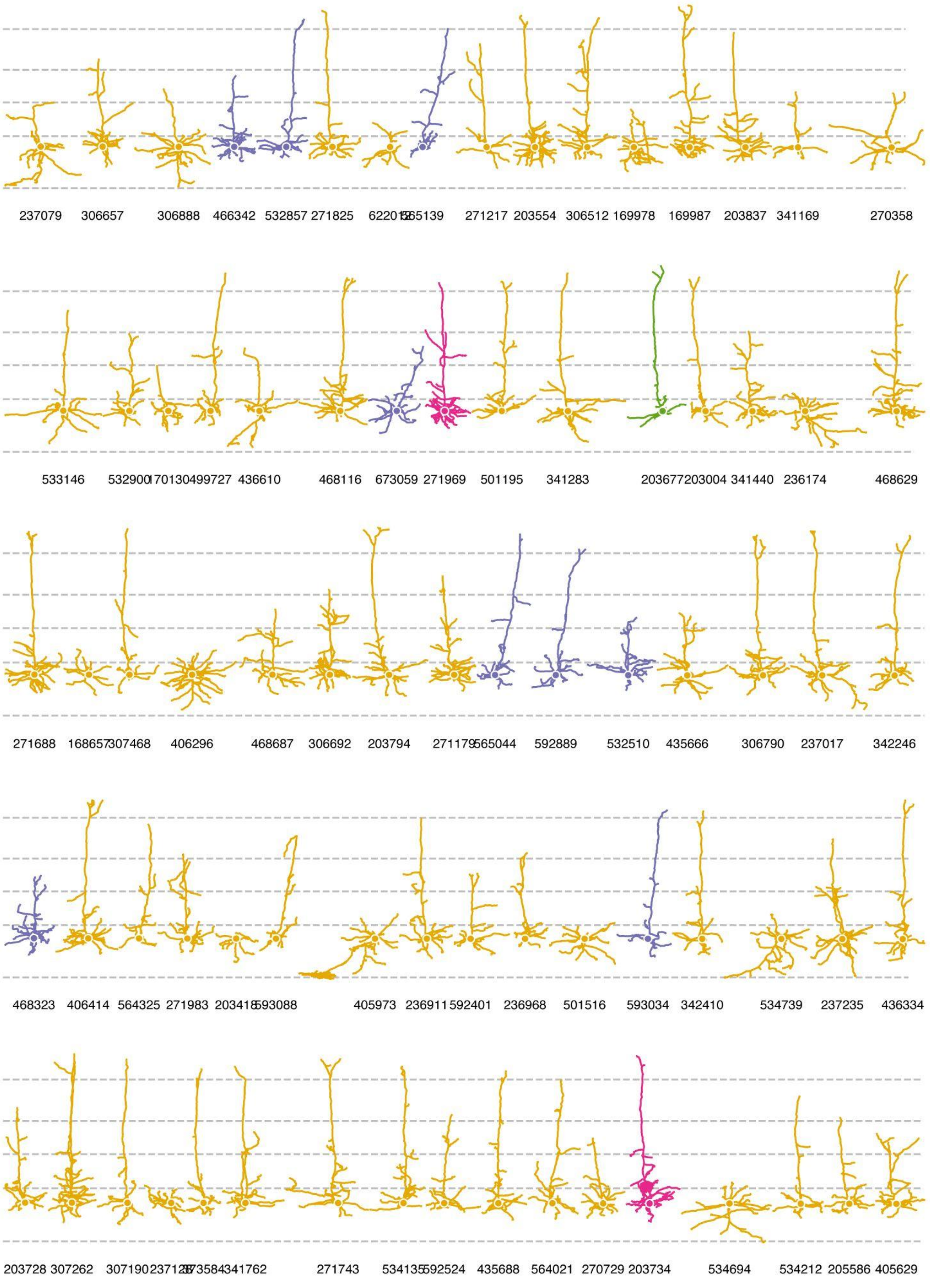
237079 306657 306888 466342 532857 271825 271217 203554 306512 169978 169987 203837 341169 270358
533146 532900 499727 436610 468116 673059 271969 501195 341283 203004 341440 236174 468629
271688 307468 406296 468687 306692 203794 592889 532510 435666 306790 237017 342246
468323 406414 564325 271983 405973 592401 236968 501516 593034 342410 534739 237235 436334
203728 307262 341762 271743 592524 435688 564021 270729 203734 534694 534212 205586 405629

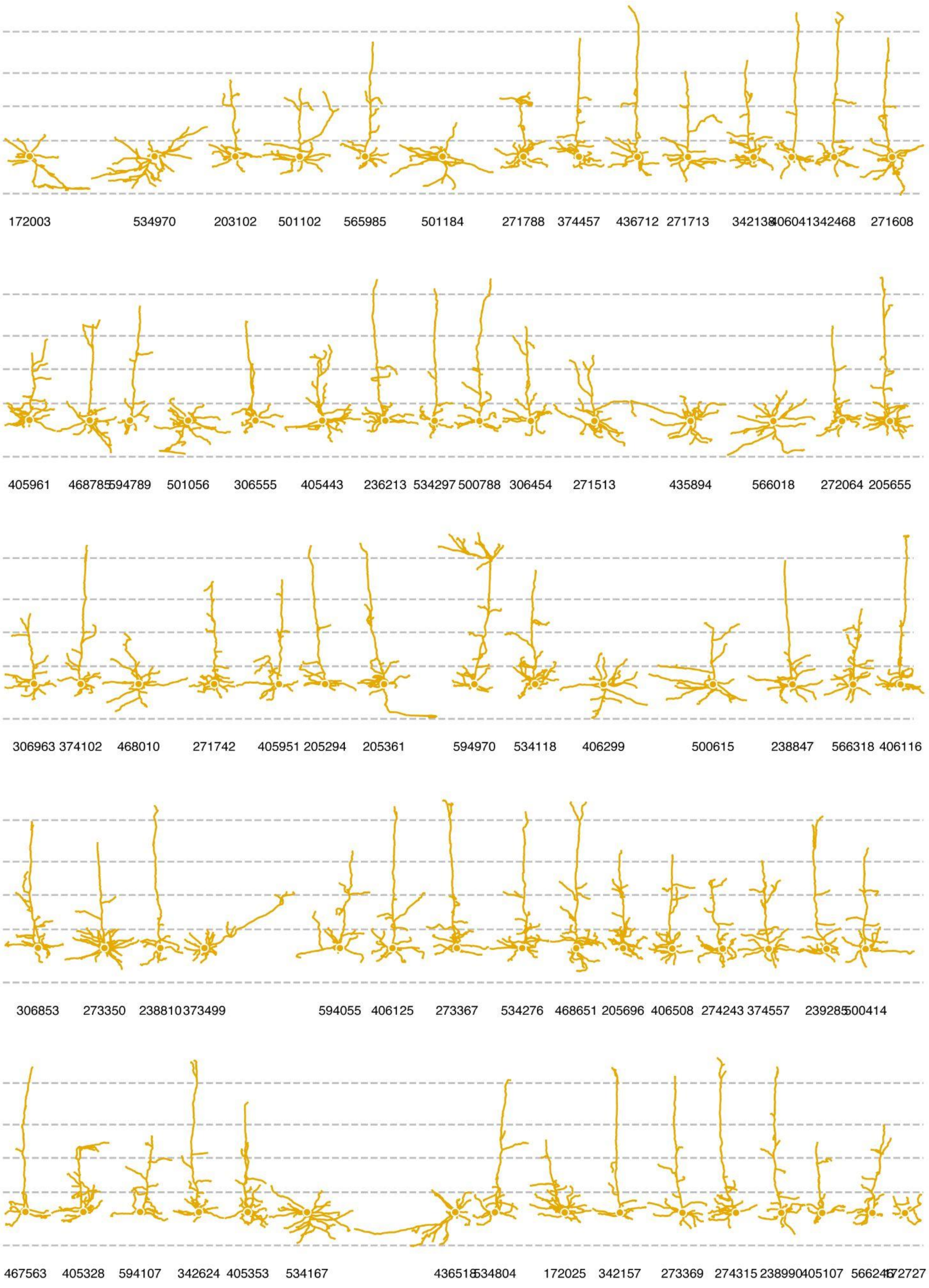
172003 534970 203102 501102 565985 501184 271788 374457 436712 271713 342138 406041 342468 271608
405961 468785 594789 501056 306555 405443 236213 534297 500788 306454 271513 435894 566018 272064 205655
306963 374102 468010 271742 405951 205294 205361 594970 534118 406299 500615 238847 566318 406116
306853 273350 238810 373499 594055 406125 273367 534276 468651 205696 406508 274243 374557 239285 500414
467563 405328 594107 342624 405353 534167 436518 534804 172025 342157 273369 274315 238990 405107 566246 172727

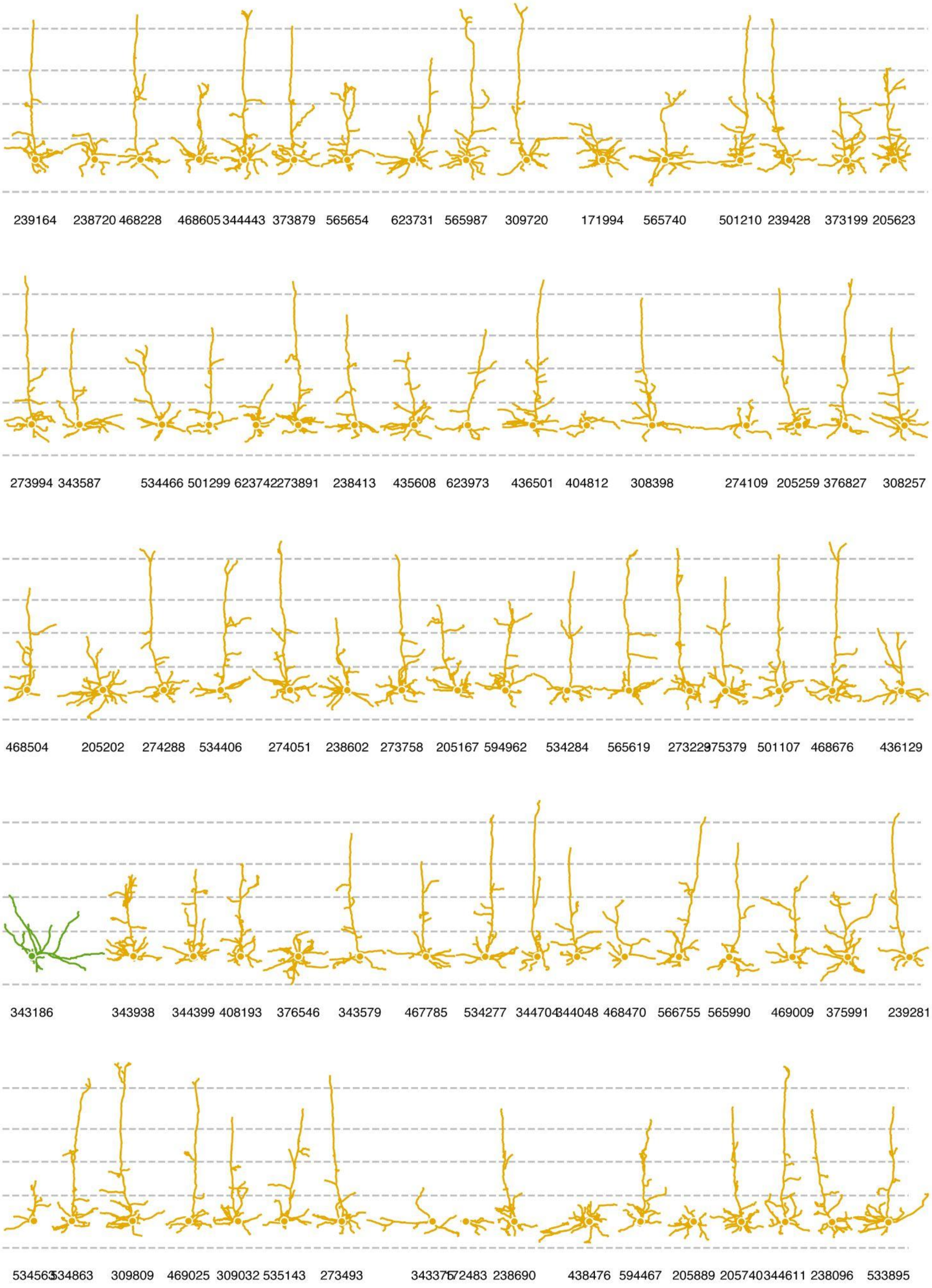
239164
238720
468228
468605
344443
373879
565654
623731
565987
309720
171994
565740
501210
239428
373199
205623
273994
343587
534466
501299
623742
273891
238413
435608
623973
436501
404812
308398
274109
205259
376827
308257
468504
205202
274288
534406
274051
238602
273758
205167
594962
534284
565619
27322975379
501107
468676
436129
343186
343938
344399
408193
376546
343579
467785
534277
344704344048
468470
566755
565990
469009
375991
239281
53456334863
309809
469025
309032
535143
273493
34337572483
238690
438476
594467
205889
205740
344611
238096
533895

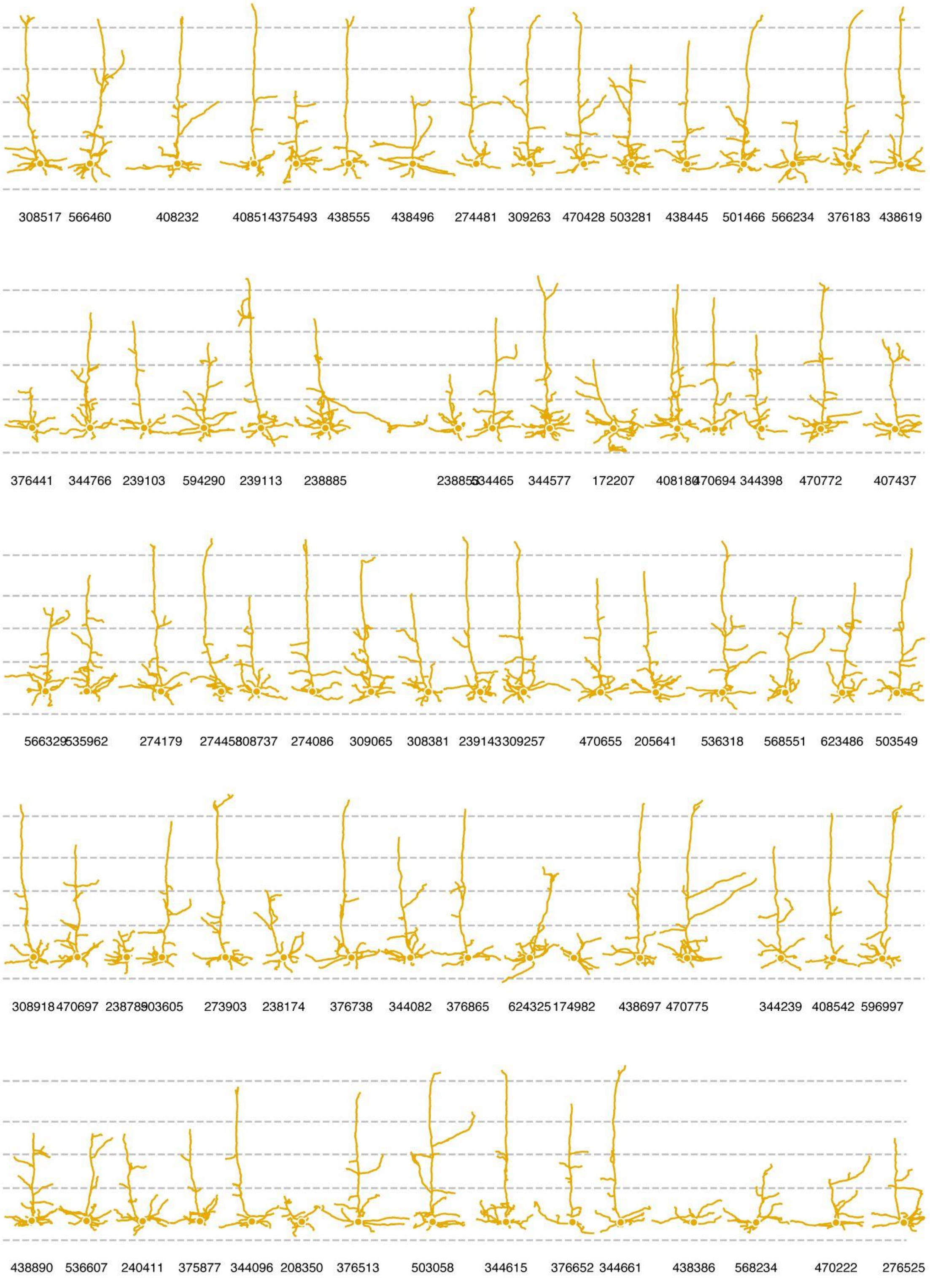
308517 566460 408232 408514375493 438555 438496 274481 309263 470428 503281 438445 501466 566234 376183 438619
376441 344766 239103 594290 239113 238885 238855344465 344577 172207 408180470694 344398 470772 407437
566329535962 274179 27445808737 274086 309065 308381 239143309257 470655 205641 536318 568551 623486 503549
308918 470697 23878903605 273903 238174 376738 344082 376865 624325 174982 438697 470775 344239 408542 596997
438890 536607 240411 375877 344096 208350 376513 503058 344615 376652 344661 438386 568234 470222 276525

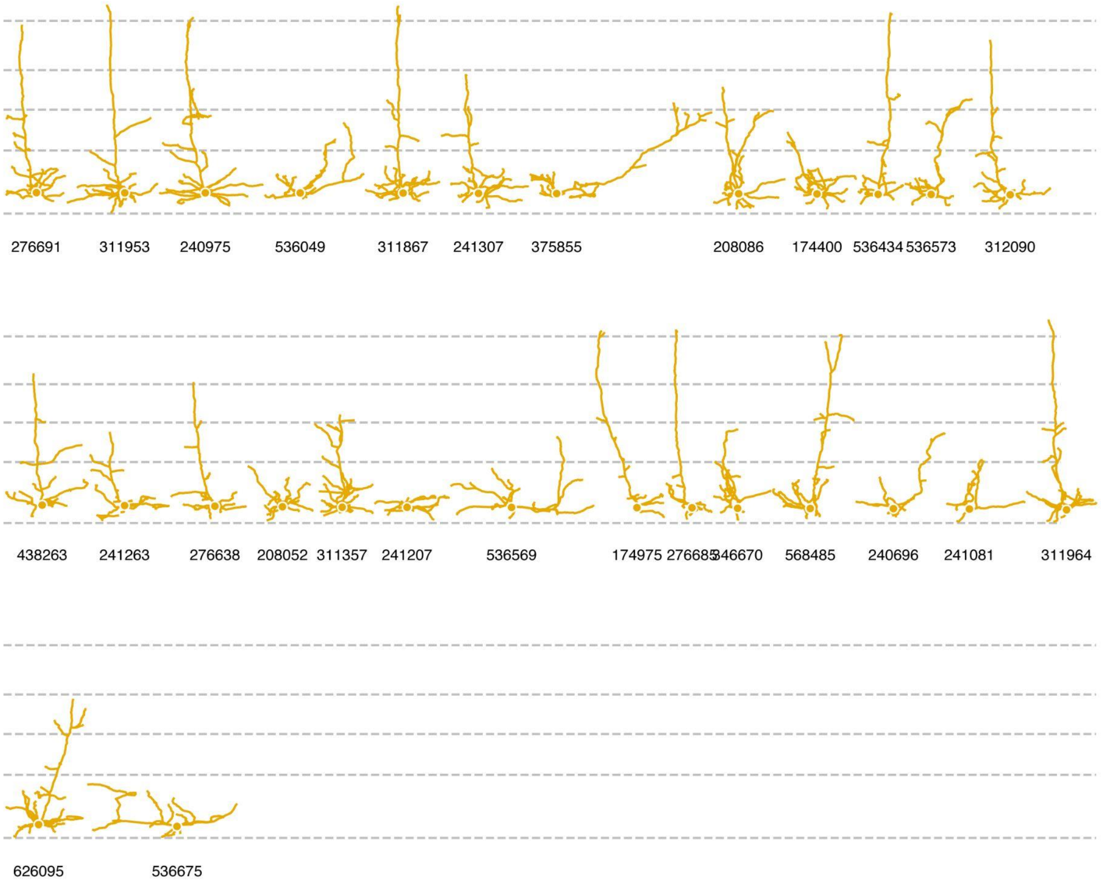
276691
311953
240975
536049
311867
241307
375855
208086
174400
536434
536573
312090
438263
241263
276638
208052
311357
241207
536569
174975
276685
346670
568485
240696
241081
311964
626095
536675
